# Dynamic engagement of dual-role regulators by the Sin3 complex

**DOI:** 10.64898/2026.02.26.708192

**Authors:** Julien Olivet, Neesha Rajesh Shewakramani, Joseph Cesare, Florent Laval, Jules Nde, Jasper Van de Veire, Yasmine Brammerloo, Bastien Brebel, Olivia Debnath, Aaron D. Richardson, Hong Yue, Yang Wang, Kerstin Spirohn-Fitzgerald, Irma Lemmens, Tong Hao, Jan Tavernier, Michael A. Calderwood, Hyuk-Soo Seo, Frederick P. Roth, Sirano Dhe-Paganon, Jean-Claude Twizere, David E. Hill, Marc Vidal, Michael P. Washburn, Kalyan Das

## Abstract

Gene expression is governed by dynamic switches between repressive and activating transcriptional states^1,2^. Among the molecules mediating these transitions, chromatin readers and transcription factors play pivotal roles^3,4^. However, how they assemble with regulatory machineries to enable crosstalk between gene repression and activation remains unknown. Here, we use an integrative structural dynamics approach – combining cryo-EM, crosslinking mass spectrometry, fragment-resolved protein interactome mapping and crystallography – to show how the dual-role chromatin reader Cti6 and transcription factors Ash1 and Ume6 engage the Sin3 deacetylase complex, a major regulatory hub in eukaryotes^5^. We find that Cti6 competes with Ash1 to drive its dynamic recruitment to a shared peripheral module, while Ume6 engages the Sin3 scaffold through a defined, minimal interface. Using high-throughput mutational scanning, we reveal deleterious and gain-of-function mutations in Sin3, identifying evolutionarily conserved residues essential for anchoring transcription factors. Together, these results provide structural and functional insights into how dual-role regulators engage the central Sin3 complex, revealing subtle assembly principles that may facilitate crosstalk between gene repression and activation. They also establish an integrative multidisciplinary framework to dissect the dynamics of macromolecular assemblies across biological systems.

## MAIN

Precise control of gene expression relies on the coordinated action of transcriptional activators and repressors that modulate chromatin structure^6^. In this context, histone acetylation and deacetylation provide central mechanisms to fine-tune gene accessibility and transcriptional outcomes^7,8^. These opposing activities are executed by large multisubunit assemblies that are co-recruited to specific loci by context-dependent chromatin readers and transcription factors^1,9,10^, enabling direct crosstalk between transcriptional repression and activation.

In budding yeast, the conserved, central Sin3 large (Sin3L) histone deacetylase (HDAC) complex functions predominantly as a transcriptional repressor and is homologous to the metazoan SIN3A complex^5,11,12^. Organized around the Sin3 scaffold and the Rpd3/HDAC catalytic subunit^13–16^, Sin3L comprises additional components that determine substrate recognition, genomic targeting and regulatory specificity^17–20^. Recent cryo-electron microscopy (cryo-EM) studies have revealed its overall horseshoe-shaped architecture^21,22^. However, substantial regions and subunits remain unresolved, including flexible scaffolding elements and dynamic modules that engage chromatin and activating machineries.

In this context, the chromatin reader Cti6 bridges the Cyc8-Tup1 corepressor to the SAGA acetyltransferase (HAT) complex while also constituting a stable Sin3L subunit^23–26^. Similarly, the GATA-like transcription factor Ash1 recruits Sin3L to the promoter of genes such as *HO*, but it also associates with SAGA through interactions with its deubiquitinase module^17,18,27,28^. Finally, although the Gal4-like transcription factor Ume6 recruits Sin3L to repress genes such as *CAR1*, *INO1* or *SPO13*, it also cooperates with the master meiotic regulator Ime1 in a Sin3/Rpd3-containing complex that colocalizes with activating SAGA components^17,18,26,29–32^. Thus, by operating at the interface of acetylation and deacetylation machineries, the Cti6, Ash1 and Ume6 subunits act as molecular hinges that can mediate transcriptional switches. However, the mechanisms underlying their dynamic engagement with the Sin3L complex remain unknown.

Here, to define how dual-function subunits engage the major transcriptional regulator Sin3L, we have developed an integrated structural dynamics strategy combining cryo-EM structures of native assemblies, crosslinking mass spectrometry (XL-MS), fragment-resolved protein interactome mapping, X-ray crystallography and data-driven modelling. This approach yields a fully resolved Sin3L core architecture and reveals that Cti6 forms a dynamic recruitment platform that directs Ash1 to a conserved peripheral module through competitive and repositioning mechanisms. Complementary fragment-resolved interactome mapping across ∼1.5x10^6^ pairwise combinations delineates subunit interactions and identifies direct contacts between Ume6 and core Sin3L components, enabling structural resolution of the Ume6-Sin3 interface that drives locus-specific recruitment. High-throughput mutational scanning further defines evolutionarily conserved determinants of Sin3L engagement, uncovering both loss- and gain-of-function variants at regulatory hotspots. Together, these findings establish a mechanistic framework for how dual-function chromatin readers and transcription factors dynamically engage Sin3L, revealing fundamental molecular principles that may coordinate acetylation and deacetylation machineries during transcriptional switching in eukaryotes.

### Core and fine-tuning subunits define locus-specific Sin3L regulation

To assess the contribution of each canonical Sin3L subunit to transcriptional regulation, we generated yeast strains lacking individual components and monitored derepression of the synthetic *SPAL10::URA3* reporter^33^. In this system, the *URS1* motif upstream of the *URA3* gene recruits Sin3L via Ume6^29,31^, enforcing transcriptional repression that prevents growth on uracil-deficient media while conferring resistance to 5-fluoroorotic acid (5FOA). Deletion of *UME6*, *SIN3*, *RPD3*, *SDS3*, *DEP1* or *SAP30* caused strong *SPAL10::URA3* derepression, producing growth phenotypes comparable to the positive control (**Fig. 1a**), thereby defining these factors as core Sin3L subunits required for repression at this locus. In contrast, *ash1Δ*, *cti6Δ* and *rxt3Δ* retained repression similar to wild-type and the *ura3Δ* negative control, indicating that these subunits are dispensable at *SPAL10::URA3*. However, mutants lacking *RXT2*, *PHO23* or *UME1* showed subtle derepression phenotypes, suggesting intermediate roles.

**Fig. 1.**
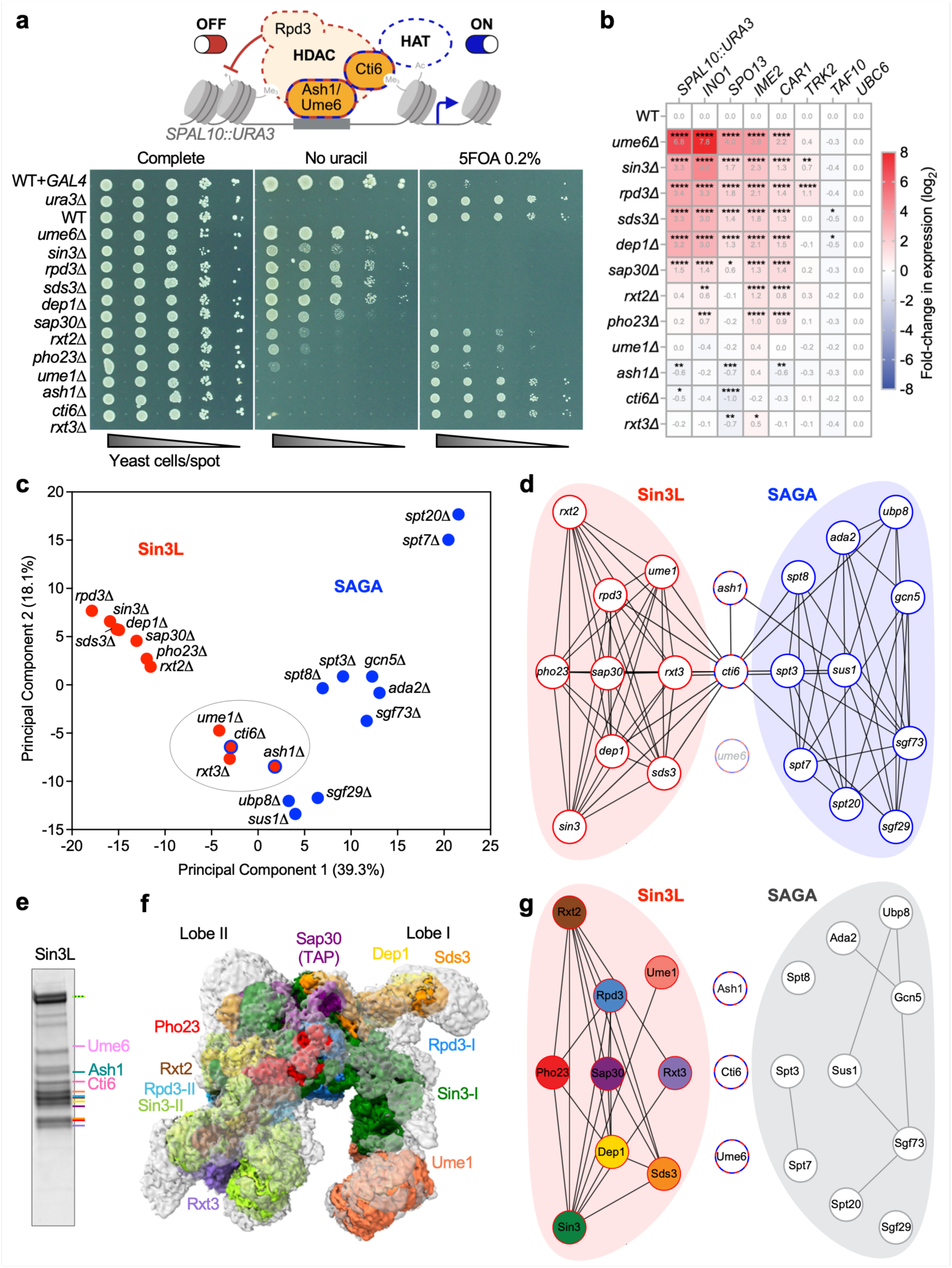
| Native Sin3L architecture displays a rigid catalytic core with flexible peripheries. **a**, Top, schematic of the dual roles of Cti6, Ash1 and Ume6 in transcriptional repression (Sin3L, red dashed line) and activation (HAT, blue dashed line). Me_3_ and Ac denote histone modifications; the + sign indicates lysine positive charge following deacetylation. Bottom, phenotypic profiling of Sin3L subunit deletion strains on SC, SC-U and 5FOA media. Each spot in the first column of the 10x dilution series contains ∼30,000 yeast cells. **b**, RT–qPCR analysis of Sin3L-regulated genes upon individual subunit deletions (n=3 biologically independent samples per group). Values represent mean log_2_ expression changes; *UBC6* was used for normalization and *TAF10* as a negative control. **c**, Principal component analysis of gene expression profiles from the Deleteome compendium^36^ for Sin3L (red) and SAGA (blue) deletions. The cluster containing *ume1Δ*, *rxt3Δ*, *cti6Δ* and *ash1Δ* is highlighted and the blue outlines indicate overlap with SAGA-associated signatures. **d**, Network representation of transcriptome similarities (false discovery rate <0.05; Pearson correlation) between Sin3L (red) and SAGA (blue) deletions. Nodes represent strains; edges denote similarity. The *ume6* node is faded due to missing data. Dual-function subunits are outlined by a dashed red/blue border. **e**, Coomassie-stained SDS-PAGE of Sap30-TAP–purified Sin3L complex. Canonical subunits are indicated by coloured lines as per their molecular weights. **f**, Unsharpened cryo-EM map of the Sap30-TAP-purified Sin3L complex (3.78 Å, contour level of 2σ), showing modelled regions in colour and unassigned, flexible densities in grey. **g**, Protein interaction network of the Sin3L core (red) and SAGA (grey) derived from structural contacts; nodes represent proteins and edges direct interactions based on the structure from (**f**) and published data for SAGA. For gel source data, see Supplementary Fig. 1.

To evaluate subunit contributions beyond the *SPAL10::URA3* reporter, we quantified the expression of five Sin3L target genes in each deletion strain. All genes except *TRK2* harbor a *URS1* motif recognized by Ume6 in their promoters, whereas *INO1* and *CAR1* additionally contain Ash1 binding sites^34^. Consistent with the *SPAL10::URA3* growth assay (**Fig. 1a**), all Sin3L-regulated loci were significantly derepressed in *ume6Δ*, *sin3Δ*, *rpd3Δ*, *sds3Δ*, *dep1Δ* and *sap30Δ* mutants (**Fig. 1b**), while *TRK2* was derepressed only in *sin3Δ* and *rpd3Δ*, as previously observed^35^. Deletions of *RXT2* and *PHO23* also derepressed *INO1*, *CAR1* and *IME2*. Strikingly, *ume6Δ* increased *INO1* expression by ∼250-fold, while *rpd3Δ* produced only a ∼10-fold effect, underscoring a predominant regulatory role for Ume6 at this locus. In contrast, deletions of *ASH1*, *CTI6* or *RXT3* reduced expression of *SPO13*, and in some cases *CAR1* and *SPAL10::URA3*, consistent with complementary functions in transcriptional activation^23,27^.

To map the genome-wide regulatory roles of Sin3L subunits, we analyzed transcriptional profiles from the Deleteome compendium^36^. To explore parallels with the SAGA complex, we also included transcriptomes from its subunit deletions. Hierarchical clustering of 21 transcriptomic signatures largely segregated Sin3L and SAGA components, except for one cluster that contained a mix of subunits including Rxt3, Ume1, Cti6 and Ash1, indicating their dual regulatory roles (**Extended Data Fig. 1**). Notably, deletion of *CTI6* or *ASH1* produced transcriptional responses globally opposite to those observed in *rpd3Δ* and other core subunit deletions, such that genes upregulated upon *RPD3* loss were downregulated or unchanged in *cti6Δ* and *ash1Δ*. Principal component analysis further confirmed the clustering by clearly separating core Sin3L and SAGA subunits, while positioning *rxt3Δ*, *ume1Δ*, *cti6Δ* and *ash1Δ* between the two distinct groups (**Fig. 1c**). Consistently, pairwise Pearson correlation networks recapitulated this pattern, highlighting the dual function of Cti6 and Ash1 as bridging factors between Sin3L and SAGA (**Fig. 1d**). These findings expose locus-specific assemblies within Sin3L, with gene-dependent core subunits driving repression and additional components fine-tuning regulation in a context-dependent manner.

### Native Sin3L architecture reveals a rigid core with flexible peripheries

To determine how the hierarchical organization of Sin3L components is reflected in three-dimensional space, we independently purified the native Sin3L complex using three distinct affinity tags, each fused to a different subunit – Sin3, Sap30, and Rxt3 – representing separate regulatory modules (**Fig. 1a-d**). Optimized endogenous tandem affinity purifications (TAP) with minimal sample handling preserved Sin3L integrity and stability, producing highly pure complexes (A_260_/A_280_ = 0.65 ± 0.10) with remarkable yields (28.6 ± 5.5 μg/L). SDS-PAGE profiles and mass spectrometry quantifications confirmed the presence of all expected components, with strong enrichment of the 12 canonical Sin3L subunits relative to the negative control (**Fig. 1e, Extended Data Fig. 2a and Supplementary Table 1**). Enzymatic assays also confirmed that the purified complexes exhibit robust HDAC activity (EC_50_ ∼0.4 nM) and are sensitive to Trichostatin A inhibition (IC_50_ ∼4.6 nM) (**Extended Data Fig. 2b,c**). Single-particle cryo-EM analyses of the independently purified and crosslinked complexes yielded three high-resolution maps at 3.37-3.61 Å (**Extended Data Fig. 2d-j and Extended Data Table 1**). The reconstructions were highly similar, with an average map-to-map correlation of 0.89. The atomic models built into the cryo-EM densities had an average molecular mass of ∼0.42 MDa, substantially lower than the ∼1 MDa mass of Sin3L in solution, as inferred from size-exclusion chromatography^17,21^, stoichiometry-based calculations, and values derived from the measured hydrodynamic radius of the particles. The structures recapitulated key architectural features such as the Rpd3-II active site blockade by Rxt2^21,22^, and revealed a conserved rigid core across purified complexes. However, the peripheral regions in the cryo-EM maps had relatively lower resolution, consistent with higher flexibility (**Extended Data Fig. 3a-c**).

To characterize the flexible regions beyond the core complex, we reprocessed the cryo-EM data to cover a larger volume, producing extended maps at 3.48-3.78 Å resolution that revealed additional, well-defined density patches (**Fig. 1f and Extended Data Fig. 4a,b**). These included a second copy of Ume1 (Ume1-II) positioned near Sin3-II and Rxt3^21,22^, as well as the previously unresolved C-terminal region of Sin3-I which appears to wrap around Ume1-I. Overall, the volumes of the extended maps indicate that the native complex has an average molecular mass of ∼1 MDa, representing an increase of ∼0.6 MDa relative to the initial atomic models and consistent with the Sin3L mass measured in solution. Together, these results indicate that the Sin3L architecture comprises a stable core flanked by flexible peripheral domains that may accommodate dynamic regulatory factors, including the dual-function proteins Cti6, Ash1 and Ume6, which were absent from previously reported Sin3L structures (**Fig. 1g**).

### Dynamic functional domains shape flexible Sin3L peripheries

To assign the additional flexible density patches, we developed a pipeline integrating cryo-EM maps, XL-MS data, HADDOCK^37,38^, and AlphaFold3^39^. Using Sap30-TAP as bait, the purified Sin3L complex was crosslinked with the mass spectrometry-cleavable reagent DSSO^40,41^ (**Extended Data Fig. 4c**), yielding a total of 395 crosslinks (**Fig. 2a,b**). A no-tag purified sample served as negative control, producing no crosslinks between Sin3L subunits and confirming tag-dependent enrichment, consistent with the purification profiles (**Fig. 2c and Extended Data Fig. 2a**). Crosslinks were mapped onto the Sin3L core cryo-EM structure (**Fig. 1f**) using a Cα-Cα distance restraint ≤37 Å^12,42^ (**Supplementary Table 2**), resulting in a 73.8% match rate. Several additional crosslinks, in particular between Ume1, Rxt3 and Sin3 (**Extended Data Fig. 4d**), could not be explained by the cryo-EM structure, and clearly defined the presence of a second Ume1 copy (Ume1-II) in lobe II of Sin3L, positioned closer to Rxt3 and Sin3-II and matching with the unassigned cryo-EM density (**Fig. 1f**).

**Fig. 2.**
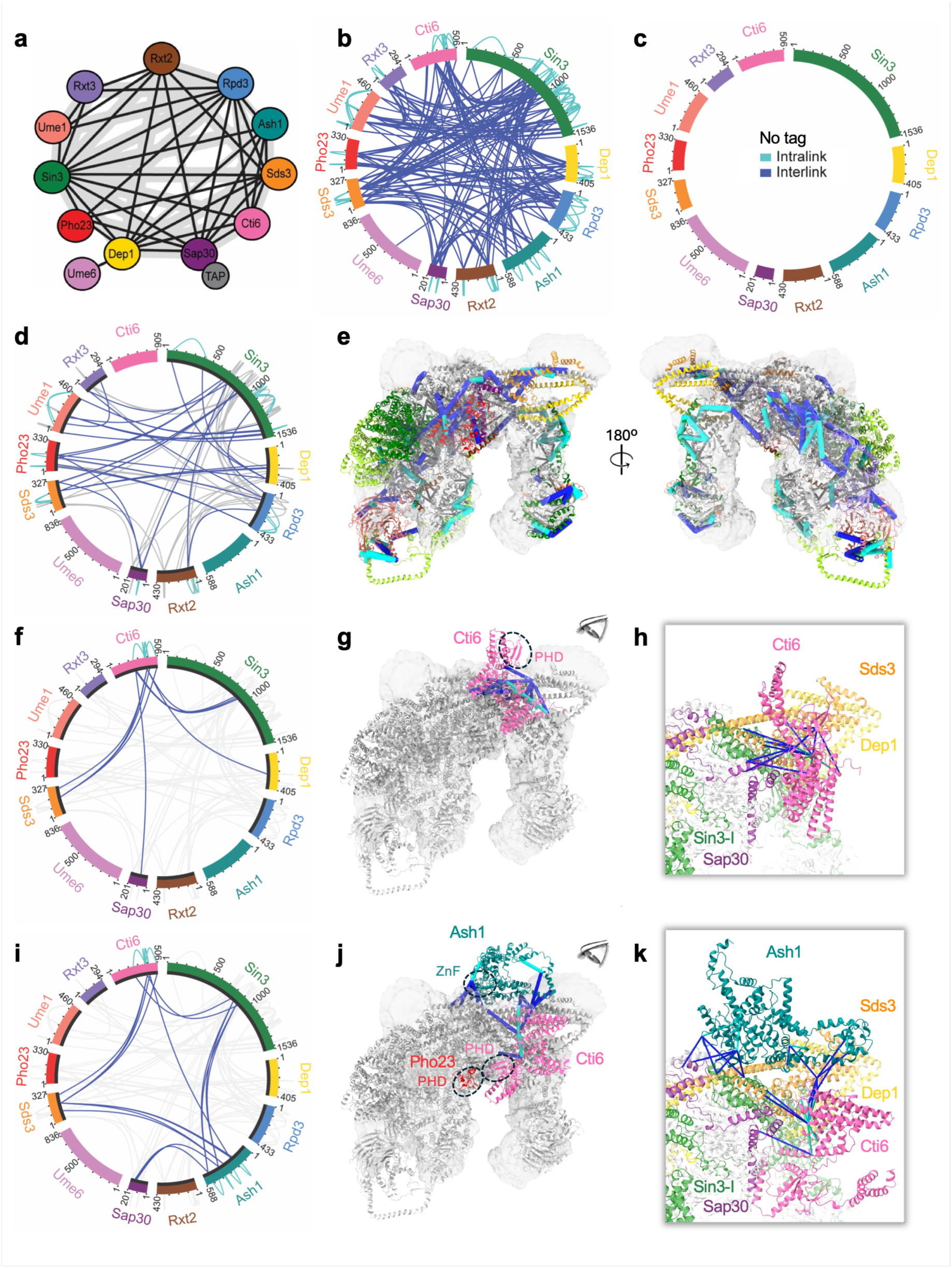
| Dynamic peripheral modules drive Ash1 recruitment to Sin3L. **a**, Schematic overview of crosslinks identified in the Sap30-TAP-purified Sin3L complex (n=3 biologically independent samples per group). **b**, Circular map of crosslinks from (**a**), with subunits represented as linear sequence bars; intra-and intersubunit links are shown in cyan and blue, respectively. **c**, No Sin3L inter- or intrasubunit crosslinks detected in the no-tag control, consistent with the SDS–PAGE profile (Extended Data Fig. 2a). **d**, Crosslinks consistent with the extended Sin3L model (Cα-Cα ≤37 Å, blue lines); crosslinks supporting the core structure are shown in light grey. **e**, Extended Sin3L model with newly built regions in colour overlaid on the core cryo-EM density (grey surface from Fig. 1f). Validated crosslinks from (**d**) are shown as solid lines. **f**, Subset of crosslinks (≤37 Å, blue lines) supporting incorporation of Cti6; background crosslinks from (**d**) are in light grey. **g**, Extended model including Cti6 (pink). Crosslinks from (**f**) are mapped as solid lines; the Cti6 PHD domain is circled. **h**, Zoomed view of Cti6 interactions with the Sin3L complex. Blue lines represent crosslinks. **i**, Crosslinks supporting positioning of Cti6 and Ash1 within the extended assembly (≤37 Å, blue lines). **j**, Model of Sin3L with Cti6 (pink) and Ash1 (teal). Crosslinks are shown as solid lines; the Cti6 and Pho23 PHD domains and the Ash1 zinc finger (ZnF) are circled. **k**, Zoomed view of the Cti6-Ash1 interactions with the Sin3L complex. The black bars within the protein bars of the circular plots (**d**,**f**,**i**) denote amino acid sequence coverage in the extended structural models of this study.

To model the architecture of individual multimeric sub-complexes, we employed AlphaFold3, retaining only models consistent with XL-MS distance restraints (Cα-Cα ≤37 Å), which streamlined rapid elimination of incompatible conformations and improved subunit interface predictions. Components selected from these assemblies were combined with distance restraints derived from the Sap30-TAP cryo-EM structure (**Fig. 1f and Extended Data Fig. 2h**) and XL-MS data in the HADDOCK software to assemble a complete Sin3L core including Ume1-II (**Fig. 2d,e**). The resulting model correlates with the Sap30-TAP (0.62), Sin3-TAP (0.58) and Rxt3-TAP (0.62) cryo-EM density maps (**Fig. 1f and Extended Data Fig. 4a,b**), and encompasses twelve fully-resolved subunits — Rpd3-I, Rpd3-II, Sin3-I, Sin3-II, Sds3, Dep1, Sap30, Rxt2, Pho23, Rxt3, Ume1-I and Ume1-II — for a total mass of 0.78 MDa, expanding the overall amino acid coverage from ∼50 to 100% (**Extended Data Fig. 4e**).

Importantly, the extended structure maintains the coiled-coil assemblies of Pho23-Rxt2 and Sds3-Dep1 observed in the earlier cryo-EM atomic models (**Extended Data Fig. 2g-i and Extended Data Fig. 4f,g**). However, the position of the Rxt2 loop blocking the Rpd3-II active site is slightly shifted in the extended structure (**Extended Data Fig. 4h**), reflecting a modest degree of conformational flexibility. In this state, the newly resolved Rxt2 C-terminal region partially blocks the catalytic site of Rpd3-I, suggesting that Rxt2 acts as a key modulator of Sin3L deacetylase activity (**Extended Data Fig. 4i**). The completed Rpd3-I and Rpd3-II models show that both C-terminal tails engage Pho23 and Sin3, potentially stabilizing Rpd3 dimerization (**Extended Data Fig. 4j**). Concurrently, the conserved Sin3 C-terminal regions (residues 1140-1412) interact with both Ume1 copies, and the expanded Sin3 N-termini encompass the paired amphipathic helix (PAH) 1 (residues 217-287) and PAH2 (residues 403-475) domains implicated in cofactor binding (**Extended Data Fig. 4k**). The model also resolves nucleosome-interacting elements, including the C-terminal plant homeodomain (PHD) of Pho23 (residues 272-329), oriented toward the central cavity between the two lobes. The remaining ∼0.2 MDa corresponding to the difference between the resolved 0.78 MDa and the estimated Sin3L mass in solution correspond to dynamic peripheral regions likely engaging transiently associated regulatory subunits, as described below.

### Cti6 drives Ash1 recruitment to Sin3L via a shared peripheral module

To characterize Sin3L peripheries in the extended structure, we examined the crosslinks involving additional subunits, including the transcription factors Ash1 and Ume6, as well as the chromatin reader Cti6. The presence of Ash1 and Cti6 was evident from the SDS-PAGE gels (**Fig. 1e and Extended Data Fig. 2a**), and numerous inter- and intra-subunit crosslinks involving Cti6, Ash1, and other Sin3L components were captured by XL-MS (**Fig. 2a,b**), enabling reliable modelling of their structural associations with the core complex.

Previous interaction network analyses showed that deletion of Cti6 results in loss of both Ash1 and Ume6, and identified Cti6, Sap30, Dep1, and Sds3 as components of the same Sin3L module^18^. Guided by these observations, we applied our AlphaFold3-HADDOCK pipeline to generate a complete Sin3L model in which Cti6 docking was restrained by XL-MS distances (Cα-Cα ≤37 Å) (**Fig. 2f**). The resulting 0.83 MDa assembly achieved a crosslinking match rate of 60.6% and recapitulated the previously defined Cti6-Sap30-Dep1-Sds3-Sin3 interaction module (**Fig. 2g,h**). In this model, Cti6 partially occupies the cryo-EM density, with its PHD finger (residues 63-153) oriented away from the core complex. The conserved basic region (CBR; residues 221-290), reported to bind the histone 2A-2B dimer^22^, forms a helix oriented perpendicular to the Dep1-Sds3 scaffold (**Fig. 2h**). The C-terminal domain (CTD), which contains a Sin3-interacting domain (SID; residues 426-499)^25^, folds into two helices that remain accessible to other subunits. Together, these features suggest that Cti6 adopts a flexible conformation that may be modulated by additional interacting proteins to facilitate nucleosome and transcription factor binding.

To further examine the flexibility of Cti6, we incorporated crosslinks involving Ash1 into our modelling pipeline. This approach enabled confident positioning of both Cti6 and Ash1 at the periphery of the extended Sin3L assembly, yielding a 0.90 MDa structure with a crosslinking agreement of 56.4% (**Fig. 2i**). Remarkably, Ash1 binding repositions Cti6 by rotating it clockwise around the Dep1-Sds3 scaffold to form a ring-like architecture (**Fig. 2j**). In this configuration, Ash1 engages not only Cti6 but also Sin3, Sap30, Dep1 and Sds3, and partially occupies the cryo-EM density previously assigned to Cti6 (**Fig. 2k and Supplementary Video 1**). The C-terminal GATA-like zinc finger (ZnF) domain of Ash1 (residues 493-560) resides above the Dep1-Sds3 scaffold and, owing to its flexibility, is poised to reposition for DNA engagement. In contrast to the Cti6-only assembly (**Fig. 2g**), the Cti6 PHD finger is relocated adjacent to the Pho23 PHD motif at the centre of the complex, with the CBR domain positioned immediately behind. This spatial convergence of Cti6 and Pho23 chromatin-reader domains upon Ash1 binding would support accommodation of at least one nucleosome between the two Sin3L lobes. Together, these integrative models reveal pronounced structural dynamics within the flexible peripheral regions of Sin3L and clarify the interconnected roles of the dual-function subunits Cti6 and Ash1. By demonstrating how dynamic peripheral modules couple regulatory factors to the core complex, the extended structures provide a framework for understanding how additional transient interactors, including Ume6, engage Sin3L to specific genomic loci.

### Ume6 engages Sin3L peripheries through Pho23 and Sin3 scaffold associations

While relatively abundant compared to other subunits, Ume6 was not readily apparent in the SDS-PAGE profiles of the purified Sin3L complexes (**Extended Data Fig. 2a and Supplementary Table 1**), and only a single crosslink between Ume6 and Dep1 was detected (**Fig. 2a,b**), limiting reliable docking based solely on XL-MS data. Thus, to gain further insights into the transient interactions of Ume6 with Sin3L, we conducted fragment-resolved protein interactome mapping *in cellulo* across all 12 canonical subunits. Using a high-precision yeast two-hybrid (Y2H) platform followed by validations with the mammalian kinase substrate sensor (KISS) assay^43^, we tested ∼1.5x10^6^ pairwise combinations between 1,020 unique fragments (**Extended Data Fig. 5**). This yielded multiple fragment-fragment interactions (FFIs) among nine Sin3L subunits, from which 11 minimal contiguous interfaces were delineated (**Fig. 3a and Extended Data Fig. 6a,b**). Importantly, 73% of these interactions (8/11) were validated using full-length subunits in the KISS assay at a 0% false discovery rate^43^ (**Extended Data Fig. 6c**). Two interfaces involving Ume6 (Ume6-Pho23 and Ume6-Sin3) were identified, and all other FFIs, including Sin3-Ume1 on lobe II, could be mapped onto the extended Sin3L structure, validating the overall architecture (**Fig. 3b**).

**Fig. 3.**
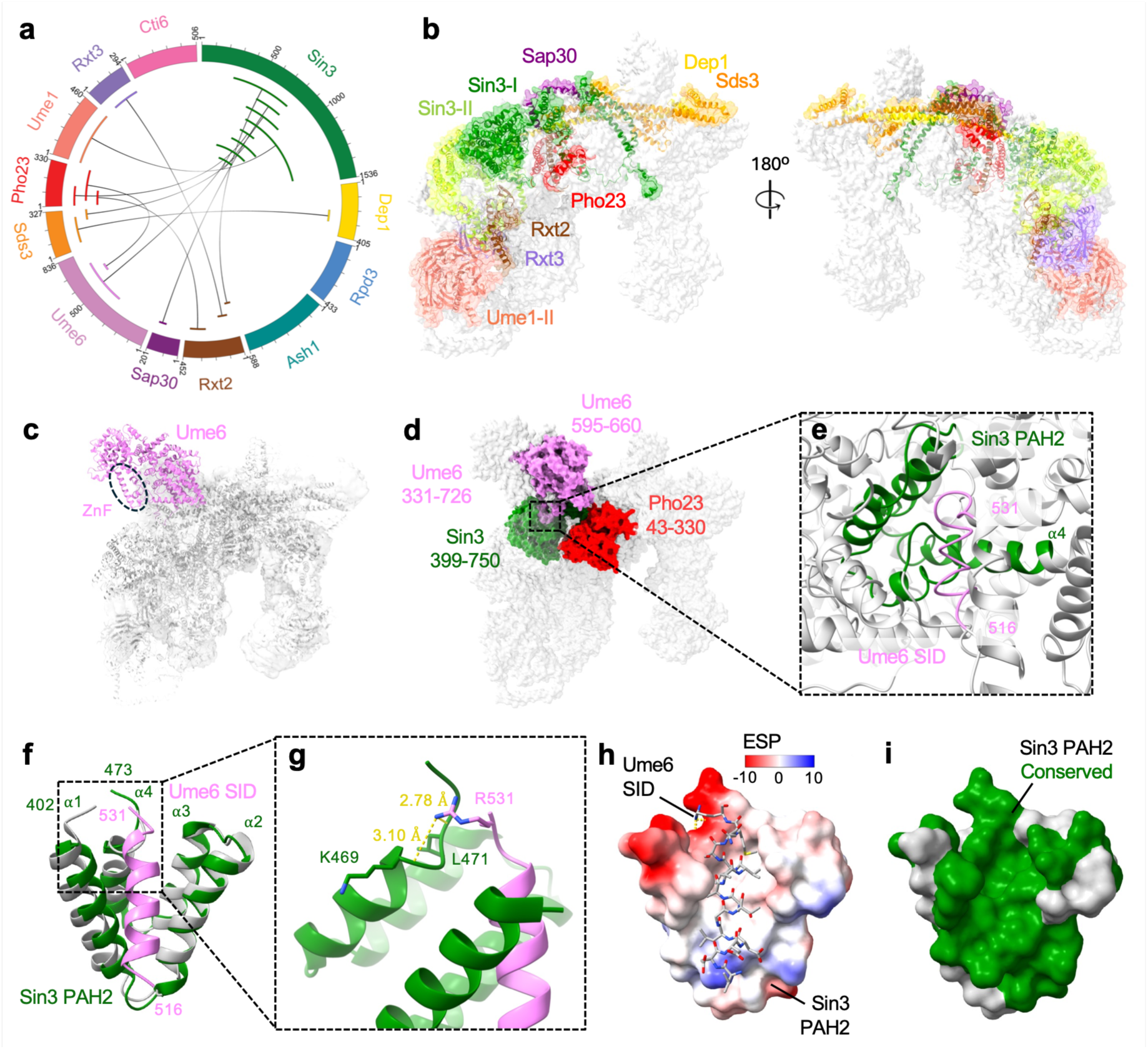
| Ume6 engages Sin3L peripheries through Pho23 and Sin3 scaffold associations. **a**, Circular summary map of minimal fragment–fragment interactions (FFIs) across the canonical Sin3L subunits. **b**, Minimal FFIs mapped onto the extended Sin3L structure containing Cti6. **c**, Model incorporating Ume6 (violet) into the extended Sin3L assembly; the Ume6 zinc finger (ZnF) is circled. **d**, Minimal Pho23-Ume6 and Sin3-Ume6 interactions mapped onto the structure, with interacting fragments indicated. **e**, Zoomed view of the Sin3 PAH2 domain engaging the Ume6 Sin3-interacting domain (SID). **f**, Crystal structure of Sin3 PAH2-Ume6 SID (PDB: 6XAW, in green) superimposed with apo Sin3 PAH2 (PDB: 6XDJ, in grey). **g**, Hydrogen bonds between Sin3 PAH2 and Ume6 SID residues. **h**, Electrostatic surface potential (ESP) of Sin3 PAH2 with Ume6 SID bound. **i**, Conserved surface residues between yeast Sin3 PAH2 and human SIN3A PAH2 highlighted in green.

To reveal how Ume6 engages the Sin3L complex, we modelled the two FFIs on the extended structure (**Fig. 2g**) using the AlphaFold3-HADDOCK pipeline applied to elucidate the dynamic interplay between Cti6 and Ash1. As an additional constraint, we included the crystal structure of the Sin3 PAH2-Ume6 SID interface (PDB: 6XAW) in our HADDOCK docking. The resulting integrated model with a molecular mass of 0.92 MDa preserved the Ume6-Sin3 interaction and positioned Ume6 in contact with Sin3-I, in the vicinity of an unassigned cryo-EM density (**Fig. 1f, Extended Data Fig. 4a,b and Fig. 3c**). However, Ume6 was located near Pho23 without forming direct contacts (**Fig. 3d**), and in proximity to the Dep1 C-terminal region, with which we detected a crosslink (**Fig. 2a,b**). Notably, the limited availability of structural information and intra- and inter-crosslinks involving Ume6 made its AlphaFold3-predicted folding difficult to interpret beyond its two established functional modules, i.e. the Sin3-interacting domain (SID) (residues 516-531) and the zinc finger (residues 765-809). In this conformation, the Ume6 C-terminus, encompassing the zinc finger domain, remained exposed and flexible (**Fig. 3c**), likely allowing DNA engagement in a cellular context. In the integrated model, the Ume6 SID-Sin3 PAH2 interface was characterized by the SID domain interacting mainly with helix α4 of PAH2 (**Fig. 3e**).

In contrast, the 1.8 Å resolution crystal structure of the Ume6 SID-Sin3 PAH2 domains revealed a helical architecture similar to mammalian orthologs^44,45^. In this structure, the PAH2 domain adopts an antiparallel four-helix bundle, with helices α1 and α2 forming a deep hydrophobic pocket that accommodates the amphipathic Ume6 α helix peptide (**Fig. 3f-h**). With a root mean square deviation (RMSD) of 1.0 Å between backbone atoms, the free (PDB: 6XDJ) and complexed PAH2 structures did not reveal significant conformational changes (**Fig. 3f and Extended Data Table 2**). In addition to the hydrophobic interactions, Ume6 Arg531 forms two hydrogen bonds with Sin3 Lys469 and Leu471 main-chain atoms to stabilize the Sin3-Ume6 interface (**Fig. 3f,g**). Interestingly, the hydrophobic pocket of the Sin3 PAH2 domain is flanked by highly charged residues at both ends, complementing the opposing charges at the termini of the Ume6 SID (**Fig. 3h**). To further illustrate the conservation between yeast and human SIN3 PAH2 domains, we mapped residues on the yeast PAH2 structure that are identical in human (**Fig. 3i**). This analysis revealed that both the hydrophobic pocket and the charged residues contacting the SID domain are evolutionarily conserved and therefore essential for transcription factor engagement with Sin3L in eukaryotes.

### Mutational scanning defines residues essential for Ume6 engagement

To define residues critical for the conserved Sin3 scaffolding interface mediating transcription factor binding, we performed mutational scanning on the Sin3-Ume6 interaction (**Extended Data Fig. 7a**). A Sin3 fragment spanning residues 272-885, encompassing the PAH2 (403-475) and PAH3 (656-727) domains that mediate Ume6 and Sap30 or Rxt2 binding, respectively, as well as the HDAC-interacting domain (HID; 747-848), was mutagenized by error-prone PCR, achieving ∼33% coverage of all possible amino acid substitutions (**Extended Data Fig. 7b,c**). Y2H selection against a Ume6 fragment (residues 265-726), followed by comparison of Z-scores derived from pre- and post-selection frequencies, enabled classification of Sin3 variants as deleterious, neutral, or enriched relative to wild-type.

In total, 4,036 mutations were evaluated. Most substitutions (3,041; ∼75%) were neutral (-1.96 < Z-score < 1.96), indicating broad mutational tolerance across the scanned region. By contrast, 995 alleles significantly affected the Sin3-Ume6 interaction (**Fig. 4a**). Of these, 582 were deleterious (Z-score < -1.96), including 508 missense and 74 nonsense mutations that disrupted 339 and 74 distinct residues, respectively, thereby defining critical contact determinants in the Sin3-Ume6 interface. Remarkably, the remaining 413 alleles were significantly enriched (Z-score > 1.96), including 353 missense and 60 nonsense substitutions, suggesting that specific changes confer a selective advantage (gain-of-function) and enhance binding relative to wild-type. Mapping of the mutational tolerance across the Sin3 fragment revealed a sharply localized cluster of deleterious substitutions between residues ∼420-475 (**Fig. 4a,b**), matching precisely with the PAH2 domain and thus validating the biological relevance of the Sin3 PAH2-Ume6 SID structure (**Fig. 3f-h**).

**Fig. 4.**
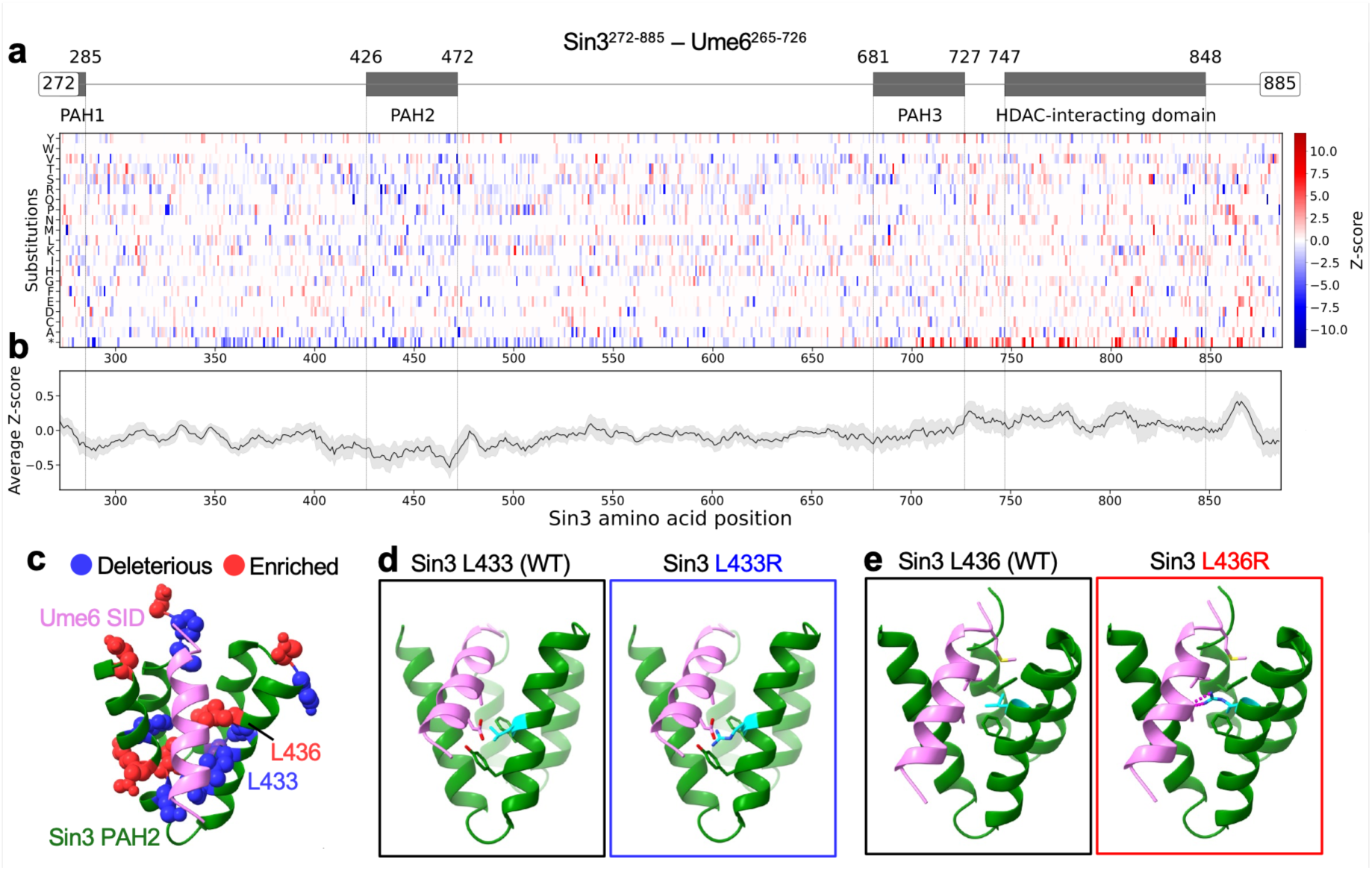
| Mutational scanning defines conserved residues essential for Ume6 engagement. **a**, Genophenogram of mutational scanning for the Sin3 (residues 272-885)-Ume6 (residues 265-726) interaction; enriched (red) and deleterious (blue) variants are shown with Z-scores (colour scale). **b**, Rolling average of Z-scores (11-residue window ± s.d.). **c**, Top 10 conserved deleterious and enriched variants mapped onto the Sin3 PAH2-Ume6 SID structure (PDB: 6XAW). **d**,**e**, Structural context of representative deleterious (L433R) and enriched (L436R) mutations.

Analysis of truncation variants (stop codons) in the final row of the genophenogram further delineated the minimal region required for binding. Indeed, a stark transition from deleterious to neutral scores near residue ∼600 indicates that sequences N-terminal to this position are sufficient for Ume6 engagement. In contrast, mutations beyond residue ∼700 were predominantly enriched. Since Ume6 binds Sin3 through residues ∼420-475, the Sin3 C-terminal region encompassing the PAH3 and HID domains could recruit endogenous Sin3L subunits promoting higher-order complex assembly. Enriched mutations in this region could thus favor formation of a tripartite configuration that cooperatively stabilizes the Sin3-Ume6 interaction.

To integrate mutational and structural insights, deleterious and enriched substitutions were mapped onto conserved residues of the Sin3 PAH2 domain in the crystal structure (**Fig. 4c and Supplementary Table 3**). Two representative alleles were picked to illustrate the structural basis of these effects. Substitution of Leu433 by an arginine introduces steric interference with the Ume6 α-helix (**Fig. 4d**), consistent with loss of binding. Conversely, replacement of Leu436 by an arginine establishes two stabilizing hydrogen bonds at the interface (**Fig. 4e**). Together, these analyses define conserved determinants within the minimal Sin3 PAH2 interface whose perturbation produces gain- or loss-of-function effects, thereby establishing the molecular basis of SID-containing transcription factor engagement and the regulatory principles governing recruitment by the Sin3L complex.

## DISCUSSION

Here, we presented an integrated structural dynamics platform that reveals how HDAC-HAT bridging proteins dynamically engage with Sin3L (**Fig. 1d and Fig. 5a**). Building on high-resolution cryo-EM structures of native assemblies that define a rigid ∼0.4 MDa core, we combined cryo-EM density maps, XL-MS and AlphaFold3-HADDOCK-based modeling to extend the architecture to ∼0.9 MDa, resolving flexible regions previously uncharacterized (**Fig. 5b and Extended Data Fig. 8**). The modest reduction in crosslink agreement in the expanded model is consistent with intrinsic conformational heterogeneity. The resulting framework positions key regulatory elements – including the Sin3 PAH1 and PAH2 scaffolds, the Ash1 zinc finger and the PHD motifs of the histone readers Cti6 and Pho23 – within a dynamic yet spatially organized periphery. Notably, a recent study of recombinantly purified Sin3L bound to a mono-nucleosome also positioned Cti6 at approximately the same site, which further supports our findings^46^. Fragment-resolved interactome mapping incorporated the transient transcription factor Ume6, and crystallographic and mutational scanning analyses defined determinants of its interface with Sin3 at residue-level precision, offering mechanistic insights into evolutionary trajectories and disease processes. Collectively, these data establish a hierarchical Sin3L architecture in which a stable catalytic core is surrounded by adaptable modules that enable locus-specific assemblies and regulatory functions, likely conserved across eukaryotes^47^.

**Fig. 5.**
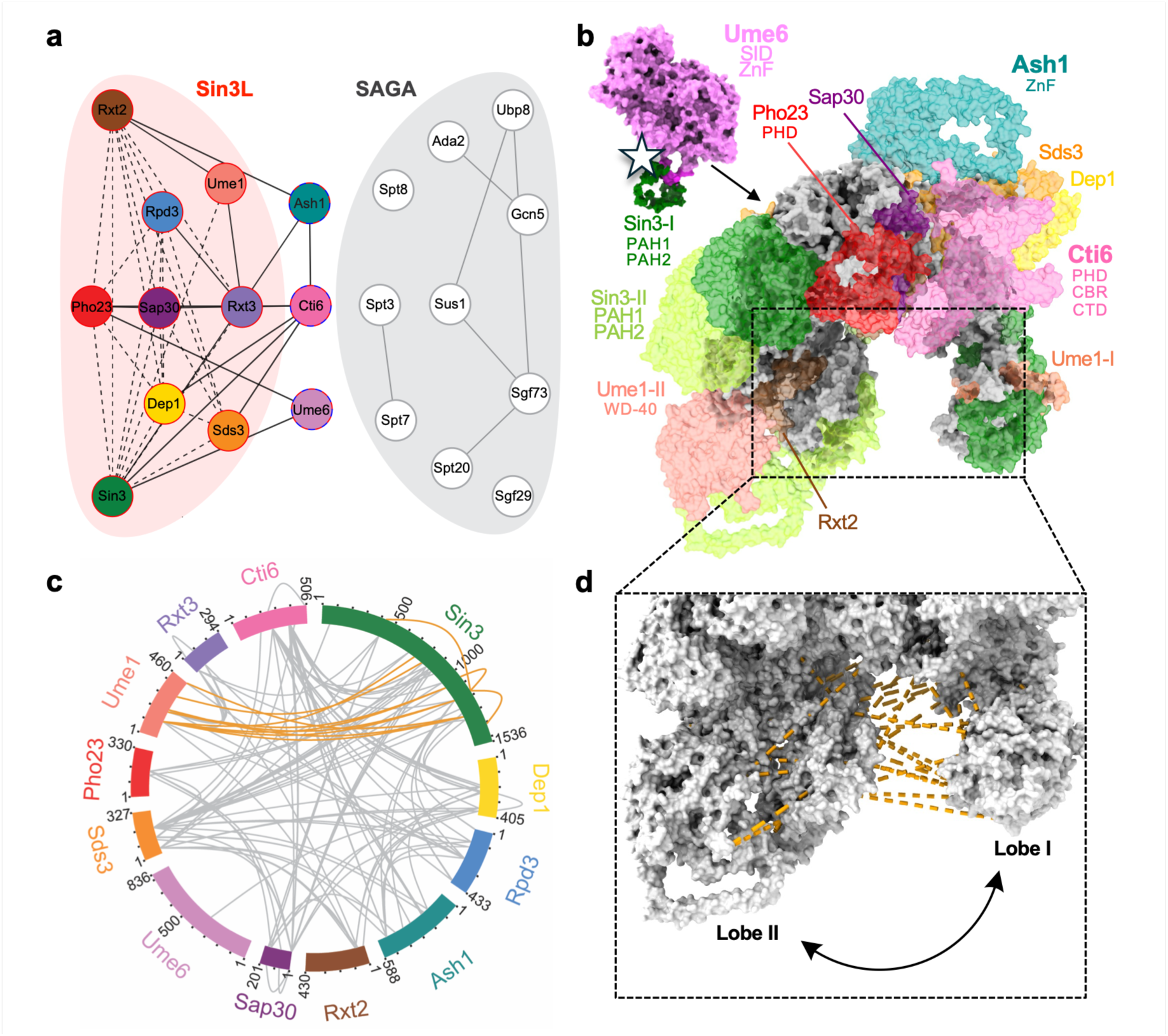
| Dynamic Sin3L peripheries expand the structural model towards SAGA. **a**, Interaction network of the extended Sin3L (red) and SAGA (grey) structures. Solid black edges indicate interactions contributed by newly resolved regions in this study; dashed edges represent core contacts identified in this study and in published structures^21,22^. Solid grey edges represent known structural information for SAGA subunits. **b**, Integrative model of Sin3L with Cti6 and Ash1. Newly resolved domains are coloured over the core surface (grey). The Sin3 PAH2-Ume6 interface and the inter-lobe clamp are indicated. The star at the Ume6-Sin3 interface schematically represents conserved residues essential for Sin3L engagement identified by mutational scanning (Fig. 4). **c**, Crosslinks unmatched to the extended model (Cα-Cα >37 Å) in grey. Sin3-Sin3 and Sin3-Ume1 crosslinks potentially bridging the two lobes are highlighted in orange. **d**, Close-up of the inter-lobe clamp with highlighted crosslinks from (**c**), illustrating potential conformational opening and closing upon binding of additional interactors or nucleosome engagement.

The extended structures provide mechanistic insight into how this organization is achieved. Complete modeling of Sin3 PAH1 and PAH2 on lobe II reveals their close spatial proximity, positioning these domains to enable competitive or cooperative engagement of SID-containing factors. Although our model includes a single Ume6 molecule bound to PAH2, additional PAH domains remain accessible, and multiple Ume6 copies may potentially be accommodated. This also raises the possibility that other partners, such as Cti6, could bind adjacent domains to support dynamic, combinatorial assemblies analogous to the Cti6-Ash1 interaction. This is in line with previous observations that Cti6 deletion leads to loss of both Ash1 and Ume6^18^, and unmatched crosslinks between Cti6 and other subunits (e.g. Cti6 K250 and Dep1 K355 or Sds3 K41) further support this hypothesis. Modeling of Rxt2 indicates that it can occlude the catalytic sites of both Rpd3-I and Rpd3-II (**Extended Data Fig. 4h,i**), suggesting a mechanism to modulate HDAC accessibility and enzymatic activity. Additional unmatched crosslinks and inter-lobe restraints are consistent with clamp-like opening and closing motions (**Fig. 5c,d**), implying that conformational transitions coordinate nucleosome engagement and positioning *in cellulo*. Together, these observations challenge a purely repressive view of HDAC complexes and instead support a model in which Sin3L functions as a dual-function regulatory hub integrating activating and repressive activities through dynamic factor exchanges.

Important limitations and open questions remain. Our integrative models rely on ensemble-averaged cryo-EM reconstructions and XL-MS restraints, which cannot fully resolve low-abundance or short-lived conformers. Portions of the expanded densities remain unassigned, and the reconstructed assembly does not fully account for the estimated native mass of ∼1 MDa, implying additional context-dependent components. Biochemical evidence suggests that non-canonical subunits may associate with Sin3L under specific experimental conditions, yet their structural incorporation remains unknown. Although we establish strong structural connections of the dual-role regulators Cti6, Ash1 and Ume6 with Sin3L, their physical interactions with SAGA subunits were not directly examined here. Systematic analyses will therefore be required to define the extent of their engagement with SAGA and to further elucidate the functional and structural interplay between the Sin3L and SAGA machineries. Moreover, the absence of native nucleosome-bound states leaves open how peripheral modules reposition upon chromatin engagement and how histone mark recognition is coupled to catalytic activity. Capturing defined functional states will be essential to address these questions.

Given the evolutionary conservation of the Sin3L complex from yeast to human, including conserved PAH-mediated transcription factor recruitment and HDAC scaffold organization^48^, the extended structures provide a mechanistic framework for understanding the assembly principles of human SIN3-HDAC architectures. They offer structural insight into how the dimerization of flexible HDAC C-terminal regions could be promoted by adjacent subunits, which remains poorly characterized in human complexes and may contribute to higher-order assembly. More broadly, the integrative platform established here offers a blueprint for resolving dynamic multisubunit assemblies at molecular resolution beyond Sin3L. Extending fragment-resolved interactomics and mutational scanning to disease-associated variants should further clarify how subtle genetic perturbations reconfigure three-dimensional subunit organization to drive epigenetic dysregulation.

## METHODS

### Culture of yeast cells

All yeast cells used in this study are from the *Saccharomyces cerevisiae* species and their genotypes are presented in the **Supplementary Table 4**. For each experiment, yeast cells from the glycerol stocks were streaked on complete (yeast extract-peptone-dextrose, YPD) or on selective solid media containing glucose as the carbon source, and grown for 3-5 days at 30°C to obtain fresh isolated single colonies. Cells were then cultured at 30°C under shaking conditions (200-220 rpm) in the appropriate medium until reaching the desired cell density. For purifications, yeast cells were grown for approximately 16 h with a starting OD_600 nm_ ∼0.05.

### Conditions for yeast cell selections

The media used in this study were previously described^49^. Cells were cultured as explained above. For culture conditions requiring specific selection(s), synthetic complete (SC) media lacking one or several components (e.g. SC-L-W corresponds to SC lacking leucine and tryptophan) were used. The letters L, W, A, H and U refer to leucine, tryptophan, adenine, histidine and uracil, respectively. 5-fluoroorotic acid (5FOA) or 3-amino-1,2,4-triazole (3AT) were added to the different media at the indicated final concentrations. When the *NatMX*, *KanMX* or *HphMX* cassette was used to delete a specific gene in the yeast genome, nourseothricin (clonnat), geneticin (G418), or hygromycin B was added to YPD or SC media to select for transformants.

### Yeast strains

The different gene deletion strains were generated via double homologous recombination, by transforming yeast cells with gene replacement cassettes as previously described^50–52^. Briefly, the marker cassettes were PCR-amplified with primers containing 5’-extensions (45-50 bases) directly adjacent to them, and homologous to the promoters or terminators of the targeted genomic loci. The resulting mutants were selected as described above, and single colonies were picked and purified before lysing cells and validating the gene deletion/marker cassette by PCR amplification followed by agarose gel electrophoresis using specific primer pairs. MaV208 was generated from MaV108 by disruption of two drug exporter genes using the *KanMX* gene cassette, and the *HIS3* marker. First, MaV108^33^ was transformed with a DNA fragment that was amplified by PCR reaction using a plasmid pLexA (Clontech) as a template for *HIS3* marker. Both ends of the PCR product included a short region homologous to *PDR5*. Transformed yeast cells were plated on SC plates lacking histidine (SC-H) and used to select a new strain named MaV118. This new yeast strain was further transformed with a DNA fragment that was amplified by PCR reaction using the plasmid pUG6 as a template for the *KanMX* gene cassette^53^. Both ends of the PCR product included a short region homologous to *SNQ2*. Transformed cells were plated on YPD plates supplemented with 200 mg/L G418, and a new strain MaV208 was selected from the resistant colonies. For the deletion strains, the *NatMX* cassette was PCR-amplified from plasmid p4339 (gift from Charles Boone, University of Toronto), with primers specific to the TEF promoter and TEF terminator. For the wild-type (WT)+*GAL4* control condition, WT MaV208 yeast cells were transformed with a pDEST-DB-scGal4 plasmid to induce expression of the Gal4 transcription factor and activation of *SPAL10::URA3* from the upstream activating sequences (*UAS*) present in the promoter^33^. For JOY128 (*(spal10::ura3)Δ::HphMX@ura3*), the *SPAL10::URA3* region corresponding to the Gal4 binding sites, the *URS1* motif from *SPO13*, the *URA3* reporter gene^33^, and 53 bases of the *URA3* terminator, was replaced by the *hph* marker. The HOY13 and HOY14 strains were generated by PCR amplifying the *SPAL10::URA3* reporter system from MaV103 genomic DNA and inserting it at the *URA3* locus in the Y8800 and Y8930 yeast strains, respectively.

### Agar spot test assays

When cells reached OD_600 nm_ ∼0.6-1 in ∼1mL, population densities were evaluated via counting grid microscope slides. Cells were then centrifuged, the supernatants were discarded and the pellets washed twice with double-distilled water. The resulting pellets were then diluted in different volumes of water to obtain similar population densities for all strains. From these starting suspensions, serial dilutions (1:10) were made for every strain, and ∼8-10 μL of culture were spotted for each dilution, on the indicated solid agar media. Plates were then incubated according to the indicated conditions.

### RNA extraction from yeast cells

After yeast cells reached OD_600_ ∼0.5 in 0.8-1 mL of YPD, they were grown for two more hours at room temperature before being centrifuged to remove supernatant and freezing pellets on dry ice. Yeast cells (three biological replicates per strain) were then harvested for RNA extraction using a RiboPure RNA Purification Kit (Invitrogen, cat# AM1926), which involves mechanical cell wall disruption, phenol extraction of the lysate, and RNA purification using glass fibre, and filter-based RNA purification columns. RNA concentration was measured for individually purified RNA samples using a NanoDrop spectrophotometer.

### RT-qPCR measurements

The RT-qPCR protocol used in this study was described elsewhere^54^. Primers used in this study to amplify yeast genes are presented in the **Supplementary Table 5**. Approximately 1 μg RNA per sample was used for reverse transcription. Using a thermocycler, RNA was denatured in the presence of 0.2 μg of 18-mer oligo-dT by heating the plate at 65°C for 10 min. The plate was then immediately cooled down on ice to anneal oligo-dT on the poly-A tail of mRNA. A master mix of AffinityScript Multiple Temperature Reverse Transcriptase (Agilent, cat# 600107) was prepared following the manufacturer’s protocol and mixed with RNA samples previously annealed with the oligo-dT. The plate was incubated at 42°C for 2 h to generate cDNAs from mRNA templates. The reverse transcriptase was then denatured by heating the plate at 70°C for 15 min. The resulting cDNA was diluted (1:20) in sterile, double-distilled water, and stored at -20°C until used for qPCR. To measure gene expression, the diluted cDNA was mixed with 5 μL PowerUp SYBR Green master mix (Applied Biosystems, cat# A25743), and the mix of primers (forward+reverse: 0.3 μM final concentration) for a total volume of 10 μL. Three technical replicates were generated per cDNA sample in a 384-well PCR plate (Applied Biosystems, cat# 4343370). The plate was tightly sealed with an optical adhesive film (Applied Biosystems, cat# 4311971). Quantitative PCR was conducted with a QuantStudio real-time PCR system or an ABI Prism system (both from Applied Biosystems), with the following experimental settings: 50°C/2 min, 95°C/10 min, and 40 cycles of 95°C/15 s and 60°C/30 s. Dissociation curves were checked for each PCR product to assess the specificity of PCR amplicons and products were sequenced to confirm target genes. Gene expression measured by RT-qPCR was calculated using the following equation:

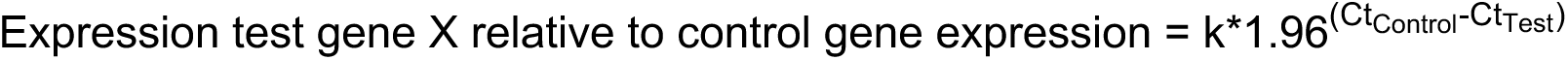

Where k is an arbitrary multiplier (k = 1000 in this study), 1.96 is the PCR amplification efficiency constant, Ct is the cycle threshold, Ct_Control_ is the Ct value for the housekeeping gene used as control for normalization (*UBC6*^55^), and Ct_Test_ is the Ct value for the tested gene. Replicates corresponding to WT yeast cells were averaged and other data points normalized using the WT values as references. *TAF10* was used as a specificity control gene as it is not known to be regulated by an HDAC complex. Three technical replicates were collected and averaged for each biologically independent sample per group. In the rare cases where the value of a technical replicate was undetermined or gave the same result as a control empty well in the plate, it was excluded from the average calculation.

### Hierarchical clustering of Deleteome data

Hierarchical clustering of Sin3L gene expression profiles from the Deleteome compendium^36^ was performed using seaborn clustermap in Python3. Knockout strains were clustered using average linkage with Euclidean distance, while genes were not clustered. The heatmap displays log₂ fold changes relative to wild type using a diverging colourmap (seismic) with a scale from -2 to +2.

### Principal component analysis of the Deletome

Selected gene expression data from the Deletome compendium^36^ were analysed using principal component analysis (PCA) to visualize transcriptional similarities across the Sin3L subunit knockout strains. Log_2_ fold change values were used as input, with knockout strains as observations and genes as features. PCA was performed using scikit-learn in Python3, and the 2D principal component plots were plotted using matplotlib.

### Network construction and visualization

Pairwise Pearson correlation coefficients were computed across genome-wide expression profiles (6,077 genes) for 21 yeast deletion strains from the Deleteome dataset^36^. Statistical significance was assessed using Student’s t-distribution (n-2 degrees of freedom) with Benjamini-Hochberg false discovery rate (FDR) correction. Edges were retained based on dual thresholds: for core Sin3L complex members (*rpd3Δ*, *sin3Δ*, *dep1Δ*, *pho23Δ*, *sap30Δ*, *sds3Δ*, *ume1Δ*, *rxt2Δ*), q < 0.005 and |r| ≥ 0.35; for all other pairs, q < 0.05 and |r| ≥ 0.20. Stricter thresholds for core-to-core interactions account for expected strong functional coupling within the HDAC complex and reduce network density.

Transcriptional similarity network layout was generated using the Fruchterman-Reingold force-directed algorithm (NetworkX v3.1; k=1.3, 350 iterations, seed=42 for reproducibility; layout stability confirmed with 500 and 1,000 iterations) with Fisher z-transformed correlation coefficients [z = arctanh(r)] as edge weights. Manual adjustments were applied post-layout to improve interpretability: Sin3L and SAGA complex nodes were radially dispersed around their respective centroids to reduce crowding, and bridge nodes (*cti6Δ*, *ash1Δ*) were positioned between complexes to emphasize their intermediate functional roles. Minor positional adjustments to individual nodes (*sds3Δ*, *ada2Δ*) avoided visual overlaps while preserving network topology. Colouring was modified to separate Sin3L and SAGA complex nodes and highlight dual-function nodes. Structural networks were prepared by maintaining the layout of the transcriptional similarity network, with edges denoting the presence of a protein-protein interaction interface between two nodes in the structures of the Sin3L core, Sin3L extended or SAGA complexes. Structural information for the SAGA complex was collected from PDB: 3MHH, 6CW2, 6T9I, and 6T9L. Transcriptional profiling figures were generated using Python 3.10 with NetworkX v3.3, NumPy v1.26, Pandas v2.2, SciPy v1.14, statsmodels v0.14, and Matplotlib v3.9 and adjusted using Adobe Illustrator. Structural networks were built using Adobe Illustrator.

### Single-step purification of the Sin3L complex

Yeast strains modified with a TAP tag at the C-terminus of Sin3, Sap30 or Rxt3, and the parental BY4741 strain (used as the no-tag control) were obtained from Horizon Discovery, except for Sap30-TAP (gift from Stephen Buratowski, Harvard Medical School). A 12 L culture of the TAP-tagged yeast strain was grown as described above. The published TAP procedure^22,56^ was modified and optimized to purify Sin3L complexes with high yield and purity suitable for systematic cryo-EM studies. Cells were pelleted from the culture by centrifuging at 4,000 x *g* for 10 min and resuspended in 2 mL TAP Extract Buffer (40 mM HEPES pH 7.6, 350 mM KCl, 50 mM NaCl, 2 mM MgCl_2_, 4 mM CaCl_2_, 10% glycerol, 0.1% Tween-20 with 2 mM DTT, 1 mM PMSF, 1 µg/mL pepstatin A, 2 µg/mL leupeptin and 0.33 mg/mL benzamidine added fresh) per litre of overnight culture. The cell suspension was frozen by passing it dropwise into a liquid nitrogen bath. The frozen yeast cell beads were lysed at 4°C using a coffee grinder (Krups F203) prechilled with liquid nitrogen with 2 rounds of grinding for 30 s each (the coffee grinder was loaded with beads equivalent to 4 L of starting culture at a time). The lysed cell powder was resuspended with 3 mL of TAP Extract Buffer per litre of cells and thawed at room temperature. The thawed lysate was transferred to 4°C and stirred with 6 µL of DNAse I (5 mg/mL; Roche) and 15 µL of RNase A (2 mg/mL; Thermo Scientific) per litre of starting culture for 20 min. The lysate was then stirred for another 10 min with 4 mg of heparin powder per litre of starting culture (Sigma-Aldrich) dissolved in water and centrifuged at 20,000 x *g* for 10 min at 4°C. The supernatant was further clarified by centrifuging at 100,000 x *g* for 30 min at 4°C. From the clarification, the clear yellow middle layer was retained and split into three tubes (one per 4 L of starting culture). The middle layer in each tube was incubated with one mini protease inhibitor cocktail tablet (Roche) and 500 µL of calmodulin (CaM) resin (Cytiva) slurry (corresponding to ∼200 µL of CaM beads) previously washed with CaM Wash 1 Buffer (20 mM HEPES pH 8.0, 750 mM KCl, 50 mM NaCl, 2 mM MgCl_2_, 2 mM CaCl_2_, 5% glycerol, 0.05% NP-40) on a gently rotating wheel for 16-18 h at 4°C. For each tube, the flow-through was eluted using a different Econo-Pac chromatography column (Bio-Rad, 7321010) and the beads were gently washed once with 15 mL TAP Extract Buffer, 15 mL CaM Wash 1 Buffer and, lastly, with 15 mL CaM Wash 2 Buffer (20 mM HEPES pH 8.0, 150 mM KCl, 50 mM NaCl, 2 mM MgCl_2_, 5% glycerol, 0.02% NP-40). For washing, the CaM Wash buffers were supplemented with 2 mM DTT. Proteins were eluted with 6 additions of 400 µL CaM Elution Buffer (20 mM HEPES pH 7.6, 50 mM KCl, 30 mM NaCl, 2 mM MgCl_2_, 4 mM EGTA, 1.5% glycerol with 1 mM DTT added fresh). Elutions 1-3 were collected immediately. Elutions 4, 5 and 6 were collected after incubation of 2.5, 5 and 10 min, respectively, with the calmodulin resin. Elutions 2-6 were grouped and concentrated to ∼70 µL using a 100 kDa-cutoff Amicon ultra-centrifugal filter (Merk Millipore) at a maximum speed of 4000 x *g* at 4°C. Concentration of the Sin3L complex was measured at 280 nm absorbance with the NanoDrop One UV-VIS spectrophotometer using calculated molecular weight and extinction coefficient of 1.057203 MDa and 919095 M^-1^cm^-1^, respectively. The molecular weight reference used was calculated using the following stoichiometry: 2:2:2:1:1:1:1:1:1:1:1:1 for Rpd3, Sin3, Ume1, Sds3, Dep1, Sap30, Rxt2, Pho23, Rxt3, Cti6, Ash1, Ume6, respectively, and the extinction coefficient was derived using the online Expasy platform (https://web.expasy.org/protparam/). The concentrated sample was snap-frozen in liquid nitrogen and stored at -80°C. Each purification was independently repeated at least three times and yielded highly similar results.

### Characterizations of Sin3L complex samples

For SDS-PAGE characterizations, samples of the purified Sin3L complexes were diluted to 5 μg each and the no tag sample was diluted to 1 μg (due to a ∼10x lower concentration of proteins obtained for that purification). Samples were mixed with 5x standard protein loading buffer and incubated for 10 min at 95°C before being loaded in a 4-20% Mini-PROTEAN^®^ TGX^TM^ Precast Gel (Bio-Rad). A ladder with precision plus protein standards, unstained (Bio-Rad), including reference bands at 20, 50 and 100 kDa corresponding to 37.5, 187.5 and 37.5 ng was loaded (2.5 μL loaded from the commercial stock). Gels were run for 35 min at 200 V, visualized for stain-free imaging with the Gel Doc EZ System (Bio-Rad) and analysed with the included Image Lab Software. For Coomassie staining, gels were stained with a 10% acetic acid solution containing Coomassie Blue G-250 overnight and subjected to multiple rounds of destaining with a 10% acetic acid solution before visualization with the Image Lab Software using the white tray for Coomassie, copper, silver and zinc stains. For dynamic light scattering (DLS) experiments, the cuvette mode of a DynaPro NanoStar (Wyatt Technologies) was used with the chamber maintained at 4°C. For each sample, 8 μL of the concentrated sample was loaded on to disposable cuvettes (Wyatt Technologies), and 30 acquisitions were collected to generate an average reading with the Dynamics software (Wyatt Technologies).

### Assessment of deacetylase enzymatic activity

The HDAC-Glo I/II^TM^ screening system^57^ (Promega, cat# G6420) was adapted to measure the activity of the purified Sin3L complexes and their inhibitions by Trichostatin A (TSA). HDAC enzymatic activity was measured following the manufacturer’s protocol. Briefly, linear ranges of the assay were determined by measuring luminescence signals, in a serial dilution manner, of the purified complexes diluted in the HDAC-Glo I/II buffer. Dilution factors corresponding to signals in the linear ranges were used to work under optimal conditions of sensitivity. To test the effect of compounds on HDAC enzymatic activity, equal volumes of purified complexes at 400 nM (15 μL), and inhibitors (15 μL) were added to the wells of a white, flat bottom 96-well plate. The plate was then briefly centrifuged and gently shaken for 1 min at 700 rpm. The plate was tightly sealed to prevent evaporation and incubated at room temperature for 1 h. Following this incubation, 30 μL of luciferase substrate were added to each well. The 96-well plate was then briefly centrifuged and shaken for 1 min at 700 rpm before being tightly sealed and incubated for 30 min at room temperature. Luminescence signals were measured using a GloMax^®^ Navigator microplate reader (Promega), with 1 s integration time per sample.

### Cryo-EM sample preparation

Each Sin3L complex sample was crosslinked with 1 mM bis(sulfosuccinimidyl)suberate (BS^3^) for 1h on ice. The crosslinking reaction was quenched with 30 mM Tris-HCl pH 8.0 for 15 min on ice. The sample was supplemented with 0.2% CHAPS immediately prior to grid making. Cryo-EM grids were prepared using a Vitrobot Mark IV grid plunger (Thermo Fisher Scientific). Quantifoil Cu300 R1.2/1.3 + 2 nm Carbon (ultra-thin) holey carbon grids were glow-discharged for 15 s at 10 mA with the chamber pressure set to 0.3 mbar (PELCO easiGlow Glow Discharge Cleaning System; Ted Pella). The glow-discharged grids were mounted in the Vitrobot Mark IV sample chamber at 4°C and 100% humidity and 4 µL of the sample was applied. The sample was incubated on the grid for 5 s, double-side blotted with Whatman 2 filter paper (Ted Pella) for 5 s at 0 N force and plunge-frozen in liquid ethane. The vitrified grids were clipped and loaded on a 200 keV Glacios cryo-transmission electron microscope (Thermo Fisher Scientific) equipped with an autoloader and a Selectris energy filter hosted by the Rega Institute for Medical Research, KU Leuven. Multi-frame movies were recorded in counting mode using a Falcon4i direct electron detector with EPU software version 3.5.1 (Thermo Fisher Scientific). Each dataset was collected at a nominal magnification of 100,000x, leading to a pixel size of 1.16 Å, with a total dose of 40 e/Å^2^. The raw movies were stored as either gain-corrected MRC or TIFF files.

### Cryo-EM data processing and modelling

For each dataset, individual movies were corrected for motion and aligned using MotionCor2^58^ as implemented in Relion 4.0^59^. Remaining processing of the high-quality density maps from each dataset was done in CryoSPARC^60^. Patch CTF was used to estimate the contrast transfer function (CTF) parameters for each motion-corrected micrograph and retain micrographs with a good CTF fit resolution (<6.5 Å). The blob picker tool in CryoSPARC was used to pick the initial particle stack and 2D classification was performed twice to clean the particles after extraction with a box size of 448 pixels. Three *ab initio* maps were generated from the cleaned particle stack and heterogeneous refinement was used for the removal of junk particles. The class containing particles of the Sin3L complex were reclassified and cleaned by 2D classification to produce the final particle set. These were used for non-uniform refinement^61^, global CTF refinement and local refinement jobs to obtain a final map and calculate a local resolution map. Map quality was assessed using the validation Fourier shell correlation(FSC)^62^ and orientation diagnostics tools in CryoSPARC and the remote 3DFSC processing server. Final sharpened maps of the Sin3L core from the Sap30-TAP, Sin3-TAP and Rxt3-TAP data were determined at 3.44, 3.37 and 3.61 Å, respectively.

The maps with extended density were processed using Relion 5.0. After motion correction, the CTFFIND4^63^ package was used to estimate the CTF parameters and filter good micrographs. The Laplacian-of-Gaussian (LoG) picking function of Relion was used to autopick the initial particle stack which were extracted for further processing. 2D classification was performed using the VDAM algorithm to clean the particle stack. In the case of the Sap30-TAP dataset, 2D classification was performed after splitting the initial particle stack into two segments. 3D classification was performed with either three or four 3D classes after *de novo* model generation. The best class was selected from the classification job and further refined with local 3D classification with two classes. Particles from best class were processed with CTF refinement and Bayesian polishing^64^ and input to 3D auto-refinement and postprocessing jobs to determine the final unsharpened and sharpened maps, respectively. Final unsharpened maps of the Sin3L complex with extended densities were determined at 3.48, 3.78 and 3.78 Å from the Sin3-TAP, Sap30-TAP and Rxt3-TAP data, respectively.

The high-resolution maps generated by data processing with cryoSPARC were used to model structures of the Sin3L core complex^21^. A published structure of the Sin3L core (PDB: 8HPO^21^) was used as the starting model for iterative rounds of model building with Coot 0.9.8.95^65^, coupled with real space refinement using Phenix 1.21.2^66–69^. Structures were visualized and analysed using UCSF ChimeraX^70^.

### DSSO crosslinking for mass spectrometry

Samples of purified Sap30-TAP tagged Sin3L complex were diluted to get 15 μg per replicate and crosslinked with 5 mM disuccinimidyl sulfoxide (DSSO) for 45 min on ice, as previously reported^12,42^. The crosslinking reaction was quenched with 30 mM Tris-HCl pH 8.0 for 15 min on ice. After quenching, 20% of the sample was retained for verifying completion of the crosslinking reaction by SDS-PAGE. The rest of the sample was prepared for quantitative and crosslinking mass spectrometry analyses using TCA precipitation. For this, an equal volume of 100 mM Tris-HCl pH 8.0 was added to the sample solution, followed by TCA in a 1:4 volume ratio to the sample to give a final concentration of 20% of TCA. The samples were incubated at 4°C overnight. Samples were spun down at 14,000 rpm for 30 min at 4°C and then washed twice with an equal volume of ice-cold acetone and spun at 14,000 rpm for 10 min at 4°C after each wash. The supernatant was aspirated and the pellet was air dried under the hood for at least 4 h. Prepared samples were stored at -20°C till processing. Samples from the no tag purification were prepared using the same procedure with a starting amount of 4.5 μg for each replicate.

### Crosslinking mass spectrometry (XL-MS)

Purified and crosslinked Sin3L complexes were prepared using an adapted Micro S-Trap protocol (Protifi). Briefly, protein pellets were resuspended in S-Trap 1X lysis buffer containing 5% SDS. Cysteines were reduced with 10 mM DTT using a thermomixer at 55°C and 850 rpm for 1 h and alkylated using 25 mM indole-3-acetic acid (IAA) in the dark using a thermomixer at 20-24°C and 850 rpm for 30 min. Following the S-trap protocol, S-Trap Acidifier and S-Trap Binding/Washing Buffer were added sequentially, and the samples were loaded onto the column and washed with S-Trap Binding/Washing Buffer. To digest proteins, 1 μg of Trypsin (Promega) and 100 ng of Lys-C (Promega) in 50 mM TEAB were added to the column and samples were incubated overnight in a stationary thermomixer at 37°C. Following digestion, peptides were eluted from the columns with 3 steps containing 50 mM TEAB, 0.2% formic acid, and 80% acetonitrile in 0.1% formic acid, respectively. Peptides were then dried down using a SpeedVac. Lyophilized peptides were then resuspended in 20 μL 0.1% formic acid for LC-MS/MS analysis. To identify binding partners and crosslinks, purified and digested, Sin3L complexes were separated by a Thermo Vanquish UHPLC system coupled with a Thermo Ascend Tribrid Mass Spectrometer (Thermo Scientific). Samples underwent an injection of 3 μL with a high resolution Orbitrap method (XL-MS2) and a stepped Higher-energy Collisional Dissociation (sHCD) as well as a separate injection of 15 μL utilizing both the Orbitrap and Ion Trap (XL-MS3) with both Collision-Induced Dissociation (CID) followed by HCD (CID-HCD). Liquid chromatography was performed by separating peptides with the Thermo Vanquish UHPLC system using a Thermo Acclaim PepMap 5 mm (length) x 1 mm (internal diameter) x 5 μm (particle size) C18 trap column and an IonOpticks Aurora Ultimate 25 mm x 75 μm x 1.7 μm C18 analytical column heated to 50°C. Using 0.1% formic acid as Buffer A, 80% acetonitrile in 0.1% formic acid as Buffer B as mobile phases as well as a flow rate of 180 nL/min, the columns were equilibrated to 1% B before applying a gradient of 8% B for 1min, increasing to 33% B over 100 min, increasing to 60% B over 24 min, then increasing to 90% B over 4 min, and finally undergoing washing at 90% B over 10 min. Peptides eluted from the column were introduced into the mass spectrometer using a Nanospray Flex Ion Source (Thermo Scientific) with a spray voltage of 1700 V. Peptides first passed through the FAIMS Pro Duo (Thermo Scientific) at the following FAIMS CV settings: -50 V, -60 V, -75 V. The system cycled through the following scan events at each FAIMS CV with a maximum cycle time of 2 s for the XL-MS2 method and 3 s for the XL-MS3 method. Each MS1 scan event was detected by the Orbitrap at resolution 60,000, scan range 375-1600 m/z, MIT 123 ms, and ACG target 400000. In the XL-MS2 method, MS2 scans were triggered with peptide charge states 3-7 and a dynamic exclusion of 45 s. Each MS2 scan was acquired with the Orbitrap with an isolation window of 1.6 m/z, sHCD collision energies 25, 30, and 35 at resolution 30000, MIT 150 ms, and AGC Target 100000. In the XL-MS3 method, MS2 scans were triggered with peptide charge states 4-8 and a dynamic exclusion of 45 s. Each MS2 scan was acquired with the Orbitrap with an isolation window of 1.6 m/z, CID collision energy of 25% at resolution 30000, MIT 120 ms, and AGC Target 75000. An MS3 scan was triggered by a mass difference of 31.9721 at 10 ppm corresponding to DSSO fragmentation. Each MS3 scan was acquired by the IonTrap in Rapid scan mode with an isolation window 2.5 m/z, HCD collision Energy 32%, MIT 200 ms, and AGC Target 20000.

### Database search and crosslinking analysis

To characterize enrichment and binding partners, XL-MS2 raw files were searched using FragPipe and MSFragger^71^ against negative control files. The yeast database, common contaminants, and decoys were loaded directly from FragPipe and were searched with the following settings: DDA mode, trypsin digestion KR, maximum missed cleavage = 2, precursor mass tolerance +/- 10 ppm, fragment mass tolerance +/- 20 ppm, variable modification Oxidation on M 15.9949 Da and N-term acetylation 42.0106 Da, fixed modification Carbamidomethyl on C 57.02146 Da, MS1 quant without Match between runs, and other default settings. Results were analysed with SAINTexpress^72^ with default settings and FragPipe Analyst^73^ with the following settings: Data Type = LFQ, Intensity Type = spectral counts, DE Adjusted p-value cutoff = 0.05, DE Log2 fold change cutoff = 1, No normalization, No imputation, Type of FDR correction = Benjamini Hochberg.

To identify crosslinks, XL-MS2 and XL-MS3 raw files were searched separately using Proteome Discoverer 3.2 (Thermo Scientific) and XlinkX^41^. The yeast database was downloaded from UniProt containing 6650 sequences and was combined with a common contaminant database containing 298 sequences. A first Sequest search was performed using the following settings: trypsin digestion KR, maximum missed cleavage = 2, precursor mass tolerance +/- 10ppm, fragment mass tolerance +/- 0.02 Da, variable modifications Oxidation on M 15.9949 Da, DSSO hydrolysed on K 176.19165 Da, DSSO-Tris on K 279.07766 Da, and N-term acetylation 42.0106 Da, fixed modification Carbamidomethyl on C 57.02146 Da, and other default settings. A second XlinkX search was performed using DSSO as the crosslinker and either MS2 strategy for XL-MS2 method raw files or MS2-MS3 strategy for XL-MS3 method raw files with the following settings: trypsin digestion KR, maximum missed cleavage = 2, precursor mass tolerance +/- 10 ppm, FTMS mass tolerance +/- 20 ppm, ITMS mass tolerance +/- 0.5 Da, variable modifications Oxidation on M 15.9949 Da, fixed modification Carbamidomethyl on C 57.02146 Da, and other default settings. Resulting crosslinks were further analysed using custom RStudio scripts, custom Python scripts, XiView^74^, XMAS^75^, and UCSF ChimeraX^70^.

### Subcomplex modelling with XL-MS and AlphaFold3

Sub-complexes of Sin3L were defined based on prior biochemical and interactome data and confirmed by the XL-MS data. AlphaFold3^39^ was employed to predict the 3D structures of these sub-complexes. For each sub-complex, multiple models were generated. We used a custom scoring approach to evaluate the compatibility of predicted models with XL-MS-derived cross-linking distance restraints (Cα–Cα distance ≤37 Å). Only sub-complex structures that showed the highest number of satisfied cross-links were retained for further analysis.

### Complex assembly via guided molecular docking

The top-scoring sub-complexes were assembled into the Sin3L extended structure using the HADDOCK2.4^37,38,76^ docking platform. Input structures were prepared using PDBTools^77^ and custom-made python scripts. Distance restraints were derived from cryo-EM data by selecting lysine to lysine contacts at a fixed distance (Cα–Cα distance ≤37 Å) to be used as pseudo-crosslinks and combined with XL-MS results. These distance restraints were converted into unambiguous interaction restraints and used to guide the docking process for the core integrated model as well as the addition of Ash1 and Cti6. Distance restraints derived from the Sin3 PAH2-Ume6 SID crystal structure (PDB: 6XAW) were set to a maximum distance of 25 Å and used as unambiguous interaction restraints. The FFI sequences of Ume6 595-660 and Pho23 147-280 were used as ambiguous interaction restraints. Multiple docking runs were performed, and the top-ranked clusters were selected based on HADDOCK scoring functions and the number of satisfied cross-links. To improve the accuracy of the final model, an iterative refinement protocol was employed. In each round, the best-scoring complex was re-docked using updated restraints based on prior cross-link evaluation. This process was repeated until no further improvement in cross-link satisfaction was observed. Final complex models were visualized and analysed using UCSF ChimeraX^70^. The video was generated using UCSF ChimeraX.

### Cloning of *SIN3* for systematic interaction mapping

As the *SIN3* open reading frame (ORF) was absent from the yeast ORFeome collection^47^, the 4.6 kb sequence was independently cloned using Gateway technology. The *SIN3* ORF was amplified from S288C genomic DNA (EMD4Biosciences) using Q5 high-fidelity polymerase (NEB) and primers FP391 and 392, which incorporate *attB* recombination sites. Purified amplicons were cloned into the pDONR223 entry vector using BP Clonase II Enzyme mix. Subsequently, the ORF was transferred into destination vectors using the LR Clonase II Enzyme mix. The integrity of all resulting constructs was verified through Oxford Nanopore long-read, whole plasmid sequencing (Plasmidsaurus).

### All-by-all pairwise test of full-length proteins

To map the Sin3L protein interaction network, we utilized Gal4 AD- and DB-ORF fusion clones, in Y8800 and Y8930 respectively, for all 12 subunits, sourced from the consolidated yeast ORFeome collection (except Sin3). An all-versus-all mating strategy was employed to generate a 12 x 12 interaction matrix. Additionally, all 12 DB-fusion proteins were tested against an empty AD vector to identify potential autoactivation, resulting in a final matrix of 156 Y2H tests. Media quality and reporter sensitivity were validated using six standardized control pairs (AY050 x AY051, AY052 x AY053, AY054 x AY055, AY056 x AY057, AY058 x AY059, AY060 x AY061) with established growth phenotypes^78^. Haploid clones were mated in liquid YPD at 30°C overnight. Diploids were subsequently selected in liquid SC-L-W for 48 hours. To assess protein-protein interactions, diploid cultures were spotted onto solid SC-L-W (mating control) and two selective media: SC-L-W-H supplemented with 1 mM 3AT and SC-L-W-A, corresponding to the *GAL1::HIS3* and *GAL2::ADE2* reporter genes, respectively. Plates were scored after a 4-day incubation at 30°C.

### Gene fragmentation

A series of truncation mutants was generated for all 12 ORFs coding for a protein of the Rpd3 complex using a two-round PCR approach. The first round of PCR consisted in amplifying fragmented derivatives of the 12 ORFs using primers harbouring universal adapter sequences, that are then recognized by a second set of primers with tails containing a sequence homologous to that of the insertion site of the expression plasmids pDEST-AD (FP387-FP388) and pDEST-DB (FP389-FP390) in a subsequent round of PCR. Gene fragments were cloned into both pDEST-AD and pDEST-DB using a gap repair strategy. Plasmids were linearized with the digestion enzymes Pst1 and Not1. Linearized expression plasmids pDEST-AD or pDEST-DB were transformed in the yeast strains HOY13 and HOY14, respectively, together with the ORF insert harbouring flanking regions homologous to that of the plasmid. All nucleotide coordinates are relative to the S288C reference genome deposited on the *Saccharomyces* Genome Database (SGD), except for those of Rxt2, for which we used the methionine codon at position -23 relative to the start codon reported on SGD. Primers used are presented in **Supplementary Tables 5 and 6**.

### Interaction mapping by primary Y2H screens

Overnight cultures of individual HOY14 yeast strains expressing a DB-ORF fusion protein (bait) were mated with 12 pools of HOY13 yeast cells expressing all AD-ORFs originating from one of the 12 genes (preys). In parallel, all yeast strains expressing a DB-ORF fusion protein were also mated with a strain harbouring a plasmid expressing the AD moiety only, referred to as empty AD control, to test for the auto-activator potential of the DB-ORFs. 6 pairs of haploid yeast strains (AY285 x AY291, AY286 x AY292, AY287 x AY293, AY288 x AY294, AY289 x AY295, AY290 x AY296) were used as controls to ensure the quality of the media. After overnight growth at 30°C in liquid YPD, mated yeast cells were transferred into liquid SC-L-W media to select for diploids. After overnight incubation at 30°C, diploid yeast cells were spotted onto the three following solid selection media: SC-L-W-H + 1mM 3AT, SC-L-W-A and SC-L-W-U. After 72 h incubation at 30°C, yeast colonies that grew on selective media were picked into liquid SC-L-W media and grown overnight. Cells were then lysed and both the AD-ORF and the DB-ORF were PCR amplified with indexed primers and sequenced via a PacBio platform to determine the identity of the respective bait and prey proteins.

Because each DB-ORF yeast strain was mated against a library of yeast strains expressing ∼100 different AD-ORFs in the first-pass screen, one bait protein could interact with more than one prey protein per pool. Multiple colonies per spot were hence picked into liquid SC-LW media. Yeast cultures were processed as previously described^78^ to generate lysates. A 3 μL aliquot of a 1:5 diluted lysate solution was used as a PCR template to produce DB-ORF and AD-ORF amplicons. To cost-effectively identify both bait and prey proteins for hundreds of positive hits, we used barcoded primers containing well-specific and plate-specific nucleotide sequences. Briefly, PCR was performed using well-indexed forward AD or DB primers together with plate-indexed reverse universal primers. PCR was conducted using the MyTaq HS polymerase (Bioline). All amplicons were pooled and sent to the Institute for Genome Sciences at the University of Maryland for sequencing using a PacBio Sequel II system. Identification of matching ORF pairs corresponding to Y2H-positive colonies was performed through demultiplexing sequencing data using the nucleotide barcodes. Similarly, as for other Y2H screens previously conducted by our group^79^, pairs identified in this primary screen were called First Pass Pairs (FiPPs).

### Interaction mapping by pairwise Y2H tests

Each FiPP identified through the primary screen was verified in a Y2H pairwise test. For all FiPPs, each yeast strain expressing a protein of interest was cherry-picked from the original collection of yeast transformants and mated with a yeast strain expressing its partner. Each individual yeast strain expressing a DB-ORF fusion protein was also mated with an empty AD control to test for *de novo* autoactivation. Cells were then spotted on SC-L-W to check for the presence of diploid cells, and the three Y2H selective media used for the primary screen. As positive and negative controls, 60 well-described benchmark interactions from a positive reference set (PRS) and 78 protein pairs from a random reference set (RRS) were included. These sets are referred to as hsPRSv2 and hsRRSv2^43^. Experiments were considered valid if most pairs detected by Choi et al were recovered while minimizing the number of RRS pairs detected. Each 96-well plate has a unique pattern of empty wells allowing the identification of any potential plate rotations or swaps and hence ensuring proper mating of the corresponding plates with each other (DB-X with AD-Y). We implemented the semi-automated pipeline described by Luck *et al.* for the scoring of growing yeast in 96-well format^79^ where each spot is attributed a growth score ranging from 0 to 4, corresponding to a negative readout to a strong readout, respectively. Two readings were performed by independent operators and scoring discrepancies were individually discussed. Both retest plates and autoactivation control plates were independently scored, and scores combined to produce a final score of each well. A pair was scored invalid if the test well or corresponding autoactivation control was contaminated, not spotted, etc. A pair was scored as *de novo* auto-activator if it had a similar or higher frequency of yeast colonies on the autoactivation plate compared to the retest plate. A pair was scored as positive if its growth score is equal to 2 or more on at least one selective media. Yeast cells from spots for which pairs were scored positive in the pairwise test were picked into liquid SC-L-W in a 96-well plate format. Identities of AD- and DB-ORFs were sequence confirmed as described for the first pass screen.

### Kinase substrate sensor assays

The 12 ORFs corresponding to full-length subunits of the Sin3L complex (see above) were PCR-amplified from yeast genomic DNA, and cloned into Gateway entry vectors, pDONR223. Following bacteria transformations, single colonies were picked, and the quality of cloning was checked for every ORF by bi-directional Sanger DNA sequencing. The 12 ORFs were introduced by LR clonase-mediated Gateway reactions (Life Technologies) into the KISS-C1 and N2-KISS destination plasmids to allow expression of the tested proteins, X and Y, as C-and N-terminal fusions (i.e. X-C_1_/N_2_-Y and Y-C_1_/N_2_-X). LR reaction products were subsequently transformed into *E. coli* DH5α competent cells and grown for 24 h on ampicillin-containing TFB medium. Plasmid DNA was extracted using a NucleoSpin 96 Plasmid kit from Macherey-Nagel. After PCR-amplification of the cloned ORFs from purified plasmid DNAs with plasmid-specific primers, the size of each DNA amplicon was confirmed by agarose gel electrophoresis. HEK293T cells, cultured as previously reported^43^, were transfected via a standard calcium phosphate transfection method with the purified bait, prey and reporter plasmids (KISS-C1 and N2-KISS) corresponding to empty controls (i.e. unfused gp130 tag or unfused TYK2 C-terminal fragment tag) or interacting proteins initially identified by Y2H. The KISS assay was then implemented as previously reported^43,80^. Luciferase activity was measured 48 h after transfection using the Luciferase Assay System kit (Promega) on a Enspire luminometer (Perkin-Elmer). The average of six independent culture wells was used. The published normalized luminescence ratio (NLR) cutoff of 4.04583, corresponding to 0% hsRRS-v2 pairs scored positive (FDR = 0%)^43^, was applied to validate Sin3L interactions. The luciferase signals obtained for each Sin3L bait-prey protein pair versus that obtained for the combination of the same bait with a negative control/irrelevant prey (unfused gp130), and versus that obtained for the combination of the same prey with a negative control/irrelevant bait (unfused TYK2 C-terminal fragment) were evaluated against the reference NLR cutoff corresponding to 0% hsRRS-v2 pair detection. NLR was calculated as follows, for each protein pair, X-Y:

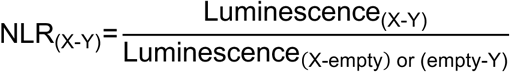

An interaction pair was scored positive when both NLR, using X-empty or empty-Y, exceeded the reference cutoff for either of the two configurations tested, corresponding to empirical p values = 0.01. Between the two NLR values of each positive pair and tested orientation, the smallest was used to construct **Extended Data Fig. 6c**.

### Protein purification for X-ray crystallography

For the yeast Sin3 PAH2-Ume6 SID proteins, a bicistronic expression construct (pET28: N-terminal, cleavable polyHis fusion) of yeast Sin3 (residues 402-473) and Ume6 (residues 500-543) was overexpressed in *E. coli* BL21 (DE3) in TB medium in the presence of 50 μg/mL of kanamycin. Cells were grown at 37°C to an OD of 0.8, cooled to 17°C, induced with 500 μM isopropyl-1-thio-D-galactopyranoside (IPTG), incubated overnight at 17°C, collected by centrifugation, and stored at -80°C. Cell pellets were lysed in buffer A (50 mM HEPES, pH 7.5, 300 mM NaCl, 10% glycerol, 20 mM Imidazole, and 7 mM β-mercaptoethanol) and the resulting lysate was centrifuged at 30,000 g for 40 min. Ni-NTA beads (Qiagen) were mixed with lysate supernatant for 30 min and washed with buffer A. Beads were transferred to an FPLC-compatible column and the bound protein was washed further with buffer A for 10 column volumes, and eluted with buffer B (50 mM HEPES pH 7.5, 300 mM NaCl, 10% glycerol, 300 mM Imidazole, and 7 mM BME). Human rhinovirus 3C protease was added to the eluted protein and incubated at 4°C overnight. The sample was concentrated and passed through a Superdex 200 10/300 column (GE Healthcare) in a buffer containing 20 mM HEPES, pH 7.5, 200 mM NaCl, 5% glycerol, and 1 mM TCEP. Fractions were pooled, concentrated to approximately 19 mg/mL and frozen at -80°C. For the free yeast Sin3 PAH2 protein, a construct of yeast MBP-Sin3 (residues 402-473 cloned into pMALX(E)^81^) was overexpressed in *E. coli* BL21 (DE3) in TB medium in the presence of 100 μg/mL of Ampicillin. Cells were grown at 37°C to an OD of 0.8, cooled to 17°C, induced with 500 μM IPTG, incubated overnight at 17°C, collected by centrifugation, and stored at -80°C. Cell pellets were lysed in buffer A (50 mM HEPES, pH 7.5, 500 mM NaCl, 10% glycerol, and 7 mM β-mercaptoethanol) and the resulting lysate was centrifuged at 30,000 g for 40 min. Amylose beads (NEB) were mixed with lysate supernatant for 1.5 h and washed with buffer A. Beads were transferred to an FPLC-compatible column and the bound protein was washed further with buffer A containing additional 1 M NaCl for 10 column volumes, and eluted with buffer B (25 mM HEPES pH 7.5, 200 mM NaCl, 5% glycerol, 20 mM maltose, and 7 mM BME). The eluted sample was concentrated and passed through a Superdex 200 16/600 column (GE Healthcare) in a buffer containing 20 mM HEPES, pH 7.5, 200 mM NaCl, 5% glycerol, 0.5 mM TCEP, and 1 mM DTT. Fractions were pooled, concentrated to approximately 20 mg/mL and frozen at -80°C.

### Crystallization of the yeast Sin3 PAH2 alone or in complex with Ume6 SID

With the aid of Formulatrix NT8, RockImager and ArtRobbins Phoenix liquid handlers, a sample of 400 μM Sin3 PAH2-Ume6 SID proteins and 5 μM trypsin were mixed and co-crystallized in an equivalent volume of 1.2 M NaCitrate and 0.1 M TrisHCl pH 8.0 by sitting-drop vapor diffusion at 20°C after three days. Similarly, a sample of 400 μM MBP-Sin3 PAH2 protein and 5 mM maltose was crystallized in (NH_4_)_2_SO_4_ and 0.1 M BisTris pH 6.0 by hanging-drop vapor diffusion at 20°C. Crystals were obtained using a combination of Formulatrix NT8 and ArtRobbins Phoenix liquid handlers and visualized using a Formulatrix RockImager. Large, single crystals were transferred into crystallization buffer containing 25% glycerol prior to flash-freezing in liquid nitrogen and shipped to the synchrotron for data collection.

### X-ray crystallography data collections and structure determinations

For the bicistronic yeast Sin3 PAH2-Ume6 SID (PDB: 6XAW), diffraction data from Sin3 PAH2-Ume6 SID complex crystals were collected at beamline 24ID-C of the NE-CAT at the Advanced Photon Source (Argonne National Laboratory). Data sets were integrated and scaled using XDS^82^. Structures were solved by Br-SAD using the programme Autosol in Phenix 1.13_2998 package^83^. Iterative manual model building and refinement using Phenix 1.13_2998 and Coot^84^ led to a model with excellent statistics. For the free yeast Sin3 PAH2 (PDB: 6XDJ), diffraction data from MBP-Sin3 PAH2 crystals were collected at beamline 24ID-E of the NE-CAT at the Advanced Photon Source (Argonne National Laboratory). Data sets were integrated and scaled using XDS^82^. Structures were solved by molecular replacement using the programme Phaser 2.8.3^85^ and the search model of MBP from PDB entry 4JBZ. Iterative manual model building and refinement using Phenix 1.16_3549^83^ and Coot^84^ led to a model with excellent statistics.

### Mutagenesis of yeast *SIN3*

The *SIN3* fragment 272-885 was mutagenized via error-prone PCR across 20 suboptimal conditions (**Supplementary Table 7**), using a recombinant Taq DNA polymerase (ThermoFisher Scientific), S288C genomic DNA (EMD4Biosciences), and primers AP434 and AP438. Successful amplicons were pooled and subjected to a second PCR round using primers with containing adapter tails (AP427 and AP267). The same fragment was PCR amplified using the High Fidelity Platinum Taq DNA Polymerase (ThermoFisher Scientific) to generate a wild-type control. The pDEST-DB expression vector was linearized with SrfI and NcoI restriction enzymes (NEB). Linearized vector and the *SIN3* insert were co-transformed in the HOY14 yeast strain. In parallel, transformation efficiency and plasmid background were monitored using undigested and linearized vector controls only, respectively. Yeast cells were plated onto solid SC-L media. To ensure library complexity, 15 independent gap-repair transformations were performed. Wild-type *UME6* fragment 295-726 cloned into pDEST-AD and transformed in HOY13 was directly picked from the yeast collection. We refer to this strain as AY280. AY280 was plated on solid SC-W media to generate a lawn of cells. All plates were incubated for three days at 30°C. Diploid libraries were generated by sequential replica-plating of HOY14 cells (expressing the *SIN3* mutant library or WT) and AY280 (AD-*UME6*) onto YPD media. To control for autoactivation, the library was crossed in parallel with a strain harbouring an empty AD vector. After overnight incubation, diploids were selected on SC-L-W media. To identify non-interfering alleles, diploids were replica-plated onto SC-L-W (pre-selection) and SC-L-W-A (post-selection) media. Following a 3-day incubation at 30°C, cells from the pre- and the post-selection plates were harvested and pooled in three independent batches representing distinct biological replicates for subsequent DNA extraction and analysis.

### Yeast DNA extraction for sequencing

Harvested yeast cells were lysed via a combined mechanical and chemical approach. Cell pellets were resuspended in 200 µL detergent lysis buffer (2% Triton X-100, 1% SDS, 100 mM NaCl, 10 mM Tris-Cl pH 8.0, and 1 mM EDTA), 200 µL phenol-chloroform-isoamyl alcohol, and ∼300 mg glass beads (Microspheres-Nanospheres). Samples were homogenized using a QIAGEN TissueLyser II (10 min, 30 Hz). Following addition of 200 µL TE buffer (pH 8.0) and centrifugation (16,300 x *g*, 5 min), the aqueous phase was collected. DNA was precipitated with 1 mL 100% ice-cold ethanol, resuspended in 400 µL TE buffer (pH 8.0), and subjected to a secondary purification step using 100 µL 4 M ammonium acetate and 1 mL ice-cold 100% ethanol precipitation. Resulting DNA pellets were air-dried for 30 minutes and resuspended in 50 µL TE. *SIN3* mutant libraries from pre- and post-selection pools were amplified using MyTaq HS polymerase (Bioline) using barcoded Y2H-DB and Y2H-term primers. To ensure robustness, three independent PCRs were performed per sample as technical replicates. Pooled amplicons were sequenced at the Molecular Biology Core Facilities (Dana-Farber Cancer Institute) on an Illumina NextSeq 500 platform (75 bp paired-end). To quantify variant representation, we calculated the fold-change (FC) in mutant abundance between post-selection and pre-selection pools. Enrichment and depletion values were normalized using the distribution of FCs from synonymous mutations to account for experimental variation. Normalized values were expressed as Z-scores. Significant mutant abundance deviations were defined using a threshold of |Z| >1.96 (p <0.05).

## Data and code availability

All raw data are provided with this paper (**Source Data**) and are available upon request. The mass spectrometry proteomics and crosslinking data have been deposited to the ProteomeXchange Consortium via the MassIVE partner repository with the dataset identifiers MSV000100035 and PXD071273. The cryo-EM structures of the Sin3-TAP, Sap30-TAP and Rxt3-TAP tagged Sin3L complexes are available under Protein Data Bank (PDB) accession codes 9TAG, 9TAF and 9TAH, respectively, and the associated cryo-EM maps deposited to the Electron Microscopy Data Bank (EMDB) under the accession codes EMD-55746, EMD-55745 and EMD-55747. The cryo-EM maps of the Sin3L complex with extended densities are available under EMDB accession codes EMD-55763, EMD-55762 and EMD-55764, for the Sin3-TAP, Sap30-TAP and Rxt3-TAP datasets, respectively. Models of the extended Sin3L structure, the extended Sin3L structure with Cti6 and the extended Sin3L structure with Cti6 and Ash1 are deposited at PDB-IHM and are available under PDB codes 9AAF, 9AAG and 9AAH, respectively. Co-complex Sin3 PAH2-Ume6 SID and free Sin3 PAH2 structural data have been deposited in PDB under the accession codes 6XAW and 6XDJ, respectively. This paper does not report original codes, and software used in the study is publicly available and referenced in the text.

## Statistical analyses

Except otherwise indicated, statistical analyses were performed on Prism (GraphPad) with each statistical test, definitions of centre (mean or median) and dispersion (standard error of the mean), and the number of samples per group (n, referring to number of independent replicates) indicated in the corresponding figures, figure legends and/or method details. Except otherwise indicated, calculations and normalizations of data were performed in Microsoft Excel 2011. Statistical significance was determined by p ≤ 0.05. Statistically significant comparisons in each figure are indicated with asterisks, *p ≤ 0.05; **p ≤ 0.01; ***p ≤ 0.001; ****p ≤ 0.0001.

## ACKNOWLEDGMENTS

We thank all members of the Das, Washburn and Vidal laboratories for helpful discussions. We thank Lars Pedersen (National Institutes of Health (NIH) National Institute of Environmental Health Sciences (NIEHS)) for providing the pMALX(E) plasmid, and Ezekiel Geffken, Zoe Yeoh, and Nick Vangos for help with purifications of the Sin3 and Ume6 proteins used for crystallography. This work was based upon research conducted at the Northeastern Collaborative Access Team beamlines, which were funded by the NIH National Institute of General Medical Sciences (NIGMS) (P30 GM124165). The Eiger 16M detector on the 24-ID-E beam line was funded by a NIH-ORIP HEI grant (S10OD021527). This research used resources of the Advanced Photon Source, a USA Department of Energy (DOE) Office of Science User Facility operated for the DOE Office of Science by Argonne National Laboratory under Contract No. DE-AC02-06CH11357. The Thermo Fisher 200 kV Cryo-TEM microscope Glacios equipped with autoloader and falcon 4i direct-electron detector with Selectris energy filter were funded by the Virology and Chemotherapy Division of the Rega Institute for Medical Research. All proteomics data were collected in the Mass Spectrometry and Proteomics Core facility utilizing the Orbitrap Ascend Tribrid System that was purchased with funds provided by the University of Kansas Cancer Center, which is supported by the National Cancer Institute Cancer Center Support Grant P30 CA168524. This work was supported by the Marie Skłodowska-Curie Actions (MSCA) European Postdoctoral Fellowship 101067095 from the European Research Executive Agency (J.O.), the Aspirant Fundamenteel Onderzoek grant 11PN624N from the Fonds Wetenschappelijk Onderzoek (FWO) (N.R.S.), the KU Leuven research grant C24E/22/035 (K.D.), the NIH NIGMS grants R01GM130885 (M.V.) and R01GM133185 (M.A.C., F.P.R. and M.V.), the NIH National Cancer Institute (NCI) grant R01CA266194 (M.V.), the Linde Family Foundation sponsored research agreement grants NIBR, 3DC, DDCF (S.D.P.), a Belgian American Educational Foundation Doctoral Research Fellowships (F.L.), a Wallonia-Brussels International (WBI)-World Excellence Fellowships (F.L.), a Fonds de la Recherche Scientifique (FRS-FNRS)-Télévie Grant (FC31747, Crédit n° 7459421F) (F.L., J.-C.T.), a Herman-van Beneden Prize (F.L.), a Josée and Jean Schmets Prize (F.L.), the Léon Fredericq Foundation (F.L., J.-C.T.), and a University of Liège mobility grants (F.L., Y.B.). Research reported in this publication was also supported by a Canada Excellence Research Chair (CERC) funding (K.D.), and the NIH NIGMS under award numbers T32GM138077 (J.C.) and R35GM145240 (M.P.W.). M.V. is a Chercheur Qualifié Honoraire and J.-C.T. a Directeur de Recherche from the FRS-FNRS, Wallonia-Brussels Federation, Belgium. The content of this work is solely the responsibility of the authors and does not necessarily represent the official views of the National Institutes of Health or other funding organizations.

## AUTHOR CONTRIBUTIONS

The project was conceived and supervised by J.O., M.V., M.P.W. and K.D. J.O., H.Y. and M.V. constructed the yeast strains. J.O. performed the yeast spot test assays and RT-qPCR experiments with help from F.L. and A.D.R. J.O. and N.R.S. purified and characterized the HDAC/Sin3L complexes for cryo-EM and XL-MS studies with help from J.V.D.V. N.R.S. collected and processed cryo-EM data with help from J.O. and K.D. N.R.S. modelled the final cryo-EM structures with feedback from J.O. and K.D. J.O., N.R.S. and K.D. analysed cryo-EM structures. J.C. collected, processed and analysed mass spectrometry data with feedback from M.P.W. J.N. developed the AlphaFold3-HADDOCK modelling pipeline to integrate XL-MS, cryo-EM, FFI and crystallographic data with feedback from J.O., N.R.S., J.C., M.P.W. and K.D. J.C. and J.N. generated and analysed the integrated models with feedback from J.O., N.R.S., M.P.W. and K.D. F.L., A.D.R., H.Y., Y.W. and K.S.-F. generated the protein fragment interaction dataset with feedback from J.O., T.H., M.A.C., J.-C.T., D.E.H. and M.V. F.L. and Y.B. selected and retested the minimal FFIs. F.L. and A.D.R. performed the mutagenesis studies with feedback from M.A.C., F.P.R., J.-C.T., D.E.H. and M.V. B.B. and O.D. analysed the Deletome dataset with help from J.O. and F.L. and feedback from M.V. I.L. conducted the KISS assays with help from J.O. and feedback from J.T. H.S.S. and S.D.P. conducted recombinant yeast protein expressions and purifications with help from J.O. and determined crystal structures. J.O. and N.R.S. constructed the figures and tables with help from J.C. and F.L. and feedback from M.V., M.P.W. and K.D. J.O. wrote the manuscript with help from N.R.S., J.C. and F.L. and feedback from M.V., M.P.W. and K.D. N.R.S. created the video with feedback from J.O., J.C., M.P.W. and K.D. All authors discussed and reviewed the results.

## ADDITIONAL INFORMATION

Supplementary Information is available for this paper. Correspondence and requests for materials should be addressed to Kalyan Das.

## EXTENDED DATA

**Extended Data Fig. 1.**
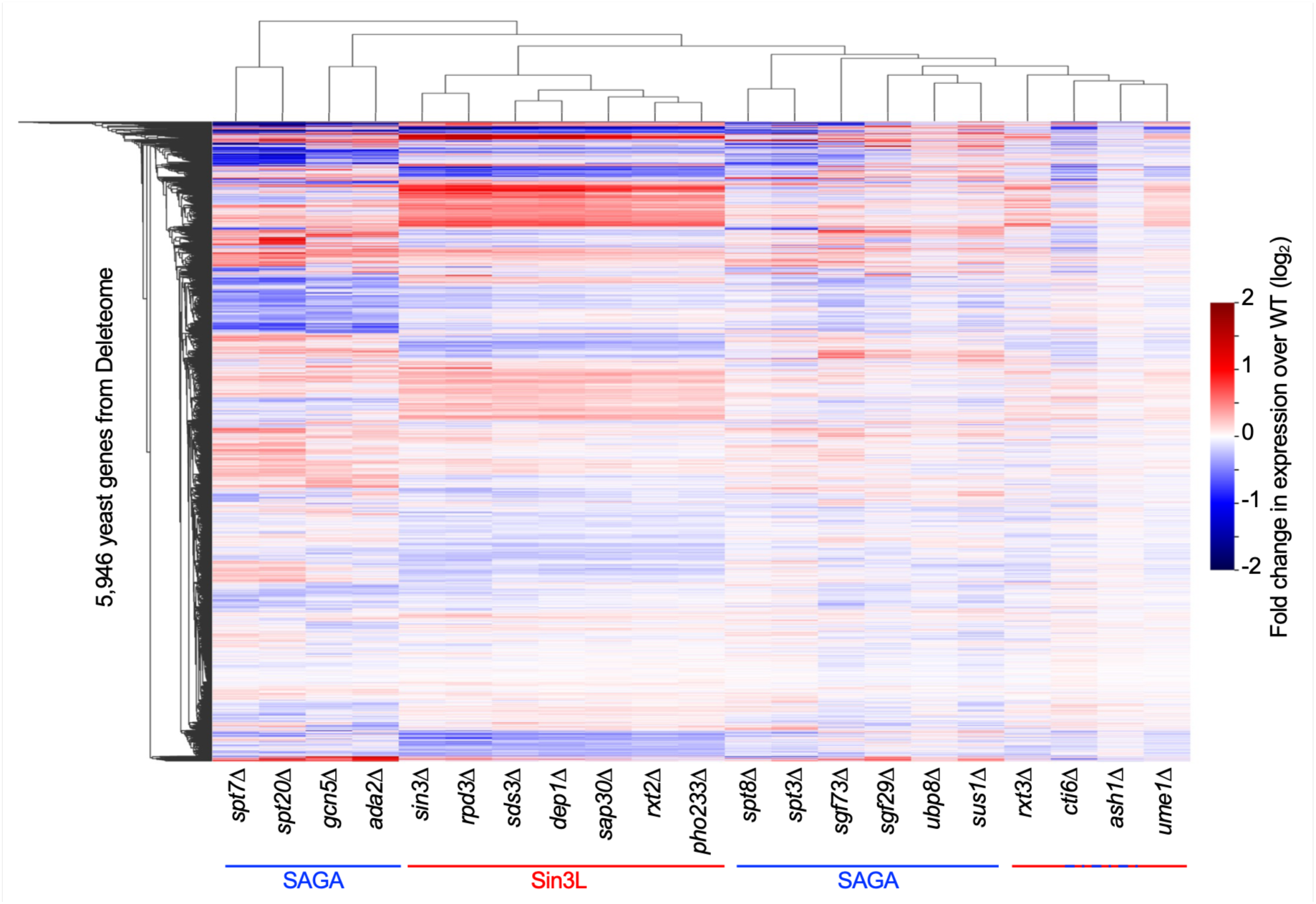
| Hierarchical clustering of gene expression changes in Sin3L and SAGA deletions. Hierarchical clustering heatmap of gene expression profiles extracted from the Deleteome compendium^36^. Dendrograms along the x- and y-axes represent clustering of deletion strains and genes, respectively.

**Extended Data Fig. 2.**
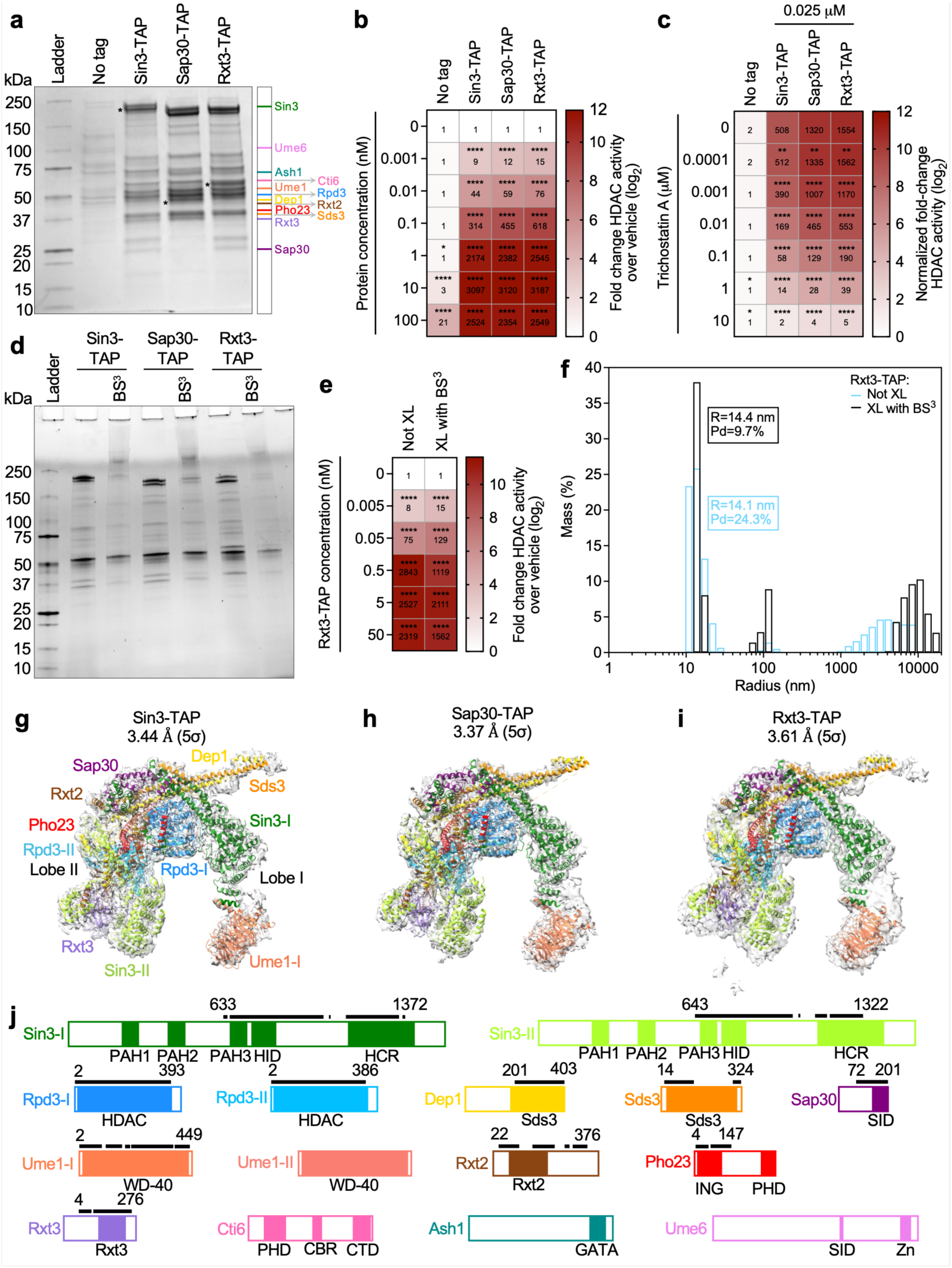
| Biophysical characterization of native Sin3L complexes. **a**, Coomassie-stained SDS-PAGE of Sin3L purified via Sin3-TAP, Sap30-TAP or Rxt3-TAP, with a no-tag control. Canonical subunits are indicated by coloured lines as per their molecular weights; asterisks denote the TAP-tagged bait (21 kDa). **b**, Heatmap of deacetylase activity for purified complexes (n=3 biologically independent samples per group). Values represent mean activity (linear scale; colour scale in log_2_). Residual activity in the no-tag control likely reflects residual HDAC enzymes. **c**, Deacetylase activity of the purified complexes following inhibition with Trichostatin A. **d**, Unstained SDS-PAGE of Sap30-TAP and Rxt3-TAP complexes following BS^3^ crosslinking. **e**, Deacetylase activity of BS^3^-crosslinked Sin3L complex purified via Rxt3-TAP (n=3 biologically independent samples per group). **f**, Dynamic light scattering profiles of non-crosslinked and crosslinked (XL) samples, showing hydrodynamic radius and mass distribution. The polydispersity (Pd) of the samples is also indicated. **g–i**, Core Sin3L structures purified via Sin3-TAP (**g**), Sap30-TAP (**h**) or Rxt3-TAP (**i**) fitted into sharpened cryo-EM maps (contour level of 5σ). **j**, Domain map of Sin3L subunits, with regions resolved in the cryo-EM core structures indicated by black lines. For gel source data, see Supplementary Fig. 1.

**Extended Data Fig. 3.**
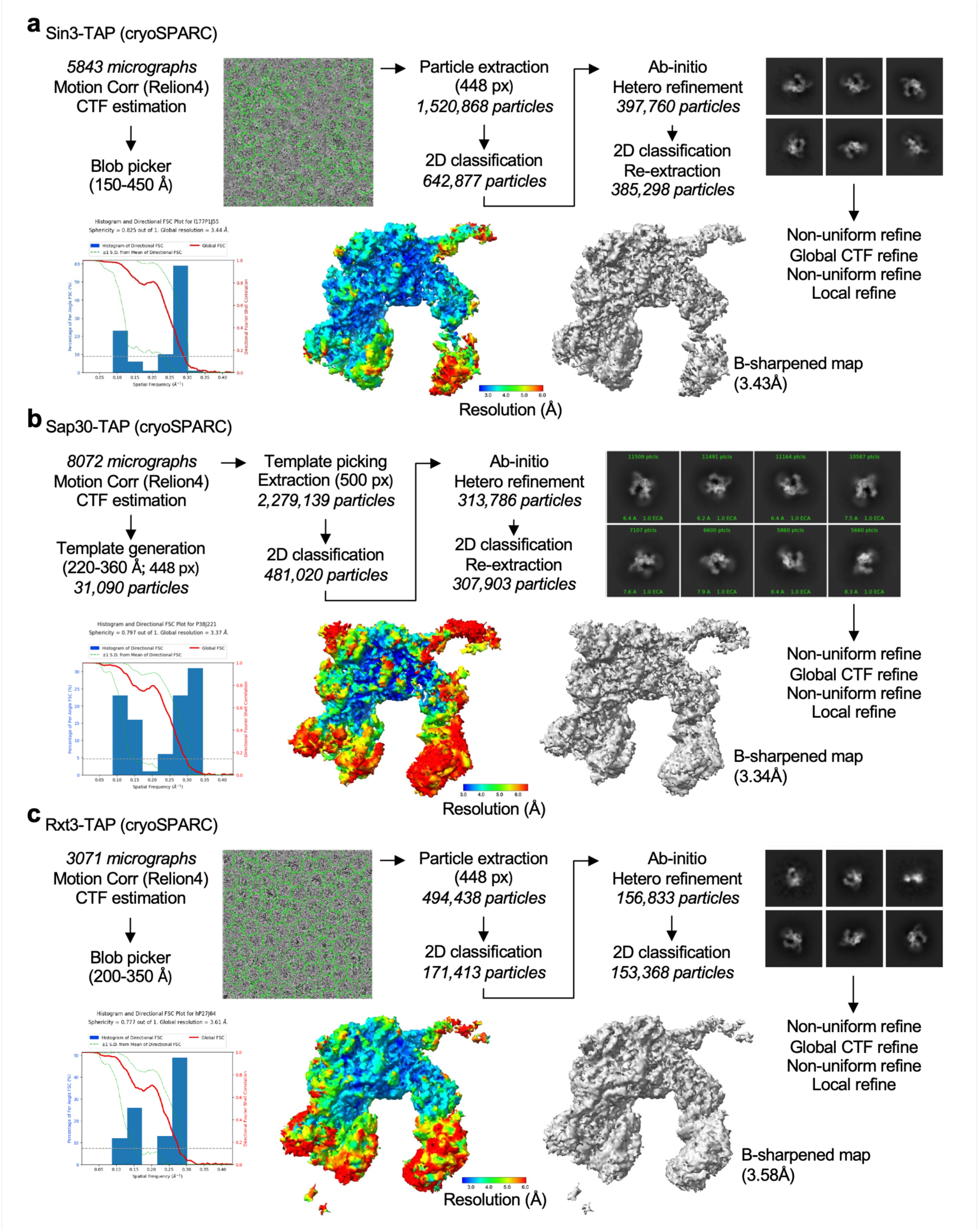
| Single-particle cryo-EM data processing workflows. **a-c**, Processing pipelines for Sin3-TAP (**a**), Sap30-TAP (**b**) and Rxt3-TAP (**c**) datasets. Movies were motion-corrected in RELION and further processed in cryoSPARC. Fourier shell correlation curves were calculated using the 3DFSC webserver.

**Extended Data Fig. 4.**
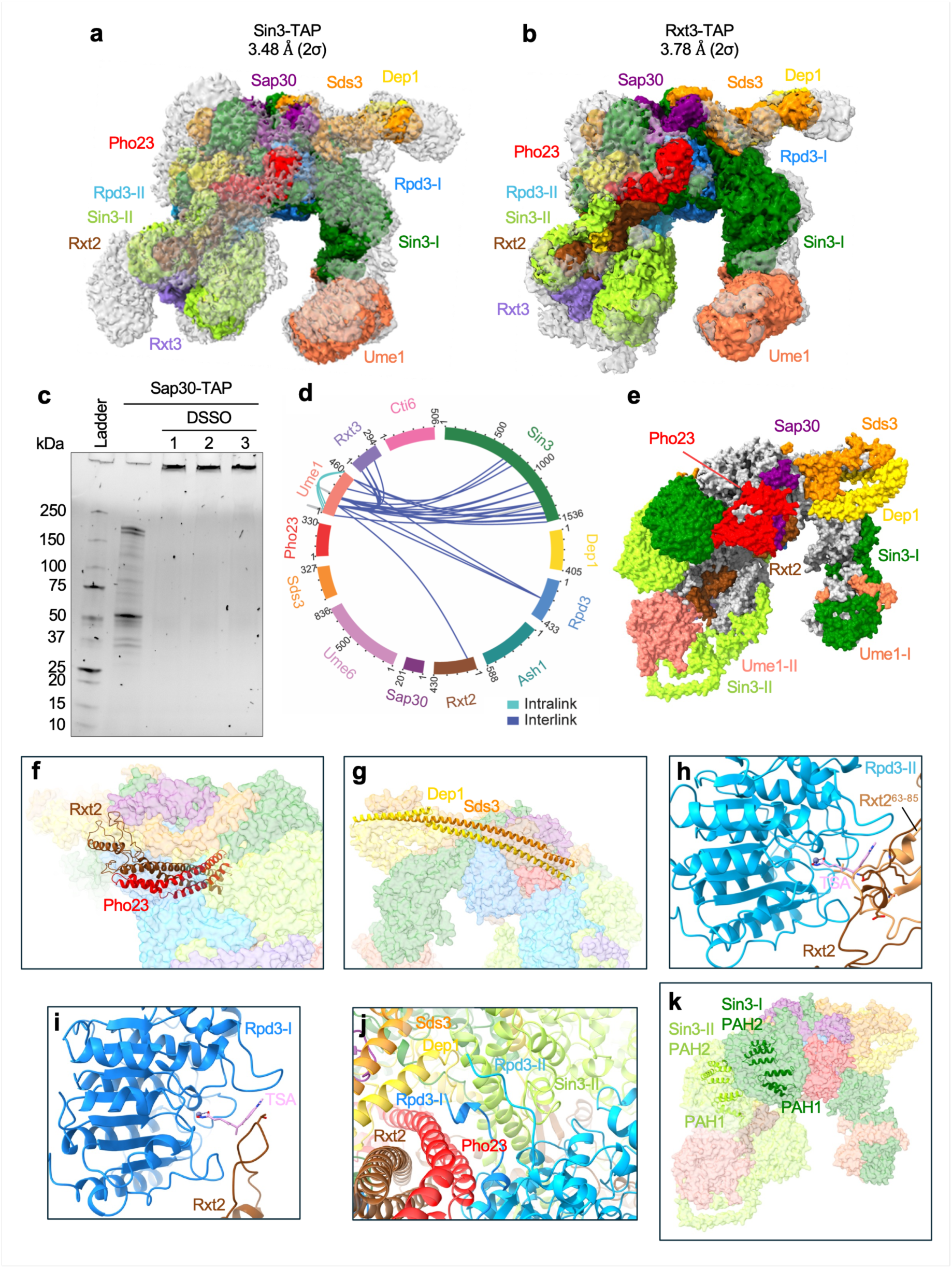
| Extended structural features of the Sin3L complex. **a**,**b**, Unsharpened cryo-EM maps of Sin3-TAP (**a**) and Rxt3-TAP (**b**) purified complexes (contour level of 2σ), showing modelled core regions in colour and unassigned, flexible densities in grey. **c**, Unstained SDS-PAGE of DSSO-crosslinked Sap30-TAP samples used for XL-MS (n=3 biologically independent samples per group). **d**, Circular map of Ume1 crosslinks; crosslinks consistent with the core structure are shown in grey, and unmatched crosslinks (Cα-Cα >37 Å) in colour. **e**, Surface representation of the extended Sin3L model, highlighting integrative modelling additions (colours) relative to the core cryo-EM structure (grey). **f**,**g**, Coiled-coil assemblies of Rxt2-Pho23 (**f**) and Sds3-Dep1 (**g**) in the extended Sin3L model. **h**,**i,** Occlusion of the Rpd3-II (**h**) and Rpd3-I (**i**) active sites by Rxt2 in the core and extended structures. For comparison, Trichostatin A-bound human HDAC8 (PDB: 1T64) is shown in the active site. **j**, Dimerization of the Rpd3 C-termini in the extended structure. **k**, Sin3 paired amphipathic helix domains (PAH1 and PAH2) from the extended Sin3L assembly. For gel source data, see Supplementary Fig. 1.

**Extended Data Fig. 5.**
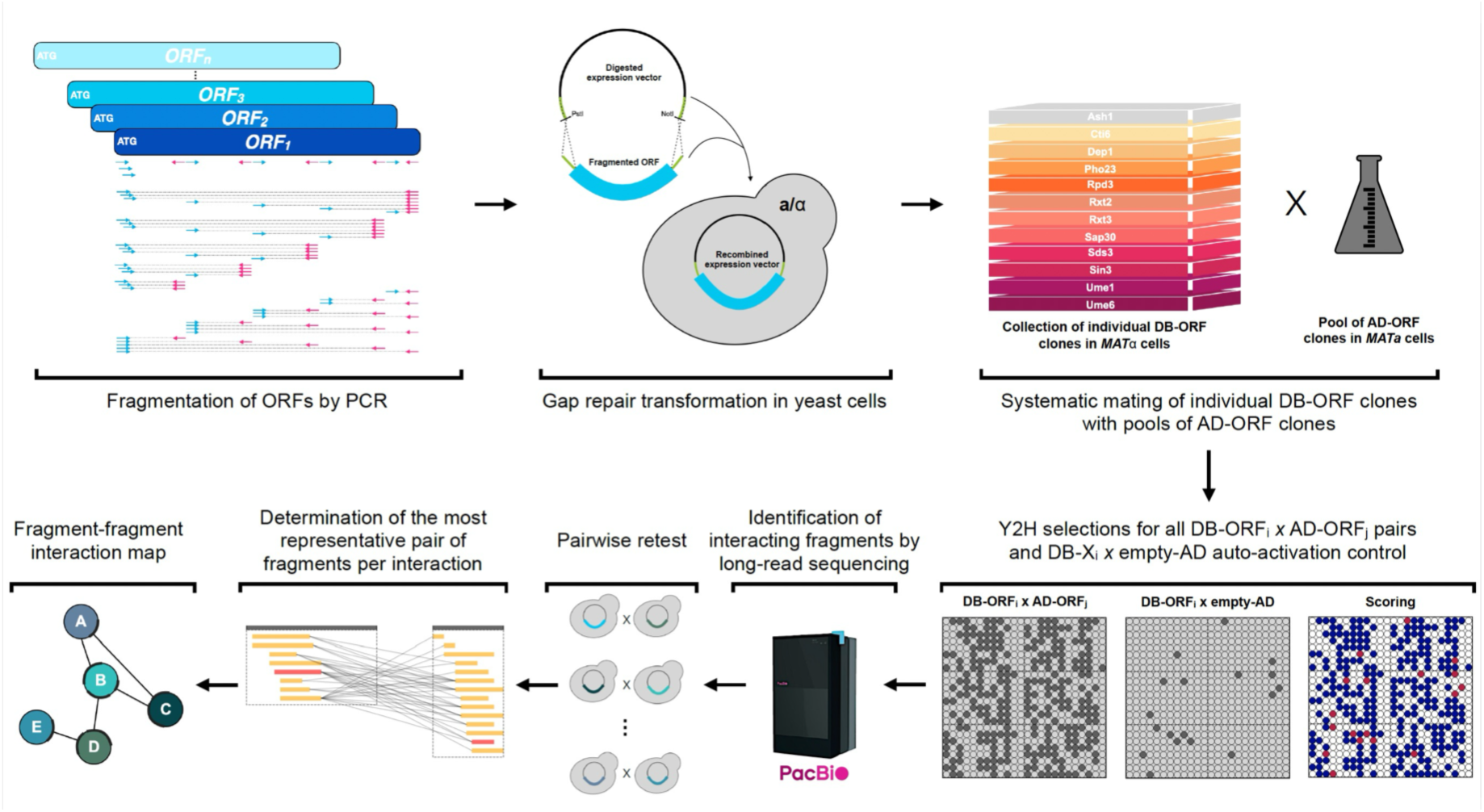
| Pipeline for the fragment-resolved interactome mapping of Sin3L. Schematic of the PCR-based fragmentation approach coupled to yeast gap-repair cloning and high-throughput sequencing for systematic fragment-fragment interaction mapping.

**Extended Data Fig. 6.**
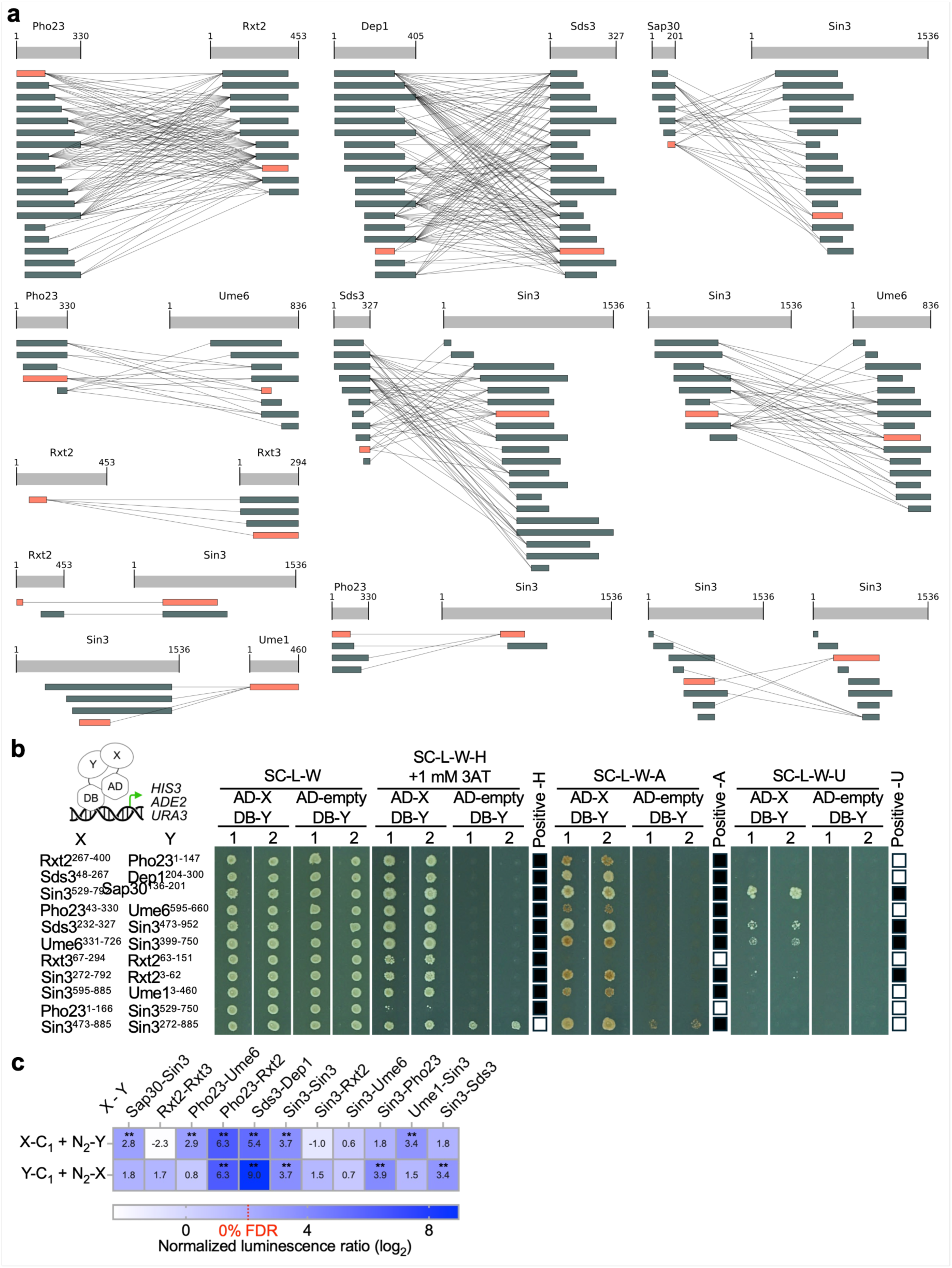
| Subunit fragment-resolved Sin3L interactome. **a**, Fragment–fragment interactions identified between Sin3L subunits; minimal interacting regions are highlighted in orange. **b**, Yeast two-hybrid validation of selected interactions using multiple reporter genes. Tested fragments fused to Gal4 activation (AD) or DNA-binding (DB) domains are indicated. Biological replicates are labelled 1 and 2. Growth conditions for each reporter and 3AT concentration are specified. Positive interactions are summarized by black squares in the different columns. Pictures were taken after five days of incubation at 30°C. **c**, Orthogonal validation of interactions using full-length proteins in the mammalian KISS assay at 0% false discovery rate (FDR).

**Extended Data Fig. 7.**
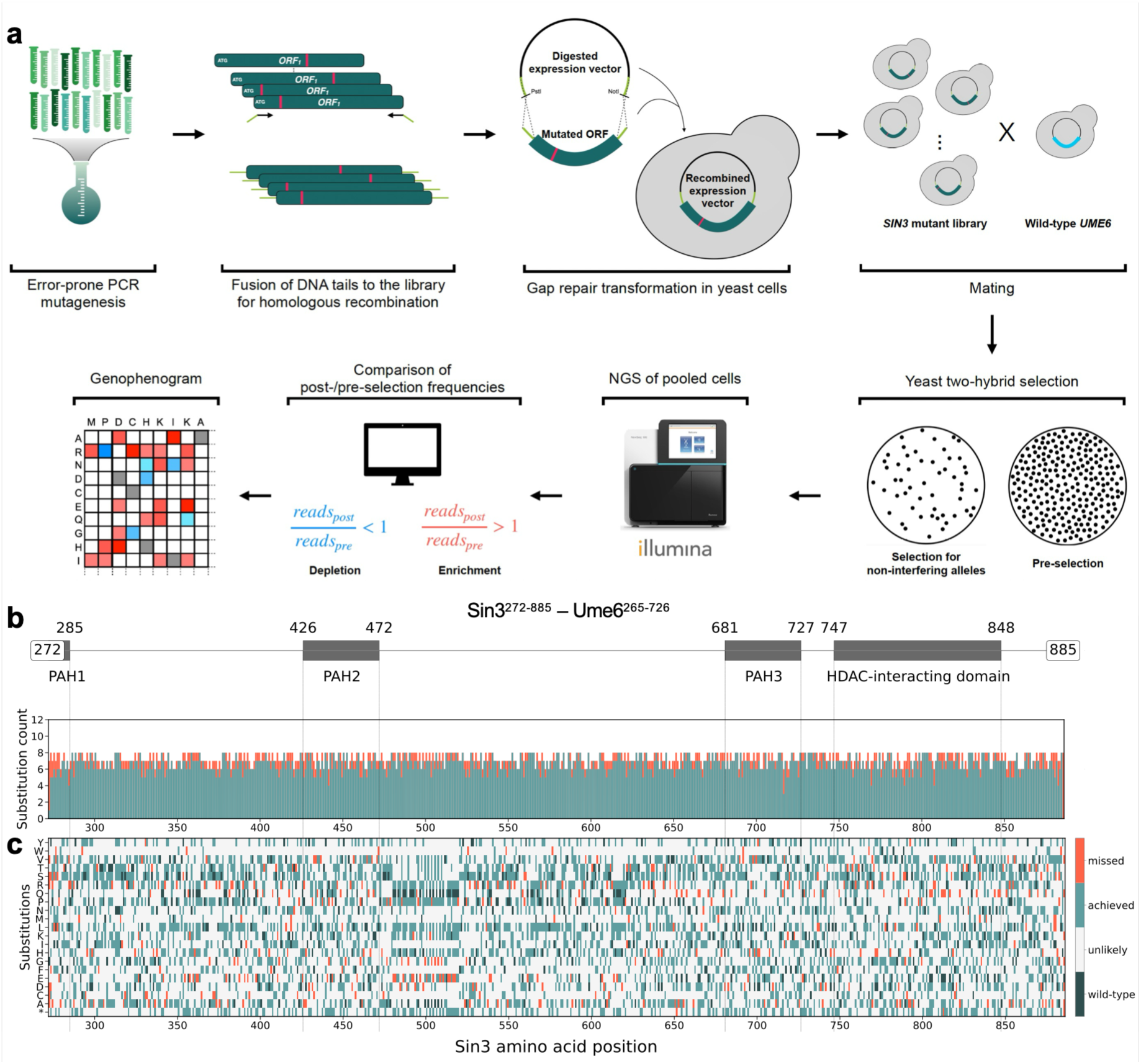
| Mutational scanning of the Sin3-Ume6 interface. **a,** Workflow for high-throughput mutagenesis screening. **b**, Absolute library coverage per residue for the Sin3 (272-885)-Ume6 (265-726) interaction. **c**, Detailed coverage map showing wild-type residues (dark green), observed single-nucleotide substitutions (light green), missing single-nucleotide substitutions (red), and multi-nucleotide substitutions (grey).

**Extended Data Fig. 8.**
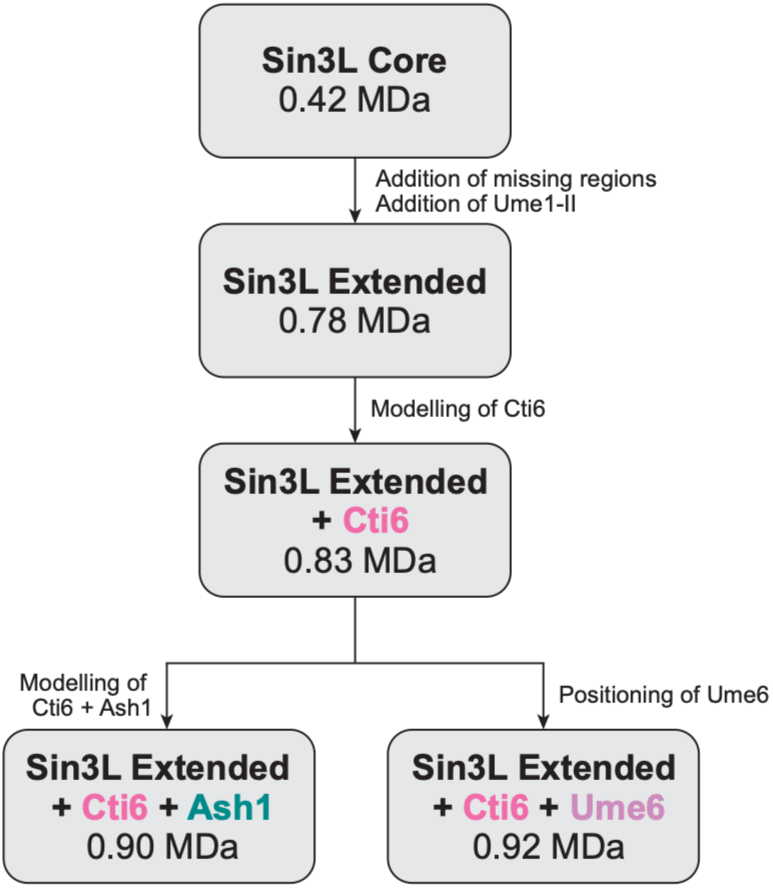
| Stepwise integrative process used to expand Sin3L structures. Schematic of the Sin3L core and extended structures determined using the integrated structural dynamics platform.

**Extended Data Table 1.** | Single-particle cryo-EM data collection, refinement and validation statistics.

|  | Sin3-TAP | Sap30-TAP | Rxt3-TAP |
| --- | --- | --- | --- |
| PDB ID | 9TAG | 9TAF | 9TAH |
| Data collection |  |  |  |
| Grid type | Quantifoil R1.2/1.3 (2 nm C) | Quantifoil R1.2/1.3 (2 nm C) | Quantifoil R1.2/1.3 (2 nm C) |
| Microscope | TFS Glacios | TFS Glacios | TFS Glacios |
| Detector | Falcon IV | Falcon IV | Falcon IV |
| Voltage (kV) | 200 | 200 | 200 |
| Electron exposure (e-/Å²) | 40 | 40 | 40 |
| Magnification | 100,000x | 100,000x | 100,000x |
| Pixel size (Å) | 1.16 | 1.16 | 1.16 |
| Defocus range (µm) | -0.6 to -1.8 | -0.6 to -1.8 | -0.6 to -1.8 |
| Data processing with cryoSPARC (Sin3L core) |  |  |  |
| EMDB ID | EMD-55746 | EMD-55745 | EMD-55747 |
| Number of micrographs | 5843 | 8072 | 3071 |
| Symmetry imposed | C1 | C1 | C1 |
| Number of initial particles | 1,520,868 | 2,279,139 | 494,438 |
| Number of final particles | 385,298 | 307,903 | 171,413 |
| Map resolution (Å) | 3.43 | 3.34 | 3.58 |
| 3DFSC map resolution (Å) | 3.44 | 3.37 | 3.61 |
| FSC threshold | 0.143 | 0.143 | 0.143 |
| Map resolution range (Å) | 2.71 – 8.11 | 2.72 – 7.75 | 2.89 – 7.93 |
| Data processing with Relion (Sin3L extended densities) |  |  |  |
| EMDB ID | E=[^?MD-55763 | EMD-55762 | EMD-55764 |
| Number of micrographs | 5880 | 6726 | 3071 |
| Symmetry imposed | C1 | C1 | C1 |
| Initial particle stack (no.) | 783,704 | 956,595 | 470,315 |
| Final particle stack (no.) | 276,299 | 164,944 | 80,030 |
| Map resolution (Å) | 3.43 | 3.81 | 3.77 |
| 3DFSC map resolution (Å) | 3.48 | 3.78 | 3.78 |
| FSC threshold | 0.143 | 0.143 | 0.143 |
| Model refinement |  |  |  |
| Initial model used (PDB) | 8HPO | 8HPO | 8HPO |
| Phenix model resolution (Å) | 3.26 | 3.14 | 3.36 |
| FSC threshold | 0.143 | 0.143 | 0.143 |
| Map/model correlation | 0.75 | 0.71 | 0.68 |
| Molprobit score | 2.27 | 2.19 | 2.40 |
| Non-hydrogen atoms | 27,036 | 27,036 | 27,036 |
| Protein residues | 3,508 | 3,508 | 3,508 |
| Clash score | 9.76 | 9.70 | 9.70 |
| B-factors |  |  |  |
| Protein | 110.89 | 108.08 | 119.90 |
| Ligand | 74.19 | 66.87 | 93.93 |
| R.m.s. deviations |  |  |  |
| Bond legnth (Å) | 0.003 | 0.003 | 0.003 |
| Bond angles (°) | 0.580 | 0.587 | 0.599 |
| Poor rotamers (%) | 3.54 | 3.33 | 3.43 |
| Ramachandran plot |  |  |  |
| Favored (%) | 95.09 | 94.72 | 91.97 |
| Allowed (%) | 4.85 | 5.11 | 7.85 |
| Not allowed (%) | 0.06 | 0.18 | 0.18 |

**Extended Data Table 2.**
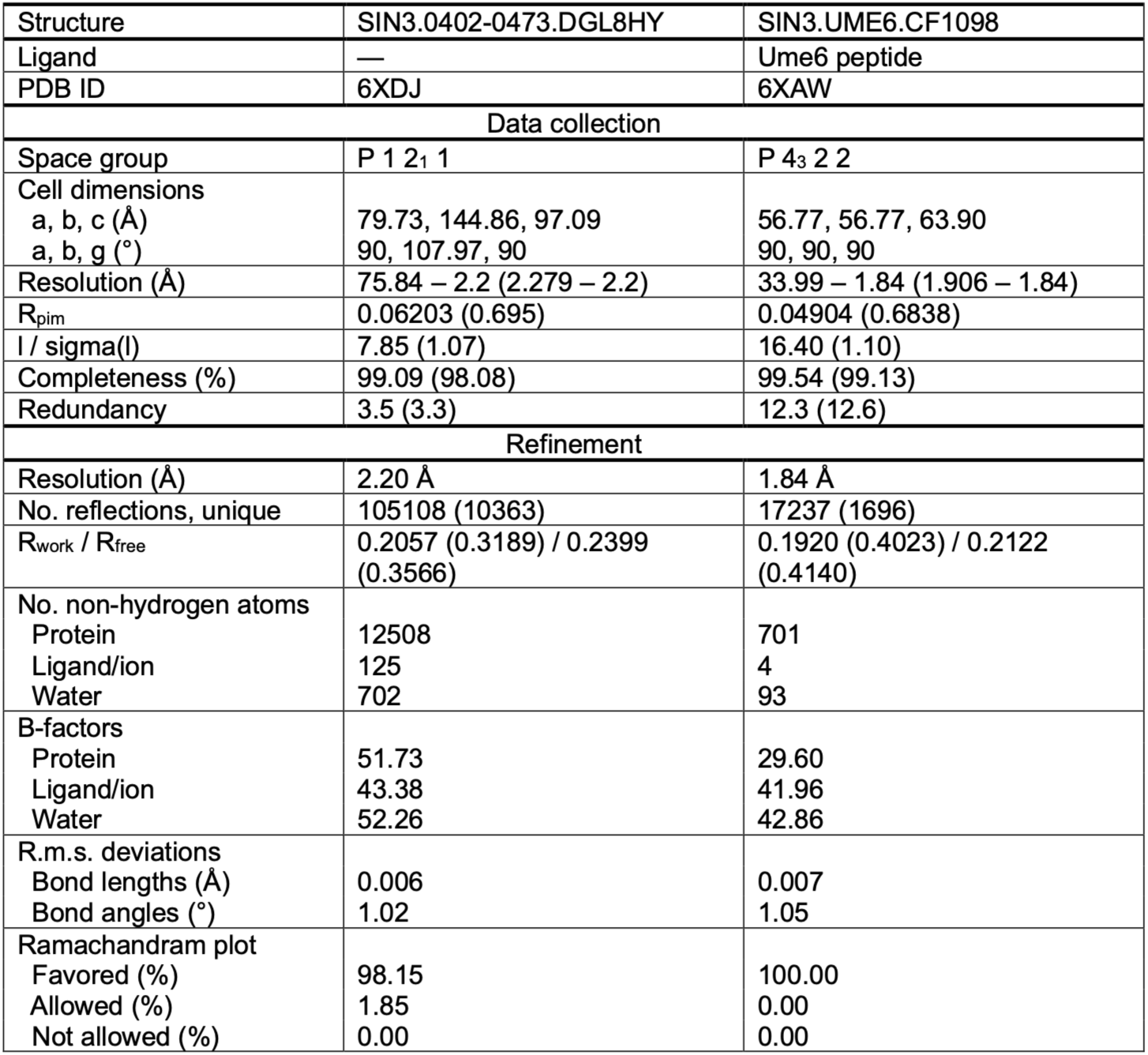
| Crystallographic data for the free Sin3 PAH2 domain and the Sin3 PAH2-Ume6 SID structures.

## MAIN REFERENCES

1 Voss, T. C. & Hager, G. L. Dynamic regulation of transcriptional states by chromatin and transcription factors. Nat Rev Genet 15, 69–81 (2014). 10.1038/nrg3623

2 Peserico, A. & Simone, C. Physical and functional HAT/HDAC interplay regulates protein acetylation balance. J Biomed Biotechnol 2011, 371832 (2011). 10.1155/2011/371832

3 Talukdar, P. D. & Chatterji, U. Transcriptional co-activators: emerging roles in signaling pathways and potential therapeutic targets for diseases. Signal Transduct Target Ther 8, 427 (2023). 10.1038/s41392-023-01651-w

4 He, J. et al. Dual-role transcription factors stabilize intermediate expression levels. Cell 187, 2746–2766 e2725 (2024). 10.1016/j.cell.2024.03.023

5 Adams, G. E., Chandru, A. & Cowley, S. M. Co-repressor, co-activator and general transcription factor: the many faces of the Sin3 histone deacetylase (HDAC) complex. Biochem J 475, 3921–3932 (2018). 10.1042/BCJ20170314

6 Talbert, P. B., Meers, M. P. & Henikoff, S. Old cogs, new tricks: the evolution of gene expression in a chromatin context. Nat Rev Genet 20, 283–297 (2019). 10.1038/s41576-019-0105-7

7 Carter, B., Ku, W. L., Pelt, J. & Zhao, K. Concurrent mapping of multiple epigenetic marks and co-occupancy using ACT2-seq. Cell Biosci 11, 198 (2021). 10.1186/s13578-021-00711-4

8 Kan, R. L., Chen, J. & Sallam, T. Crosstalk between epitranscriptomic and epigenetic mechanisms in gene regulation. Trends Genet 38, 182–193 (2022). 10.1016/j.tig.2021.06.014

9 Narlikar, G. J., Fan, H. Y. & Kingston, R. E. Cooperation between complexes that regulate chromatin structure and transcription. Cell 108, 475–487 (2002). 10.1016/s0092-8674(02)00654-2

10 Patel, A. B., He, Y. & Radhakrishnan, I. Histone acetylation and deacetylation - Mechanistic insights from structural biology. Gene 890, 147798 (2024). 10.1016/j.gene.2023.147798

11 Adams, M. K. et al. Differential Complex Formation via Paralogs in the Human Sin3 Protein Interaction Network. Mol Cell Proteomics 19, 1468–1484 (2020). 10.1074/mcp.RA120.002078

12 Banks, C. A. S. et al. Integrative Modeling of a Sin3/HDAC Complex Sub-structure. Cell Rep 31, 107516 (2020). 10.1016/j.celrep.2020.03.080

13 Vidal, M. & Gaber, R. F. RPD3 encodes a second factor required to achieve maximum positive and negative transcriptional states in Saccharomyces cerevisiae. Mol Cell Biol 11, 6317–6327 (1991). 10.1128/mcb.11.12.6317-6327.1991

14 Taunton, J., Hassig, C. A. & Schreiber, S. L. A mammalian histone deacetylase related to the yeast transcriptional regulator Rpd3p. Science 272, 408–411 (1996). 10.1126/science.272.5260.408

15 Vidal, M. Playing Hide-and-Seek with Yeast. Cell 166, 1069–1073 (2016). 10.1016/j.cell.2016.08.018

16 Grzenda, A., Lomberk, G., Zhang, J. S. & Urrutia, R. Sin3: master scaffold and transcriptional corepressor. Biochim Biophys Acta 1789, 443–450 (2009). 10.1016/j.bbagrm.2009.05.007

17 Carrozza, M. J. et al. Stable incorporation of sequence specific repressors Ash1 and Ume6 into the Rpd3L complex. Biochim Biophys Acta 1731, 77–87; discussion 75-76 (2005). 10.1016/j.bbaexp.2005.09.005

18 Sardiu, M. E. et al. Determining protein complex connectivity using a probabilistic deletion network derived from quantitative proteomics. PLoS One 4, e7310 (2009). 10.1371/journal.pone.0007310

19 Carrozza, M. J. et al. Histone H3 methylation by Set2 directs deacetylation of coding regions by Rpd3S to suppress spurious intragenic transcription. Cell 123, 581–592 (2005). 10.1016/j.cell.2005.10.023

20 Keogh, M. C. et al. Cotranscriptional set2 methylation of histone H3 lysine 36 recruits a repressive Rpd3 complex. Cell 123, 593–605 (2005). 10.1016/j.cell.2005.10.025

21 Guo, Z. et al. Structure of a SIN3-HDAC complex from budding yeast. Nat Struct Mol Biol 30, 753–760 (2023). 10.1038/s41594-023-00975-z

22 Patel, A. B. et al. Cryo-EM structure of the Saccharomyces cerevisiae Rpd3L histone deacetylase complex. Nat Commun 14, 3061 (2023). 10.1038/s41467-023-38687-z

23 Papamichos-Chronakis, M., Petrakis, T., Ktistaki, E., Topalidou, I. & Tzamarias, D. Cti6, a PHD domain protein, bridges the Cyc8-Tup1 corepressor and the SAGA coactivator to overcome repression at GAL1. Mol Cell 9, 1297–1305 (2002). 10.1016/s1097-2765(02)00545-2

24 Puig, S., Lau, M. & Thiele, D. J. Cti6 is an Rpd3-Sin3 histone deacetylase-associated protein required for growth under iron-limiting conditions in Saccharomyces cerevisiae. J Biol Chem 279, 30298–30306 (2004). 10.1074/jbc.M313463200

25 Aref, R. & Schuller, H. J. Functional analysis of Cti6 core domain responsible for recruitment of epigenetic regulators Sin3, Cyc8 and Tup1. Curr Genet 66, 1191–1203 (2020). 10.1007/s00294-020-01109-4

26 Lettow, J., Kliewe, F., Aref, R. & Schuller, H. J. Functional characterization and comparative analysis of gene repression-mediating domains interacting with yeast pleiotropic corepressors Sin3, Cyc8 and Tup1. Curr Genet 69, 127–139 (2023). 10.1007/s00294-023-01262-6

27 Dai, W. et al. PKA plays a conserved role in regulating gene expression and metabolic adaptation by phosphorylating Rpd3/HDAC1. Nat Commun 16, 4030 (2025). 10.1038/s41467-025-59064-y

28 Parnell, E. J., Parnell, T. J., Yan, C., Bai, L. & Stillman, D. J. Ash1 and Tup1 dependent repression of the Saccharomyces cerevisiae HO promoter requires activator-dependent nucleosome eviction. PLoS Genet 16, e1009133 (2020). 10.1371/journal.pgen.1009133

29 Kadosh, D. & Struhl, K. Repression by Ume6 involves recruitment of a complex containing Sin3 corepressor and Rpd3 histone deacetylase to target promoters. Cell 89, 365–371 (1997). 10.1016/s0092-8674(00)80217-2

30 Yamada, Y. RPD3 and UME6 are involved in the activation of PDR5 transcription and pleiotropic drug resistance in rho(0) cells of Saccharomyces cerevisiae. BMC Microbiol 21, 311 (2021). 10.1186/s12866-021-02373-1

31 Harris, A. & Unal, E. The transcriptional regulator Ume6 is a major driver of early gene expression during gametogenesis. Genetics 225 (2023). 10.1093/genetics/iyad123

32 Raithatha, S. A., Vaza, S., Islam, M. T., Greenwood, B. & Stuart, D. T. Ume6 Acts as a Stable Platform To Coordinate Repression and Activation of Early Meiosis-Specific Genes in Saccharomyces cerevisiae. Mol Cell Biol 41, e0037820 (2021). 10.1128/MCB.00378-20

33 Vidal, M., Brachmann, R. K., Fattaey, A., Harlow, E. & Boeke, J. D. Reverse two-hybrid and one-hybrid systems to detect dissociation of protein-protein and DNA-protein interactions. Proc Natl Acad Sci U S A 93, 10315–10320 (1996). 10.1073/pnas.93.19.10315

34 Maxon, M. E. & Herskowitz, I. Ash1p is a site-specific DNA-binding protein that actively represses transcription. Proc Natl Acad Sci U S A 98, 1495–1500 (2001). 10.1073/pnas.98.4.1495

35 Vidal, M., Buckley, A. M., Hilger, F. & Gaber, R. F. Direct selection for mutants with increased K+ transport in Saccharomyces cerevisiae. Genetics 125, 313–320 (1990). 10.1093/genetics/125.2.313

36 Kemmeren, P. et al. Large-scale genetic perturbations reveal regulatory networks and an abundance of gene-specific repressors. Cell 157, 740–752 (2014). 10.1016/j.cell.2014.02.054

37 Dominguez, C., Boelens, R. & Bonvin, A. M. HADDOCK: a protein-protein docking approach based on biochemical or biophysical information. J Am Chem Soc 125, 1731–1737 (2003). 10.1021/ja026939x

38 van Zundert, G. C. P. et al. The HADDOCK2.2 Web Server: User-Friendly Integrative Modeling of Biomolecular Complexes. J Mol Biol 428, 720–725 (2016). 10.1016/j.jmb.2015.09.014

39 Abramson, J. et al. Accurate structure prediction of biomolecular interactions with AlphaFold 3. Nature 630, 493–500 (2024). 10.1038/s41586-024-07487-w

40 Kao, A. et al. Development of a novel cross-linking strategy for fast and accurate identification of cross-linked peptides of protein complexes. Mol Cell Proteomics 10, M110 002212 (2011). 10.1074/mcp.M110.002212

41 Liu, F., Lossl, P., Scheltema, R., Viner, R. & Heck, A. J. R. Optimized fragmentation schemes and data analysis strategies for proteome-wide cross-link identification. Nat Commun 8, 15473 (2017). 10.1038/ncomms15473

42 Liu, X. et al. Driving integrative structural modeling with serial capture affinity purification. Proc Natl Acad Sci U S A 117, 31861–31870 (2020). 10.1073/pnas.2007931117

43 Choi, S. G. et al. Maximizing binary interactome mapping with a minimal number of assays. Nat Commun 10, 3907 (2019). 10.1038/s41467-019-11809-2

44 He, Y. & Radhakrishnan, I. Solution NMR studies of apo-mSin3A and -mSin3B reveal that the PAH1 and PAH2 domains are structurally independent. Protein Sci 17, 171–175 (2008). 10.1110/ps.073097308

45 Kumar, G. S., Xie, T., Zhang, Y. & Radhakrishnan, I. Solution structure of the mSin3A PAH2-Pf1 SID1 complex: a Mad1/Mxd1-like interaction disrupted by MRG15 in the Rpd3S/Sin3S complex. J Mol Biol 408, 987–1000 (2011). 10.1016/j.jmb.2011.03.043

46 Zhao, H. L., H.; Wang, C.; Yang, X.; Li, H.; Zou, B.; Dong, S.; Zhang, N.; Zhou, Y.; Yi, L.; Zhang, Y.; Xie, Y.; Qin, D.; Chao, W. C .H.; Pei, D.; He, J. Chromatin context-dependent deacetylation by the asymmetric Rpd3L (bioRxiv, 2026). 10.64898/2026.01.08.698523

47 Lambourne, L. Y., A.; Wang, Y.; Desbuleux, A.; Kim, D.-K.; Cafarelli, T.; Pons, C.; Kovács, I. A.; Jailkhani, N.; Schlabach, S.; De Ridder, D.; Luck, K.; Bian, W.; Shen, Y.; Yang, Z.; Mee, M. W.; Helmy, M.; Jacob, Y.; Lemmens, I.; Rolland, T.; Coté, A. G.; Gebbia, M.; Kishore, N.; Knapp, J. J.; Mellor, J. C.; Reimand, J.; Tavernier, J.; Cusick, M. E.; Falter-Braun, P.; Spirohn, K.; Zhong, Q.; Aloy, P.; Hao, T.; Charloteaux, B.; Roth, F. P.; Hill, D. E.; Calderwood, M. A.; Twizere, J.-C; Vidal, M. Binary interactome models of inner-versus outer-complexome organisation (bioRxiv, 2021). 10.1101/2021.03.16.435663

48 Wan, M. S. M. et al. Mechanism of assembly, activation and lysine selection by the SIN3B histone deacetylase complex. Nat Commun 14, 2556 (2023). 10.1038/s41467-023-38276-0

## METHODS REFERENCES

49 Yu, H. et al. High-quality binary protein interaction map of the yeast interactome network. Science 322, 104–110 (2008). 10.1126/science.1158684

50 Baganz, F., Hayes, A., Marren, D., Gardner, D. C. & Oliver, S. G. Suitability of replacement markers for functional analysis studies in Saccharomyces cerevisiae. Yeast 13, 1563–1573 (1997). 10.1002/(SICI)1097-0061(199712)13:16<1563::AID-YEA240>3.0.CO;2-6

51 Wendland, J. PCR-based methods facilitate targeted gene manipulations and cloning procedures. Curr Genet 44, 115–123 (2003). 10.1007/s00294-003-0436-x

52 Gardner, J. M. & Jaspersen, S. L. Manipulating the yeast genome: deletion, mutation, and tagging by PCR. Methods Mol Biol 1205, 45–78 (2014). 10.1007/978-1-4939-1363-3_5

53 Guldener, U., Heck, S., Fielder, T., Beinhauer, J. & Hegemann, J. H. A new efficient gene disruption cassette for repeated use in budding yeast. Nucleic Acids Res 24, 2519–2524 (1996). 10.1093/nar/24.13.2519

54 Choi, S. G., Jia, J., Pfeffer, R. L. & Sealfon, S. C. G proteins and autocrine signaling differentially regulate gonadotropin subunit expression in pituitary gonadotrope. J Biol Chem 287, 21550–21560 (2012). 10.1074/jbc.M112.348607

55 Teste, M. A., Duquenne, M., Francois, J. M. & Parrou, J. L. Validation of reference genes for quantitative expression analysis by real-time RT-PCR in Saccharomyces cerevisiae. BMC Mol Biol 10, 99 (2009). 10.1186/1471-2199-10-99

56 van der Feltz, C. & Pomeranz Krummel, D. Purification of Native Complexes for Structural Study Using a Tandem Affinity Tag Method. J Vis Exp (2016). 10.3791/54389

57 Hsu, C. W. et al. Identification of HDAC Inhibitors Using a Cell-Based HDAC I/II Assay. J Biomol Screen 21, 643–652 (2016). 10.1177/1087057116629381

58 Zheng, S. Q. et al. MotionCor2: anisotropic correction of beam-induced motion for improved cryo-electron microscopy. Nat Methods 14, 331–332 (2017). 10.1038/nmeth.4193

59 Kimanius, D., Dong, L., Sharov, G., Nakane, T. & Scheres, S. H. W. New tools for automated cryo-EM single-particle analysis in RELION-4.0. Biochem J 478, 4169–4185 (2021). 10.1042/BCJ20210708

60 Punjani, A., Rubinstein, J. L., Fleet, D. J. & Brubaker, M. A. cryoSPARC: algorithms for rapid unsupervised cryo-EM structure determination. Nat Methods 14, 290–296 (2017). 10.1038/nmeth.4169

61 Punjani, A., Zhang, H. & Fleet, D. J. Non-uniform refinement: adaptive regularization improves single-particle cryo-EM reconstruction. Nat Methods 17, 1214–1221 (2020). 10.1038/s41592-020-00990-8

62 Tan, Y. Z. et al. Addressing preferred specimen orientation in single-particle cryo-EM through tilting. Nat Methods 14, 793–796 (2017). 10.1038/nmeth.4347

63 Rohou, A. & Grigorieff, N. CTFFIND4: Fast and accurate defocus estimation from electron micrographs. J Struct Biol 192, 216–221 (2015). 10.1016/j.jsb.2015.08.008

64 Zivanov, J., Nakane, T. & Scheres, S. H. W. A Bayesian approach to beam-induced motion correction in cryo-EM single-particle analysis. IUCrJ 6, 5–17 (2019). 10.1107/S205225251801463X

65 Emsley, P., Lohkamp, B., Scott, W. G. & Cowtan, K. Features and development of Coot. Acta Crystallogr D Biol Crystallogr 66, 486–501 (2010). 10.1107/S0907444910007493

66 Afonine, P. V. et al. New tools for the analysis and validation of cryo-EM maps and atomic models. Acta Crystallogr D Struct Biol 74, 814–840 (2018). 10.1107/S2059798318009324

67 Williams, C. J. et al. MolProbity: More and better reference data for improved all-atom structure validation. Protein Sci 27, 293–315 (2018). 10.1002/pro.3330

68 Afonine, P. V. et al. Real-space refinement in PHENIX for cryo-EM and crystallography. Acta Crystallogr D Struct Biol 74, 531–544 (2018). 10.1107/S2059798318006551

69 Liebschner, D. et al. Macromolecular structure determination using X-rays, neutrons and electrons: recent developments in Phenix. Acta Crystallogr D Struct Biol 75, 861–877 (2019). 10.1107/S2059798319011471

70 Meng, E. C. et al. UCSF ChimeraX: Tools for structure building and analysis. Protein Sci 32, e4792 (2023). 10.1002/pro.4792

71 Kong, A. T., Leprevost, F. V., Avtonomov, D. M., Mellacheruvu, D. & Nesvizhskii, A. I. MSFragger: ultrafast and comprehensive peptide identification in mass spectrometry-based proteomics. Nat Methods 14, 513–520 (2017). 10.1038/nmeth.4256

72 Teo, G. et al. SAINTexpress: improvements and additional features in Significance Analysis of INTeractome software. J Proteomics 100, 37–43 (2014). 10.1016/j.jprot.2013.10.023

73 Hsiao, Y. et al. Analysis and Visualization of Quantitative Proteomics Data Using FragPipe-Analyst. J Proteome Res 23, 4303–4315 (2024). 10.1021/acs.jproteome.4c00294

74 Combe, C. W., Graham, M., Kolbowski, L., Fischer, L. & Rappsilber, J. xiVIEW: Visualisation of Crosslinking Mass Spectrometry Data. J Mol Biol 436, 168656 (2024). 10.1016/j.jmb.2024.168656

75 Lagerwaard, I. M. A., P.; Jankevics, A.; Scheltema, R. A. Xlink Mapping and AnalySis (XMAS) - Smooth Integrative Modeling in ChimeraX (bioRxiv, 2022).

76 Honorato, R. V. et al. The HADDOCK2.4 web server for integrative modeling of biomolecular complexes. Nat Protoc 19, 3219–3241 (2024). 10.1038/s41596-024-01011-0

77 Rodrigues, J., Teixeira, J. M. C., Trellet, M. & Bonvin, A. pdb-tools: a swiss army knife for molecular structures. F1000Res 7, 1961 (2018). 10.12688/f1000research.17456.1

78 Dreze, M. et al. High-quality binary interactome mapping. Methods Enzymol 470, 281–315 (2010). 10.1016/S0076-6879(10)70012-4

79 Luck, K. et al. A reference map of the human binary protein interactome. Nature 580, 402–408 (2020). 10.1038/s41586-020-2188-x

80 Lievens, S. et al. Kinase Substrate Sensor (KISS), a mammalian in situ protein interaction sensor. Mol Cell Proteomics 13, 3332–3342 (2014). 10.1074/mcp.M114.041087

81 Moon, A. F., Mueller, G. A., Zhong, X. & Pedersen, L. C. A synergistic approach to protein crystallization: combination of a fixed-arm carrier with surface entropy reduction. Protein Sci 19, 901–913 (2010). 10.1002/pro.368

82 Kabsch, W. Integration, scaling, space-group assignment and post-refinement. Acta Crystallogr D Biol Crystallogr 66, 133–144 (2010). 10.1107/S0907444909047374

83 Adams, P. D. et al. PHENIX: a comprehensive Python-based system for macromolecular structure solution. Acta Crystallogr D Biol Crystallogr 66, 213–221 (2010). 10.1107/S0907444909052925

84 Emsley, P. & Cowtan, K. Coot: model-building tools for molecular graphics. Acta Crystallogr D Biol Crystallogr 60, 2126–2132 (2004). 10.1107/S0907444904019158

85 McCoy, A. J. et al. Phaser crystallographic software. J Appl Crystallogr 40, 658–674 (2007). 10.1107/S0021889807021206

