## Supplementary Information for "Dynamic engagement of dual-role regulators by the Sin3 complex"

Julien Olivet *et al.*

**a**

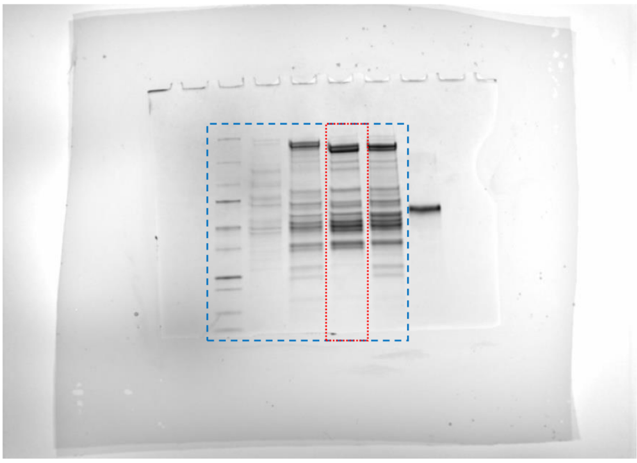

**b**

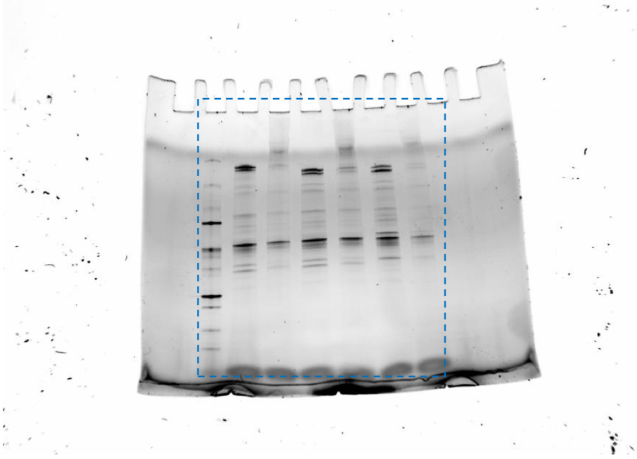

**c**

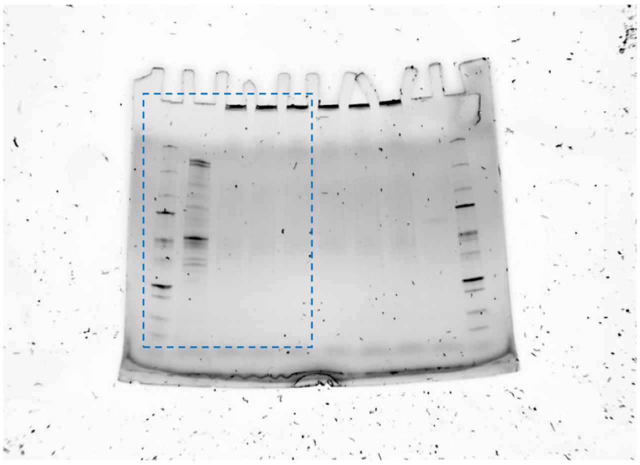

**Supplementary Figure 1 | Gels reported in this study.**
**a**, Uncropped Coomassie-stained SDS-PAGE from Fig. 1e (red) and Extended Data Fig. 2a (blue).
**b**, Uncropped unstained SDS-PAGE from Extended Data Fig. 2d (blue).
**c**, Uncropped unstained SDS-PAGE from Extended Data Fig. 4c (blue).

**Supplementary Table 1 | Enrichment of the Sin3L complex subunits in the Sap30-TAP purified sample by quantitative mass spectrometry.**

| Pull-down | Subunit | Fold-change enrichment over no tag control | Adjusted P-value |
|---|---|---|---|
| Sap30-TAP | Sin3 | 134 | 0.00435 |
| Sap30-TAP | Ume1 | 96 | 0.000652 |
| Sap30-TAP | Rpd3 | 55 | 0.000798 |
| Sap30-TAP | Cti6 | 42 | 0.000862 |
| Sap30-TAP | Pho23 | 33 | 0.000288 |
| Sap30-TAP | Rxt2 | 25 | 0.00033 |
| Sap30-TAP | Sap30 | 24 | 0.00793 |
| Sap30-TAP | Ume6 | 21 | 0.00913 |
| Sap30-TAP | Ash1 | 17 | 0.0111 |
| Sap30-TAP | Dep1 | 17 | 0.000248 |
| Sap30-TAP | Rxt3 | 15 | 0.0156 |
| Sap30-TAP | Sds3 | 10 | 0.00163 |

**Supplementary Table 2 | Summary of XL-MS results mapped to Sin3L complex models.** ISM: integrated structural modeling.

| Model | Min. Dist. ( Å) | Max. Dist. (Å) | Mean Dist. (Å) | Median Dist. (Å) | Total XL matched | Total XL mapped | Percent match |
|---|---|---|---|---|---|---|---|
| Core structure Sap30-TAP (cryo-EM; PDB: 9TAF) | 5.1 | 128.8 | 33.7 | 17.2 | 79 | 107 | 73.8 |
| Extended structure (ISM; PDB: 9AAF) | 6.7 | 130.9 | 37.7 | 29.3 | 89 | 146 | 61.0 |
| Extended+Cti6 (ISM; PDB: 9AAG) | 5.2 | 131.1 | 36.8 | 29.9 | 140 | 231 | 60.6 |
| Extended+Cti6+Ash1 (ISM; PDB: 9AAH) | 5.2 | 131.4 | 39.0 | 32.6 | 150 | 266 | 56.4 |

**Supplementary Table 3 | Top 10 conserved deleterious and enriched mutations identified in the minimal Sin3-Ume6 interface.**

| Yeast Sin3 residue | Human SIN3A ortholog residue | Mutation | Z score | Fold change | Significant enrichment | Significant depletion |
|---|---|---|---|---|---|---|
| L433 | L329 | L433R | -10.02 | 0.22 | No | Yes |
| K419 | K315 | K419Q | -9.67 | 0.23 | No | Yes |
| L471 | L380 | L471V | -9.34 | 0.24 | No | Yes |
| P472 | P381 | P472A | -7.97 | 0.30 | No | Yes |
| P472 | P381 | P472R | -7.74 | 0.31 | No | Yes |
| E442 | E338 | E442Q | -6.22 | 0.39 | No | Yes |
| L456 | L365 | L456F | -6.03 | 0.40 | No | Yes |
| Y429 | Y325 | Y429H | -5.97 | 0.40 | No | Yes |
| I435 | I331 | I435R | -5.91 | 0.41 | No | Yes |
| Y414 | Y310 | Y414S | -5.90 | 0.41 | No | Yes |
| K417 | K313 | K417T | 5.16 | 2.16 | Yes | No |
| L456 | L365 | L456R | 3.43 | 1.66 | Yes | No |
| D473 | D382 | D473E | 3.28 | 1.63 | Yes | No |
| Q443 | Q339 | Q443K | 2.98 | 1.56 | Yes | No |
| L436 | L332 | L436R | 2.75 | 1.50 | Yes | No |
| Q443 | Q339 | Q443L | 2.58 | 1.46 | Yes | No |
| V453 | V362 | V453G | 2.53 | 1.45 | Yes | No |
| I418 | I314 | I418F | 2.47 | 1.44 | Yes | No |
| N416 | N312 | N416Y | 2.42 | 1.43 | Yes | No |
| E407 | E303 | E407V | 2.24 | 1.39 | Yes | No |

**Supplementary Table 4 | Yeast *S. cerevisiae* strains used in this study.**

| Strain | Genotype | Source |
|---|---|---|
| MaV108 | *MATa leu2-3,112 trp1-901 his3-200 ura3-52 ade2-101 gal4Δ gal80Δ SPAL10::URA3@ura3 GAL1::LacZ* $can1^R$ $cyh2^R$ | Vidal et al. *PNAS* 1996 |
| MaV118 | *MaV108*, *pdr5::HIS3* | This paper |
| MaV208 | *MaV118*, *snq2::KanMX* | This paper |
| JOY111 | *MaV208*, *rpd3Δ::NatMX* | This paper |
| JOY112 | *MaV208*, *ume6Δ::NatMX* | This paper |
| JOY113 | *MaV208*, *sds3Δ::NatMX* | This paper |
| JOY114 | *MaV208*, *sap30Δ::NatMX* | This paper |
| JOY115 | *MaV208*, *pho23Δ::NatMX* | This paper |
| JOY116 | *MaV208*, *sin3Δ::NatMX* | This paper |
| JOY117 | *MaV208*, *rxt2Δ::NatMX* | This paper |
| JOY118 | *MaV208*, *rxt3Δ::NatMX* | This paper |
| JOY119 | *MaV208*, *dep1Δ::NatMX* | This paper |
| JOY120 | *MaV208*, *cti6Δ::NatMX* | This paper |
| JOY121 | *MaV208*, *ume1Δ::NatMX* | This paper |
| JOY122 | *MaV208*, *ash1Δ::NatMX* | This paper |
| JOY128 | *MaV208*, *(spal10::ura3)Δ::HpHMX@ura3* | This paper |
| BY4741 | *MATa his3Δ1 leu2Δ0 met15Δ0 ura3Δ0* | Open biosystems (Horizon Discovery) |
| Sin3-TAP | *BY4741, SIN3-TAP::HIS3MX6* | Open biosystems (Horizon Discovery) |
| Sap30-TAP | *BY4741, SAP30-TAP::HIS3MX6* | Gift from Stephen Buratowski, Harvard Medical School |
| Rxt3-TAP | *BY4741, RXT3-TAP::HIS3MX6* | Open biosystems (Horizon Discovery) |
| HOY013 | *Y8800*, *SPAL10::URA3@ura3* | This paper |
| HOY014 | *Y8930*, *SPAL10::URA3@ura3* | This paper |
| AY280 | *HoY13 <pDEST-AD-UME6 (aa 265-726>* | This paper |
| AY285 | *HoY13 <pDEST-AD>* | This paper |
| AY286 | *HoY13 <pDEST-AD-CBCB>* | This paper |
| AY287 | *HoY13 <pDEST-AD-XIAP>* | This paper |
| AY288 | *HoY13 <pDEST-AD>* | This paper |
| AY289 | *HoY13 <pDEST-AD>* | This paper |
| AY290 | *HoY13 <pDEST-AD-UME6>* | This paper |
| AY291 | *HoY14 <pDEST-DB>* | This paper |
| AY292 | *HoY14 <pDEST-DB-GRB2>* | This paper |
| AY293 | *HoY14 <pDEST-DB-CSAP9>* | This paper |
| AY294 | *HoY14 <pDEST-DB-GAL4>* | This paper |
| AY295 | *HoY14 <pDEST-DB-WDR62>* | This paper |
| AY296 | *HoY14 <pDEST-DB>* | This paper |

**Supplementary Table 5 | Oligonucleotide primer sequences used in this study.**

| Name | Sequence (5' – 3') | Orientation | Purpose |
|---|---|---|---|
| UBC6 | AATTGGATGAGGGGGATGCG | Forward | RT-qPCR |
| UBC6 | CGCTTGTTCAGCGCGTATTC | Reverse | |
| TAF10 | AACAACAGTCAGGCGAGAGC | Forward | |
| TAF10 | AACAGCGCTACTGAGATCGT | Reverse | |
| TRK2 | GCGCTTATGGTACAGTGGGT | Forward | |
| TRK2 | CCAGTTTGTCACTTGGCAGC | Reverse | |
| SPO13 | AACGGCCGGAGTTGCTTTAT | Forward | |
| SPO13 | CTGCCCTTGGTGTCTCTTGG | Reverse | |
| CAR1 | ATACGGCATCAACGCTGTCA | Forward | |
| CAR1 | GGTCAACCCACCTCTCACTG | Reverse | |
| INO1 | TATATGAAGCCCGTCGGGGA | Forward | |
| INO1 | TCGATGATCAAGGGCGTAGC | Reverse | |
| IME2 | GGTGATGCCTCTTTAGGCGA | Forward | |
| IME2 | TGAGGCTCGAACTTTTCCCG | Reverse | |
| URA3 | TGACATTGCGAAGAGCGACA | Forward | |
| URA3 | GAGACCACATCATCCACGGT | Reverse | |
| Y2H-DB | GGCTTCAGTGGAGACTGATATGCCTC | Forward | Y2H |
| Y2H-term | GGAGACTTGACCAAACCTCTGGCG | Reverse | |
| Y2H-AD | CGCGTTTGGAATCACTACAGGG | Forward | |
| FP387 | CCACCAAACCCAAAAAAAGAGGGTGGGTCGAATCAAACAAGTTTG | Forward | |
| FP388 | ACTTAGAGCTCGACGTCTTACTTACTTAGCGGCCATCAAACCACT | Reverse | |
| FP389 | CAAAGACAGTTGACTGTATCGTCGAGGTCGAATCAAACAAGTTTG | Forward | |
| FP390 | ACTTAGAGCTCGACGTCTTACTTACTTAGCGGCCATCAAACCACT | Reverse | |
| FP391 | GGGGACAACTTTGTACAAAAAAGTTGGCATGTCACAGGTTTGGCATAATTCGAATT | Forward | |
| FP392 | GGGGACAACTTTGTACAAGAAAGTTGGCTTGAATCTTAGCCCCCTTGTCTGAAGA | Reverse | |
| AP427 | CAAAGACAGTTGACTGTATCGTCGAGGTCGAATCAAACAAGTTTG | Forward | Mutational scanning |
| AP267 | ACTTAGAGCTCGACGTCTTACTTACTTAGCGGCCATCAAACCACT | Reverse | |
| AP434 | GGTCGAATCAAACAAGTTTGGGTTATCCAATTTTGATTCAAGGGT | Forward | |
| AP438 | CTTAGCGGCCATCAAACCACTCTCATGCAAAGCATCAATAATCTCG | Reverse | |

**Supplementary Table 6 | Oligonucleotide primer sequences used to generate the library of DNA fragments.**

| Gene | Sequence (5' – 3') | Start (nt.) | Orientation |
|---|---|---|---|
| *ASH1* | GGTCGAATCAAACAAGTTTGATGTCAAGCTTATACATCAAAACACCA | 1 | Forward |
| *ASH1* | GGTCGAATCAAACAAGTTTGTCAAGCTTATACATCAAAACACCACT | 4 | Forward |
| *ASH1* | GGTCGAATCAAACAAGTTTGAGCTTATACATCAAAACACCACTGC | 7 | Forward |
| *ASH1* | GGTCGAATCAAACAAGTTTGATAAATTCAGCAAGTCCTAGGAAGT | 154 | Forward |
| *ASH1* | GGTCGAATCAAACAAGTTTGTTAACACTAGGCAATTCACCAGA | 307 | Forward |
| *ASH1* | GGTCGAATCAAACAAGTTTGTCTTTATCTAAGAGACCGGAGCG | 451 | Forward |
| *ASH1* | GGTCGAATCAAACAAGTTTGTTCAAGCCATTAAGTATACCCAACT | 658 | Forward |
| *ASH1* | GGTCGAATCAAACAAGTTTGTTCCATTCCCCGTCTAAGGA | 751 | Forward |
| *ASH1* | GGTCGAATCAAACAAGTTTGATGGAGCGCACGACGCTTAG | 901 | Forward |
| *ASH1* | GGTCGAATCAAACAAGTTTGCCAAATGCCAGTCACAAAAAGAC | 1051 | Forward |
| *ASH1* | GGTCGAATCAAACAAGTTTGGGTAACACTGCAAGACAATTGGT | 1201 | Forward |
| *ASH1* | GGTCGAATCAAACAAGTTTGCCAAGCCCAAGGTCACCC | 1351 | Forward |
| *ASH1* | GGTCGAATCAAACAAGTTTGTCGTGTCATTCGAGTGATTCG | 1501 | Forward |
| *ASH1* | CTTAGCGGCCATCAAACCACTATTGTTATCGGTTGTGATATTGTTCC | 153 | Reverse |
| *ASH1* | CTTAGCGGCCATCAAACCACTTGAGGAAATTGGATTGTAATCCA | 306 | Reverse |
| *ASH1* | CTTAGCGGCCATCAAACCACTGACGGACGATAGCCTGTCTA | 450 | Reverse |
| *ASH1* | CTTAGCGGCCATCAAACCACTATGAGCCACTTCTTCGCGTA | 657 | Reverse |
| *ASH1* | CTTAGCGGCCATCAAACCACTCTTGGACGACCTAGTCGATTC | 750 | Reverse |
| *ASH1* | CTTAGCGGCCATCAAACCACTGGTTCTATTGGTTGGTGGACTC | 900 | Reverse |
| *ASH1* | CTTAGCGGCCATCAAACCACTCATGGGAAATGAATTTCCACGT | 1050 | Reverse |
| *ASH1* | CTTAGCGGCCATCAAACCACTGAAAACATTTGATGCTTGCTTTGA | 1200 | Reverse |
| *ASH1* | CTTAGCGGCCATCAAACCACTAGTGTATGCCTTGGGACGC | 1350 | Reverse |
| *ASH1* | CTTAGCGGCCATCAAACCACTCACGCACACTCTTGTGGTG | 1500 | Reverse |
| *ASH1* | CTTAGCGGCCATCAAACCACTATTCTCTACTGTCTCAGTTATGT | 1764 | Reverse |
| *CTI6* | GGTCGAATCAAACAAGTTTGATGGAATCGACAGCAATAGTGC | 1 | Forward |
| *CTI6* | GGTCGAATCAAACAAGTTTGGAATCGACAGCAATAGTGCCT | 4 | Forward |
| *CTI6* | GGTCGAATCAAACAAGTTTGTCGACAGCAATAGTGCCTAA | 7 | Forward |
| *CTI6* | GGTCGAATCAAACAAGTTTGAAACAAGAAGATGTTCCTATGGAAGG | 157 | Forward |
| *CTI6* | GGTCGAATCAAACAAGTTTGTGTGTGAGTATTACTCAAGACAACG | 307 | Forward |
| *CTI6* | GGTCGAATCAAACAAGTTTGGCAAGTAAATCTCATGCCGCC | 472 | Forward |
| *CTI6* | GGTCGAATCAAACAAGTTTGAAGGACGGCAACACGAGG | 604 | Forward |
| *CTI6* | GGTCGAATCAAACAAGTTTGAGAATGCTCGAAAAAGCCTTGA | 760 | Forward |
| *CTI6* | GGTCGAATCAAACAAGTTTGGAAGGCAAACCGGACTCGG | 907 | Forward |
| *CTI6* | GGTCGAATCAAACAAGTTTGGAGAAAACTGATGAACCCATCCT | 1051 | Forward |
| *CTI6* | GGTCGAATCAAACAAGTTTGAACCAAAATAGAAGGAGTGCTGA | 1201 | Forward |
| *CTI6* | GGTCGAATCAAACAAGTTTGCAGTCTGATCGAGAGGAATTTGT | 1351 | Forward |
| *CTI6* | CTTAGCGGCCATCAAACCACTTACACTTTCTTCCTGTACACTTCC | 156 | Reverse |
| *CTI6* | CTTAGCGGCCATCAAACCACTATAACCGTGCTGCCAGGAA | 306 | Reverse |
| *CTI6* | CTTAGCGGCCATCAAACCACTAGCTGCACTTCGTGCCTT | 471 | Reverse |
| *CTI6* | CTTAGCGGCCATCAAACCACTGGCTAATGCTACAGCCGAAG | 603 | Reverse |
| *CTI6* | CTTAGCGGCCATCAAACCACTTTGATATTGCTTTTCTTCTCTTGCC | 759 | Reverse |
| *CTI6* | CTTAGCGGCCATCAAACCACTCGTTAGCATAACATCCGTTTGT | 906 | Reverse |
| *CTI6* | CTTAGCGGCCATCAAACCACTAGTGTCTTGCGCCGAACC | 1050 | Reverse |
| *CTI6* | CTTAGCGGCCATCAAACCACTGGTGTCTGAACTTGTATTGCT | 1200 | Reverse |
| *CTI6* | CTTAGCGGCCATCAAACCACTATCTTCACTCAATTCCCATTGAGT | 1350 | Reverse |
| *CTI6* | CTTAGCGGCCATCAAACCACTTTGAATGGCATTAGTGTTATTTGA | 1518 | Reverse |

| | | | |
|---|---|---|---|
| *DEP1* | GGTCGAATCAAACAAGTTTGATGAGTCAGCAAACACCACAG | 1 | Forward |
| *DEP1* | GGTCGAATCAAACAAGTTTGAGTCAGCAAACACCACAGGA | 4 | Forward |
| *DEP1* | GGTCGAATCAAACAAGTTTGCAGCAAACACCACAGGAAAGT | 7 | Forward |
| *DEP1* | GGTCGAATCAAACAAGTTTGACAGAGAAGATGGATAGCGACG | 151 | Forward |
| *DEP1* | GGTCGAATCAAACAAGTTTGAAAGTGCCGGGAGAGAACG | 313 | Forward |
| *DEP1* | GGTCGAATCAAACAAGTTTGAACGAGGAGGATAATGAAAACGA | 451 | Forward |
| *DEP1* | GGTCGAATCAAACAAGTTTGCAATTGGTGCGGTTGCAAAC | 610 | Forward |
| *DEP1* | GGTCGAATCAAACAAGTTTGATCAACACAGAAACAATCGCTACC | 754 | Forward |
| *DEP1* | GGTCGAATCAAACAAGTTTGCCAGATGTCAATTACCACGTCC | 901 | Forward |
| *DEP1* | GGTCGAATCAAACAAGTTTGAACCCGGTGGACAAACTCG | 1051 | Forward |
| *DEP1* | CTTAGCGGCCATCAAACCACTTCCTGCGTCACTGGATATACA | 150 | Reverse |
| *DEP1* | CTTAGCGGCCATCAAACCACTCATCTTCTTGGCCTCATCTATCG | 312 | Reverse |
| *DEP1* | CTTAGCGGCCATCAAACCACTGTTTTCCTCCTCATTATCACCT | 450 | Reverse |
| *DEP1* | CTTAGCGGCCATCAAACCACTATTGTCATATAGTTTTTGGCGCA | 609 | Reverse |
| *DEP1* | CTTAGCGGCCATCAAACCACTGCATGAAAGCTCGTACTTCTGT | 753 | Reverse |
| *DEP1* | CTTAGCGGCCATCAAACCACTGATGACTATATCCATATCGCGGC | 900 | Reverse |
| *DEP1* | CTTAGCGGCCATCAAACCACTGGCTCTGTAGCGGTACTCG | 1050 | Reverse |
| *DEP1* | CTTAGCGGCCATCAAACCACTCTGGGCCCACTGGTGG | 1215 | Reverse |
| *PHO23* | GGTCGAATCAAACAAGTTTGATGAGTTCACCAGCGAACCT | 1 | Forward |
| *PHO23* | GGTCGAATCAAACAAGTTTGAGTTCACCAGCGAACCTATTCC | 4 | Forward |
| *PHO23* | GGTCGAATCAAACAAGTTTGTCACCAGCGAACCTATTCCC | 7 | Forward |
| *PHO23* | GGTCGAATCAAACAAGTTTGATGCCGAATTTGAACGAGAGG | 127 | Forward |
| *PHO23* | GGTCGAATCAAACAAGTTTGTCGCTGGAGGAGAAAATGCA | 244 | Forward |
| *PHO23* | GGTCGAATCAAACAAGTTTGCGGTTAGAATTGGCGTATGAAGT | 307 | Forward |
| *PHO23* | GGTCGAATCAAACAAGTTTGAAATCGTCGCAGGCACTGAA | 442 | Forward |
| *PHO23* | GGTCGAATCAAACAAGTTTGAGGCAGGGCGAACATTACTC | 499 | Forward |
| *PHO23* | GGTCGAATCAAACAAGTTTGACTGGTAACAACACAAACTCAAGA | 595 | Forward |
| *PHO23* | GGTCGAATCAAACAAGTTTGGCCGTTTCACCAAGCACTATC | 694 | Forward |
| *PHO23* | GGTCGAATCAAACAAGTTTGAACAACAGCAGGATATCAAGACCA | 793 | Forward |
| *PHO23* | GGTCGAATCAAACAAGTTTGGGCGCAGACTGTGAGCTAG | 892 | Forward |
| *PHO23* | CTTAGCGGCCATCAAACCACTAGAATGCACACATTTTGCATCT | 126 | Reverse |
| *PHO23* | CTTAGCGGCCATCAAACCACTTGGCATCAGTTCTTCATAAATCT | 243 | Reverse |
| *PHO23* | CTTAGCGGCCATCAAACCACTGGACGTCAATCTGTCTAGAT | 306 | Reverse |
| *PHO23* | CTTAGCGGCCATCAAACCACTGCTGTTTGATTTGCTCTCTA | 441 | Reverse |
| *PHO23* | CTTAGCGGCCATCAAACCACTCCTGTTGGCAGCCATGG | 498 | Reverse |
| *PHO23* | CTTAGCGGCCATCAAACCACTGTGGTCTTGGCTCTCGTGC | 594 | Reverse |
| *PHO23* | CTTAGCGGCCATCAAACCACTTGTGGTGGCAACTCTCCTC | 693 | Reverse |
| *PHO23* | CTTAGCGGCCATCAAACCACTGCTGTTTCCGACGCTGCTAA | 792 | Reverse |
| *PHO23* | CTTAGCGGCCATCAAACCACTATCACACCCCACCATTTCCC | 891 | Reverse |
| *PHO23* | CTTAGCGGCCATCAAACCACTCAGTTTTTTTTTGCAGTCGTCG | 990 | Reverse |
| *RPD3* | GGTCGAATCAAACAAGTTTGATGGTATATGAAGCAACACCT | 1 | Forward |
| *RPD3* | GGTCGAATCAAACAAGTTTGGTATATGAAGCAACACCTTTTGATCC | 4 | Forward |
| *RPD3* | GGTCGAATCAAACAAGTTTGTATGAAGCAACACCTTTTGATCCG | 7 | Forward |
| *RPD3* | GGTCGAATCAAACAAGTTTGGAAATTTACAGAGCTAAGCCGGC | 184 | Forward |
| *RPD3* | GGTCGAATCAAACAAGTTTGGATGATTGTCCTGTCTTTGATGGG | 322 | Forward |
| *RPD3* | GGTCGAATCAAACAAGTTTGCATGCAAAAAAATCGGAAGCTTCT | 451 | Forward |
| *RPD3* | GGTCGAATCAAACAAGTTTGGATCGTGTCATGACATGTTCT | 601 | Forward |
| *RPD3* | GGTCGAATCAAACAAGTTTGGCTGTCGTGTTACAGTGTGG | 796 | Forward |
| *RPD3* | GGTCGAATCAAACAAGTTTGATCCCAATGATGGTTGTTGGTG | 907 | Forward |
| *RPD3* | GGTCGAATCAAACAAGTTTGTATGCCCCTAGTGTTCAGTTGA | 1153 | Forward |

| | | | |
|---|---|---|---|
| *RPD3* | CTTAGCGGCCATCAAACCACTCATCTTCTTGTACAAGCCAT | 183 | Reverse |
| *RPD3* | CTTAGCGGCCATCAAACCACTTCCGACATTAAACTTGACACT | 321 | Reverse |
| *RPD3* | CTTAGCGGCCATCAAACCACTATGCAAACCACCCGCATAGT | 450 | Reverse |
| *RPD3* | CTTAGCGGCCATCAAACCACTCGTTGTATAAAACGCTTCCTCT | 600 | Reverse |
| *RPD3* | CTTAGCGGCCATCAAACCACTAGAAGGTTGATACCATTCCA | 795 | Reverse |
| *RPD3* | CTTAGCGGCCATCAAACCACTCCCAAAGGATTTCACATAGT | 906 | Reverse |
| *RPD3* | CTTAGCGGCCATCAAACCACTCTTTGTGTTTTCCAAATTAGCA | 1152 | Reverse |
| *RPD3* | CTTAGCGGCCATCAAACCACTATAGAATTCATTGTCATGCTCAACA | 1299 | Reverse |
| *RXT2* | GGTCGAATCAAACAAGTTTGATGACAGCATCTCCTGCAAA | 1 | Forward |
| *RXT2* | GGTCGAATCAAACAAGTTTGACAGCATCTCCTGCAAAAAAGA | 4 | Forward |
| *RXT2* | GGTCGAATCAAACAAGTTTGGCATCTCCTGCAAAAAAGAGGT | 7 | Forward |
| *RXT2* | GGTCGAATCAAACAAGTTTGAACTATCAGGTACTGAAAAGATCCCT | 187 | Forward |
| *RXT2* | GGTCGAATCAAACAAGTTTGGACTTGAACAATTCTAAGCCA | 304 | Forward |
| *RXT2* | GGTCGAATCAAACAAGTTTGGATGAAAACGACATCGATGGT | 454 | Forward |
| *RXT2* | GGTCGAATCAAACAAGTTTGTATTCCCAGTTTTTGGAGGT | 700 | Forward |
| *RXT2* | GGTCGAATCAAACAAGTTTGGAGGATAGAGGGGCAAGCGA | 799 | Forward |
| *RXT2* | GGTCGAATCAAACAAGTTTGATGGAAGAATTTGGAGAGGAAGA | 904 | Forward |
| *RXT2* | GGTCGAATCAAACAAGTTTGTCGCAGATTGCACTACAGAGG | 1072 | Forward |
| *RXT2* | GGTCGAATCAAACAAGTTTGGACGATGATATTACTATCCCGGTGG | 1201 | Forward |
| *RXT2* | CTTAGCGGCCATCAAACCACTACCGCTTTTTTCCTTTATTATCCT | 186 | Reverse |
| *RXT2* | CTTAGCGGCCATCAAACCACTTCTTCTTGTTACTACTTCGCTTCT | 303 | Reverse |
| *RXT2* | CTTAGCGGCCATCAAACCACTACCGACTAAAACAGGCTCGA | 453 | Reverse |
| *RXT2* | CTTAGCGGCCATCAAACCACTTCTAACAACTGACATCTGTTCT | 699 | Reverse |
| *RXT2* | CTTAGCGGCCATCAAACCACTTGGAAGGGTCAAATTGTGAT | 798 | Reverse |
| *RXT2* | CTTAGCGGCCATCAAACCACTGCCATTATTGATATGACCAGCT | 903 | Reverse |
| *RXT2* | CTTAGCGGCCATCAAACCACTCAGCTGTCGGGCAGACT | 1071 | Reverse |
| *RXT2* | CTTAGCGGCCATCAAACCACTAGATATACCCAAATATTCCCGACTC | 1200 | Reverse |
| *RXT2* | CTTAGCGGCCATCAAACCACTGGCTCTATTAGCTTCTTCATCAGG | 1359 | Reverse |
| *RXT3* | GGTCGAATCAAACAAGTTTGATGTCGGTAAGCGAACAAGA | 1 | Forward |
| *RXT3* | GGTCGAATCAAACAAGTTTGTCGGTAAGCGAACAAGATCCT | 4 | Forward |
| *RXT3* | GGTCGAATCAAACAAGTTTGGTAAGCGAACAAGATCCTAATAGGG | 7 | Forward |
| *RXT3* | GGTCGAATCAAACAAGTTTGAATAAGCAAGAGGAGGGACAAGA | 100 | Forward |
| *RXT3* | GGTCGAATCAAACAAGTTTGGTAAGTTCACAGTCGGTTCTTGC | 199 | Forward |
| *RXT3* | GGTCGAATCAAACAAGTTTGTTTGATGAAGACTCCACAAGT | 298 | Forward |
| *RXT3* | GGTCGAATCAAACAAGTTTGATCTATTCTGATGACTCAGACCCA | 508 | Forward |
| *RXT3* | GGTCGAATCAAACAAGTTTGTTGAGAAGAACCCCTGTAAATGT | 595 | Forward |
| *RXT3* | GGTCGAATCAAACAAGTTTGCCTACTCTACAAAAGTATCCTAGCGT | 700 | Forward |
| *RXT3* | CTTAGCGGCCATCAAACCACTTTTTGTTCTCGCCGAATTTAGT | 99 | Reverse |
| *RXT3* | CTTAGCGGCCATCAAACCACTATTAGGCAGGTCGTAGGCC | 198 | Reverse |
| *RXT3* | CTTAGCGGCCATCAAACCACTAGACCCTTCTTTGAAACGATTGT | 297 | Reverse |
| *RXT3* | CTTAGCGGCCATCAAACCACTATCACATCCCCATATCTCAT | 507 | Reverse |
| *RXT3* | CTTAGCGGCCATCAAACCACTTTTGTGAAATGAGCCACCAGA | 594 | Reverse |
| *RXT3* | CTTAGCGGCCATCAAACCACTCAAAAACAACAACTCCACCTCT | 699 | Reverse |
| *RXT3* | CTTAGCGGCCATCAAACCACTGGTCCATTTTAAATTTTTAATATACCCGT | 882 | Reverse |
| *SAP30* | GGTCGAATCAAACAAGTTTGATGGCTAGGCCAGTTAATACA | 1 | Forward |
| *SAP30* | GGTCGAATCAAACAAGTTTGGCTAGGCCAGTTAATACAAACGC | 4 | Forward |
| *SAP30* | GGTCGAATCAAACAAGTTTGAGGCCAGTTAATACAAACGCTG | 7 | Forward |
| *SAP30* | GGTCGAATCAAACAAGTTTGAATAACGGGCCTACCTCCAG | 169 | Forward |
| *SAP30* | GGTCGAATCAAACAAGTTTGAATGGGAAGCAAAGACTCACAG | 199 | Forward |
| *SAP30* | GGTCGAATCAAACAAGTTTGCACCCAATGGATTTCGAAGAGT | 298 | Forward |

| | | | |
|---|---|---|---|
| *SAP30* | GGTCGAATCAAACAAGTTTGTCCAAATTGGGCGCGAAGA | 406 | Forward |
| *SAP30* | GGTCGAATCAAACAAGTTTGGATGAGCATTCCATTAAAGAGACAGA | 511 | Forward |
| *SAP30* | CTTAGCGGCCATCAAACCACTGTTATTGCTGTTATTGCTGT | 168 | Reverse |
| *SAP30* | CTTAGCGGCCATCAAACCACTGGTTCTCCCGCTGGAGGTAG | 198 | Reverse |
| *SAP30* | CTTAGCGGCCATCAAACCACTACTCTTGGGGCGCAGATC | 297 | Reverse |
| *SAP30* | CTTAGCGGCCATCAAACCACTACCCAATAAGTATCCCTGCAG | 405 | Reverse |
| *SAP30* | CTTAGCGGCCATCAAACCACTGAAATGCCTTCTCACCACGT | 510 | Reverse |
| *SAP30* | CTTAGCGGCCATCAAACCACTACCCCGAAATTCCATCTTGA | 603 | Reverse |
| *SDS3* | GGTCGAATCAAACAAGTTTGATGGCTATTCAAAAGGTTAGTAACAAGG | 1 | Forward |
| *SDS3* | GGTCGAATCAAACAAGTTTGGCTATTCAAAAGGTTAGTAACAAGGACT | 4 | Forward |
| *SDS3* | GGTCGAATCAAACAAGTTTGATTCAAAAGGTTAGTAACAAGGACTTGT | 7 | Forward |
| *SDS3* | GGTCGAATCAAACAAGTTTGCAAACGGATTTAACTTCTCTGCA | 142 | Forward |
| *SDS3* | GGTCGAATCAAACAAGTTTGGATCTAGAGTTAGTCAGGTTGCG | 220 | Forward |
| *SDS3* | GGTCGAATCAAACAAGTTTGCTAATGGATGTGGCCAATGTGC | 400 | Forward |
| *SDS3* | GGTCGAATCAAACAAGTTTGTCTTCGTCCAACGAATATGGA | 496 | Forward |
| *SDS3* | GGTCGAATCAAACAAGTTTGAAAGACACGAGAGGTAACAACA | 595 | Forward |
| *SDS3* | GGTCGAATCAAACAAGTTTGTTCCCCCATTTCAACAACCT | 694 | Forward |
| *SDS3* | GGTCGAATCAAACAAGTTTGAAGAACCCCAAAGACAATGCT | 802 | Forward |
| *SDS3* | CTTAGCGGCCATCAAACCACTCAATGCTGTCAGTCTATCTT | 141 | Reverse |
| *SDS3* | CTTAGCGGCCATCAAACCACTTCTTTCTTCCTCCAAATCTCGT | 219 | Reverse |
| *SDS3* | CTTAGCGGCCATCAAACCACTTAACCTCTCCTCCTGTAATTTCT | 399 | Reverse |
| *SDS3* | CTTAGCGGCCATCAAACCACTATCCCAACCACTTACTGTGTG | 495 | Reverse |
| *SDS3* | CTTAGCGGCCATCAAACCACTAGAAGCATTCCTTCTCCTTAGC | 594 | Reverse |
| *SDS3* | CTTAGCGGCCATCAAACCACTTTGCGACCCTTGTCTAGAAC | 693 | Reverse |
| *SDS3* | CTTAGCGGCCATCAAACCACTTTCGCCAAACAAAAATGCAT | 801 | Reverse |
| *SDS3* | CTTAGCGGCCATCAAACCACTGTCAGACCTTAGTCTGAAAGGAG | 981 | Reverse |
| *SIN3* | GGTCGAATCAAACAAGTTTGATGTCACAGGTTTGGCATAATTCG | 1 | Forward |
| *SIN3* | GGTCGAATCAAACAAGTTTGTCACAGGTTTGGCATAATTCGA | 4 | Forward |
| *SIN3* | GGTCGAATCAAACAAGTTTGCAGGTTTGGCATAATTCGAATTCG | 7 | Forward |
| *SIN3* | GGTCGAATCAAACAAGTTTGAATGGCCAACAGGCTCTAACT | 199 | Forward |
| *SIN3* | GGTCGAATCAAACAAGTTTGGTGGGAGCCGCCAGTTTTTC | 397 | Forward |
| *SIN3* | GGTCGAATCAAACAAGTTTGAACAACGAAAATTCTCACGATGA | 613 | Forward |
| *SIN3* | GGTCGAATCAAACAAGTTTGGGTTATCCAATTTTGATTCAAGGGT | 814 | Forward |
| *SIN3* | GGTCGAATCAAACAAGTTTGTTTCCAGAAAGCGATGGAAATGG | 991 | Forward |
| *SIN3* | GGTCGAATCAAACAAGTTTGGATGCTAAGAAAAACGTTGATGTCG | 1195 | Forward |
| *SIN3* | GGTCGAATCAAACAAGTTTGGACTCTTCAGCTTCTGCCAA | 1417 | Forward |
| *SIN3* | GGTCGAATCAAACAAGTTTGGGCCATCCTTCTAACCGAGG | 1585 | Forward |
| *SIN3* | GGTCGAATCAAACAAGTTTGCCTCTATCAGATCTAAGAACGTCTCT | 1783 | Forward |
| *SIN3* | GGTCGAATCAAACAAGTTTGTCGCTTAATGAGGAAGTCACT | 1981 | Forward |
| *SIN3* | GGTCGAATCAAACAAGTTTGTTTGGCCCAAGTTACAAGAGG | 2251 | Forward |
| *SIN3* | GGTCGAATCAAACAAGTTTGTCGGGATTTATTGCTCATCGT | 2377 | Forward |
| *SIN3* | GGTCGAATCAAACAAGTTTGCACCCTGCAGTGACAGCC | 2656 | Forward |
| *SIN3* | GGTCGAATCAAACAAGTTTGAGCAGCATCAAAGTTGATCAAACA | 2857 | Forward |
| *SIN3* | GGTCGAATCAAACAAGTTTGACCACAGCCTATTCTAATCCCG | 2995 | Forward |
| *SIN3* | GGTCGAATCAAACAAGTTTGAGGCCCTATCAACAAGAAATGAGT | 3169 | Forward |
| *SIN3* | GGTCGAATCAAACAAGTTTGGCAAACTCTCAGGGTATTATCCA | 3370 | Forward |
| *SIN3* | GGTCGAATCAAACAAGTTTGATGGGACTAGATTTTGTTGGTGA | 3580 | Forward |
| *SIN3* | GGTCGAATCAAACAAGTTTGGACGCTAAAACTGCGGAAAT | 3775 | Forward |
| *SIN3* | GGTCGAATCAAACAAGTTTGCTAAAGGAACCAAAGGCAGACG | 3970 | Forward |
| *SIN3* | GGTCGAATCAAACAAGTTTGGGGTCTTACGATGTTTTTACCCG | 4213 | Forward |

| | | | |
|---|---|---|---|
| *SIN3* | GGTCGAATCAAACAAGTTTGCTAAAAGACAGCATAGCAAAGACGA | 4399 | Forward |
| *SIN3* | CTTAGCGGCCATCAAACCACTGGAATCTCTTCTATCTTCCTCCT | 198 | Reverse |
| *SIN3* | CTTAGCGGCCATCAAACCACTTGTGGGGAGGGGAGCTG | 396 | Reverse |
| *SIN3* | CTTAGCGGCCATCAAACCACTATTATTGTCATCGTTAGCATCTGC | 612 | Reverse |
| *SIN3* | CTTAGCGGCCATCAAACCACTTCTGAACAAAGTGGATACTCTTTCA | 813 | Reverse |
| *SIN3* | CTTAGCGGCCATCAAACCACTAGAACCAAGTTCCTGTGCATC | 990 | Reverse |
| *SIN3* | CTTAGCGGCCATCAAACCACTTTCTTGAGGCACTAAAGATTGACT | 1194 | Reverse |
| *SIN3* | CTTAGCGGCCATCAAACCACTCGGCAAGAATTTCTTGAAATCT | 1416 | Reverse |
| *SIN3* | CTTAGCGGCCATCAAACCACTGTAATAGCCCGAAGCTGGAT | 1584 | Reverse |
| *SIN3* | CTTAGCGGCCATCAAACCACTTGGATTTTCATTCGCTATCCCT | 1782 | Reverse |
| *SIN3* | CTTAGCGGCCATCAAACCACTTATATTGTTTTCGATGGGCTCAGT | 1980 | Reverse |
| *SIN3* | CTTAGCGGCCATCAAACCACTTGCCTCACATAAATCTAAATCCA | 2250 | Reverse |
| *SIN3* | CTTAGCGGCCATCAAACCACTATCTTCGGAAGCCCATACAGG | 2376 | Reverse |
| *SIN3* | CTTAGCGGCCATCAAACCACTCTCATGCAAAGCATCAATAATCTCG | 2655 | Reverse |
| *SIN3* | CTTAGCGGCCATCAAACCACTAATCTCTGATATCAACTGCTTTGT | 2856 | Reverse |
| *SIN3* | CTTAGCGGCCATCAAACCACTATGGGTTATAAAAGTGTCAGCCA | 2994 | Reverse |
| *SIN3* | CTTAGCGGCCATCAAACCACTCTTCCTTGATGCAATAGAACTGC | 3168 | Reverse |
| *SIN3* | CTTAGCGGCCATCAAACCACTTTCCTCCACTAAATTCCCAGT | 3369 | Reverse |
| *SIN3* | CTTAGCGGCCATCAAACCACTTTCGGAAAGTTGACTCGATA | 3579 | Reverse |
| *SIN3* | CTTAGCGGCCATCAAACCACTAGTCATCAAGGTATGAGCATGC | 3774 | Reverse |
| *SIN3* | CTTAGCGGCCATCAAACCACTTGTCAAATCATCAAGTGCAA | 3969 | Reverse |
| *SIN3* | CTTAGCGGCCATCAAACCACTGTTTTCAATATGCAGTTGGTAGGT | 4212 | Reverse |
| *SIN3* | CTTAGCGGCCATCAAACCACTGTTCTCCCATTTTTTTTGCA | 4398 | Reverse |
| *SIN3* | CTTAGCGGCCATCAAACCACTTTGAATCTTAGCCCCCTTGTCT | 4608 | Reverse |
| *UME1* | GGTCGAATCAAACAAGTTTGATGAGCACTTTAGATATTGCAGA | 1 | Forward |
| *UME1* | GGTCGAATCAAACAAGTTTGAGCACTTTAGATATTGCAGAAGACA | 4 | Forward |
| *UME1* | GGTCGAATCAAACAAGTTTGACTTTAGATATTGCAGAAGACAACA | 7 | Forward |
| *UME1* | GGTCGAATCAAACAAGTTTGGTTTTCACTAACGATAGCTCATGT | 157 | Forward |
| *UME1* | GGTCGAATCAAACAAGTTTGCCGTTAGTACAACCGGATTATACCA | 301 | Forward |
| *UME1* | GGTCGAATCAAACAAGTTTGAATGGTTCTTTGGCATGGTTCA | 451 | Forward |
| *UME1* | GGTCGAATCAAACAAGTTTGGAAGTGCTTCGTACAATACCTGT | 673 | Forward |
| *UME1* | GGTCGAATCAAACAAGTTTGGACGGTATCATAAGGTTTTGGGG | 763 | Forward |
| *UME1* | GGTCGAATCAAACAAGTTTGGGGGCTCTCAAGGTTTGGG | 901 | Forward |
| *UME1* | GGTCGAATCAAACAAGTTTGTCCATTTCGCCTATGGAAGTGG | 1051 | Forward |
| *UME1* | GGTCGAATCAAACAAGTTTGACTGAGGGCTGCAGAAGAGA | 1207 | Forward |
| *UME1* | CTTAGCGGCCATCAAACCACTTATACTCCTAAGTGTCGATGGAGA | 156 | Reverse |
| *UME1* | CTTAGCGGCCATCAAACCACTTTCTGAAATGCTTTCAGGTT | 300 | Reverse |
| *UME1* | CTTAGCGGCCATCAAACCACTGGTAGAAAGGGCTATCACCT | 450 | Reverse |
| *UME1* | CTTAGCGGCCATCAAACCACTTCCTATTTTACCTGAGTTGTCT | 672 | Reverse |
| *UME1* | CTTAGCGGCCATCAAACCACTATCAGAACAAGTGGCGAATA | 762 | Reverse |
| *UME1* | CTTAGCGGCCATCAAACCACTACCACTAGTTCCCGTCATAA | 900 | Reverse |
| *UME1* | CTTAGCGGCCATCAAACCACTAGAAAACTCTATTTTGGACACTTGC | 1050 | Reverse |
| *UME1* | CTTAGCGGCCATCAAACCACTGTGATAAAATGCCATGCTTTCGG | 1206 | Reverse |
| *UME1* | CTTAGCGGCCATCAAACCACTACTTTTGGCAGCTCCAACCT | 1380 | Reverse |
| *UME6* | GGTCGAATCAAACAAGTTTGATGCTAGACAAGGCGCGC | 1 | Forward |
| *UME6* | GGTCGAATCAAACAAGTTTGCTAGACAAGGCGCGCTCTC | 4 | Forward |
| *UME6* | GGTCGAATCAAACAAGTTTGGACAAGGCGCGCTCTCAAAG | 7 | Forward |
| *UME6* | GGTCGAATCAAACAAGTTTGCCTACAATTTCCAGTGCTAGCA | 199 | Forward |
| *UME6* | GGTCGAATCAAACAAGTTTGACTACACTTCAAGATCGCCGT | 397 | Forward |
| *UME6* | GGTCGAATCAAACAAGTTTGTCCACTAATAGTAACACTGCCACT | 595 | Forward |

| | | | |
|---|---|---|---|
| *UME6* | GGTCGAATCAAACAAGTTTGGGTAAGACTACCAATTCGCCT | 793 | Forward |
| *UME6* | GGTCGAATCAAACAAGTTTGCAATGTGCCGTGGGCGT | 991 | Forward |
| *UME6* | GGTCGAATCAAACAAGTTTGAACTCTCGACATGCGAACAC | 1189 | Forward |
| *UME6* | GGTCGAATCAAACAAGTTTGTTTTACGGCGATTTGCCG | 1387 | Forward |
| *UME6* | GGTCGAATCAAACAAGTTTGTCATCCCCATATAGAACTCATGA | 1594 | Forward |
| *UME6* | GGTCGAATCAAACAAGTTTGGCTGTTTTTGATGAAGACCAGGA | 1783 | Forward |
| *UME6* | GGTCGAATCAAACAAGTTTGTCGAGAAGGTCAGTTGGAAACG | 1981 | Forward |
| *UME6* | GGTCGAATCAAACAAGTTTGGCAAAATCAAAGGCGAAACAGT | 2179 | Forward |
| *UME6* | CTTAGCGGCCATCAAACCACTATGGCGGGAGTTAGCTCCAT | 198 | Reverse |
| *UME6* | CTTAGCGGCCATCAAACCACTATTGGTCGTAGTAGTAGTATCGT | 396 | Reverse |
| *UME6* | CTTAGCGGCCATCAAACCACTGGTGGAGGAGTAAGGGAAAAC | 594 | Reverse |
| *UME6* | CTTAGCGGCCATCAAACCACTGTTGTTCTTGTTGGCGCTGG | 792 | Reverse |
| *UME6* | CTTAGCGGCCATCAAACCACTGTCATCATGCACATTATGCTGG | 990 | Reverse |
| *UME6* | CTTAGCGGCCATCAAACCACTTGTAGGCGTATCCGTGTTTG | 1188 | Reverse |
| *UME6* | CTTAGCGGCCATCAAACCACTAAACCCAGTAGAAGGTCTTGAC | 1386 | Reverse |
| *UME6* | CTTAGCGGCCATCAAACCACTTCTCATGTTTGATAGCACTGCTG | 1593 | Reverse |
| *UME6* | CTTAGCGGCCATCAAACCACTTGTAGAATTGTTGCTTTCGA | 1782 | Reverse |
| *UME6* | CTTAGCGGCCATCAAACCACTTTTTTTCAATGGTGGAACTTCCAG | 1980 | Reverse |
| *UME6* | CTTAGCGGCCATCAAACCACTCTTAGGCTTGACATTTTTTCCT | 2178 | Reverse |
| *UME6* | CTTAGCGGCCATCAAACCACTTTTTTTTTTCATTGCTCTTCTTTTGGC | 2508 | Reverse |

**Supplementary Table 7 | Suboptimal error-prone PCR conditions used for random mutagenesis.**

| | Primer 1 (μM) | Primer 2 (μM) | dATP (μM) | dGTP (μM) | dCTP (μM) | dTTP (μM) | $MgCl_2$ (mM) | $MnCl_2$ (μM) | Enzyme (units) | DNA (ng/μM) |
|---|---|---|---|---|---|---|---|---|---|---|
| A1 | 0.2 | 0.2 | 0.2 | 0.2 | 0.2 | 0.2 | 2 | — | 2.5 | 1 |
| B1 | | | | | | | | 0.15 | | |
| C1 | | | | | | | 7 | — | | |
| D1 | | | | | | | | 0.15 | | |
| A2 | 0.2 | 0.2 | 0.2 | 0.32 | 0.2 | 0.2 | 2 | — | 2.5 | 1 |
| B2 | | | | | | | | 0.15 | | |
| C2 | | | | | | | 7 | — | | |
| D2 | | | | | | | | 0.15 | | |
| A3 | 0.2 | 0.2 | 0.2 | 0.2 | 1 | 1 | 2 | — | 2.5 | 1 |
| B3 | | | | | | | | 0.15 | | |
| C3 | | | | | | | 7 | — | | |
| D3 | | | | | | | | 0.15 | | |
| A4 | 0.2 | 0.2 | 0.2 | 1 | 1 | 1 | 2 | — | 2.5 | 1 |
| B4 | | | | | | | | 0.15 | | |
| C4 | | | | | | | 7 | — | | |
| D4 | | | | | | | | 0.15 | | |
